# Guide to the construction and use of an adaptive optics two-photon microscope with direct wavefront sensing

**DOI:** 10.1101/2023.01.24.525307

**Authors:** Pantong Yao, Rui Liu, Thomas Broginni, Martin Thunemann, David Kleinfeld

**Affiliations:** Department of Neurosciences, University of California San Diego, La Jolla, CA 92093, USA.; Department of Physics, University of California San Diego, La Jolla, CA 92093, USA.; Department of Bioengineering, Boston University, Boston, MA 02215 USA.; Department of Neurobiology, University of California San Diego, La Jolla, CA 92093, USA.

## Abstract

Two-photon microscopy, combined with appropriate optical labeling, has enabled the study of structure and function throughout nervous systems. This methodology enables, for example, the measurement and tracking of sub-micrometer structures within brain cells, the spatio-temporal mapping of spikes in individual neurons, and the spatio-temporal mapping of transmitter release in individual synapses. Yet the spatial resolution of two-photon microscopy rapidly degrades as imaging is attempted at depths more than a few scattering lengths into tissue, i.e., below the superficial layers that constitute the top 300 to 400 µm of neocortex. To obviate this limitation, we measure the wavefront at the focus of the excitation beam and utilize adaptive optics that alters the incident wavefront to achieve an improved focal volume. We describe the constructions, calibration, and operation of a two-photon microscopy that incorporates adaptive optics to restore diffraction-limited resolution throughout the nearly 900 µm depth of mouse cortex. Our realization utilizes a guide star formed by excitation of red-shifted dye within the blood serum to directly measure the wavefront. We incorporate predominantly commercial optical, optomechanical, mechanical, and electronic components; computer aided design models of the exceptional custom components are supplied. The design is modular and allows for expanded imaging and optical excitation capabilities. We demonstrate our methodology in mouse neocortex by imaging the morphology of somatostatin-expressing neurons at 700 µm beneath the pia, calcium dynamics of layer 5b projection neurons, and glutamate transmission to L4 neurons.

---

A fundamental goal of neuroscience is to understand the complex dynamics of neuronal activity and decipher how the dynamics encodes the information and guide behavior. Two-photon laser scanning microscopy (TPLSM) can image through the mammalian brain with cellular-to-subcellular resolution^1^. With a variety of organic and genetically encoded sensors, TPLSM permits investigators to record neuronal activity in terms of calcium dynamics, neurotransmitter release, even voltage signals^2–5^. Imaging method conserves the spatial information of the recorded neurons for studying their spatial organization such as tonotopic map^6^. The spatial information also be used for aligning the neuronal activity with the gene expression information acquired ex-vivo, allowing the investigator to study the neuronal dynamics in molecularly defined cell type^7^.

The spatial resolution and the fluorescent efficiency of TPLSM degrades with increasing imaging depths because of tissue inhomogeneities and refractive index mismatches. These imperfections in the optical properties of tissue aberrate the wavefront of the excitation beam and lead to an enlarged focus with diminished focal intensity (**Figure 1a,b**). To counteract the optical aberrations, adaptive optics TPLSM (AO-TPLSM) integrates a phase modulator, i.e., a deformable mirror or spatial light modulator, to the conventional TPLSM to compensate the wavefront distortion^8^ (**Figure 1c**). The desired wavefront can be determined by either direct measurement of the distorted wavefront via a wavefront sensor^9–11^ (**Figure 1d**) or by inferring it indirectly from the acquired images^12–16^.

**Figure 1.**
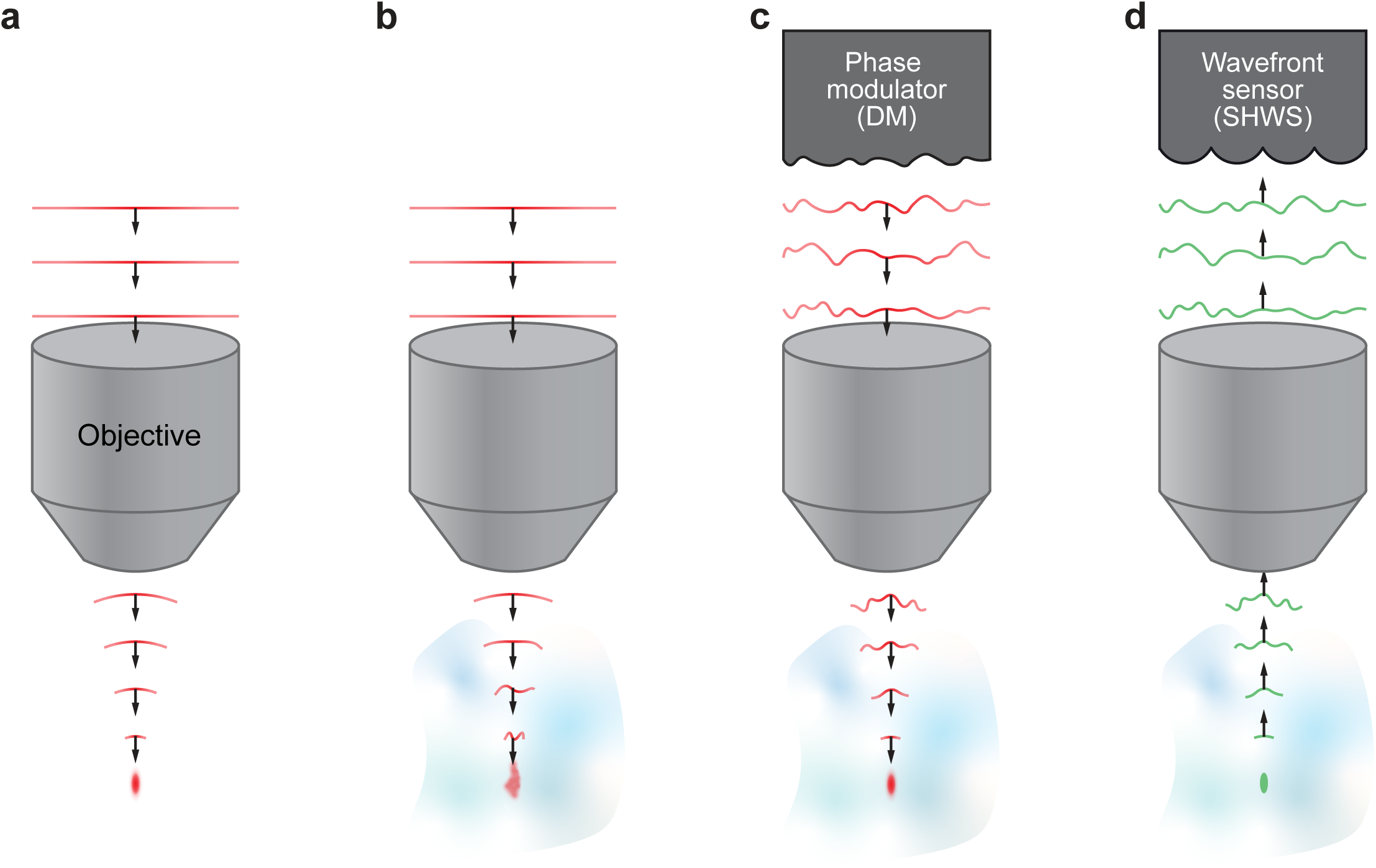
Concepts of adaptive optics and wavefront sensing. **a,** Schematic diagram illustrating the propagating wavefront of the excitation beam in an aberration-free imaging system in which the beam focused as a diffraction limited spot. **b,** Aberrated wavefront as it propagates through the inhomogeneous imaging sample, resulting in an enlarged and distorted focus. **c,** Compensating the aberration by the phase modulator to recover the diffraction limited focus. **d,** Acquiring the desired wavefront for the phase modulator by direct wavefront sensing of the aberrated wavefront that generates by a point source in the sample.

**Figure 2.**
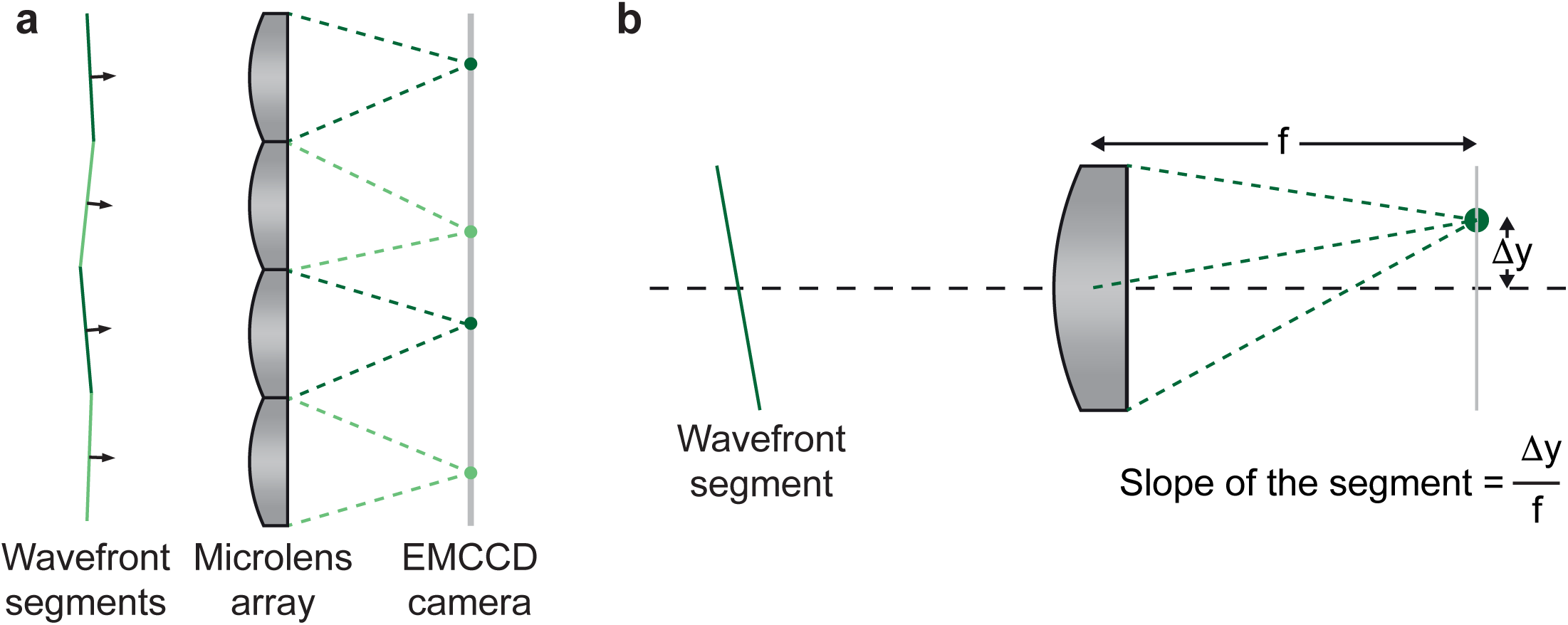
Concepts of Shark-Hartmann wavefront sensing. **a,** Diagram showing that the aberrated wavefront is split into segments by the lens array and focus on the EMCCD respectively. **b**, Diagram demonstrating that the wavefront can be estimated by calculating the slope of each segment based on the deflection of the focal spot. Δy, the distance of the deflected spot from the central axis of the lenslet; f, the focal length of the lenslet.

## Overview of the procedure

Despite its advantages and potential usage in neuroscience research, implementing AO-TPLSM presents challenges. Here we describe a comprehensive solution to make it relatively simple to construct and perform AO-TPLSM imaging. We share the design of our system (**Figure 3,4**) in terms of assembly drawings for all components (**Supplementary Data 1)**. We use commercially available parts as much as possible (**Tables 1** to **3**) and, for the customized parts, detailed drawings and computer aided design (CAD) models are supplied **(Table 4** and **Supplementary Data 2**). We present illustrated step-by-step instructions (**Figure 5**) for the construction (**Procedure I**: steps 1 to 22, and **Supplementary Data 1**), alignment (**Procedure II**: steps 23 to 41), calibration (**Procedures IIIa and IIIb**: steps 42 to 52), and quality control (**Procedure IIIc**: steps 43 and 54). The custom MATLAB codes for calibrating and operating the system as provided (**Supplementary Software**). We provide details of the preparation of the fluorescent molecule, Cy5.5-dextran, that underlies the guide star (**Procedure IV**: steps 55 to 67). Lastly, we provide example demonstrations of the procedure for performing *in vivo* imaging experiments with AO corrections (Procedure V: steps 68 to 74).

**Figure 3.**
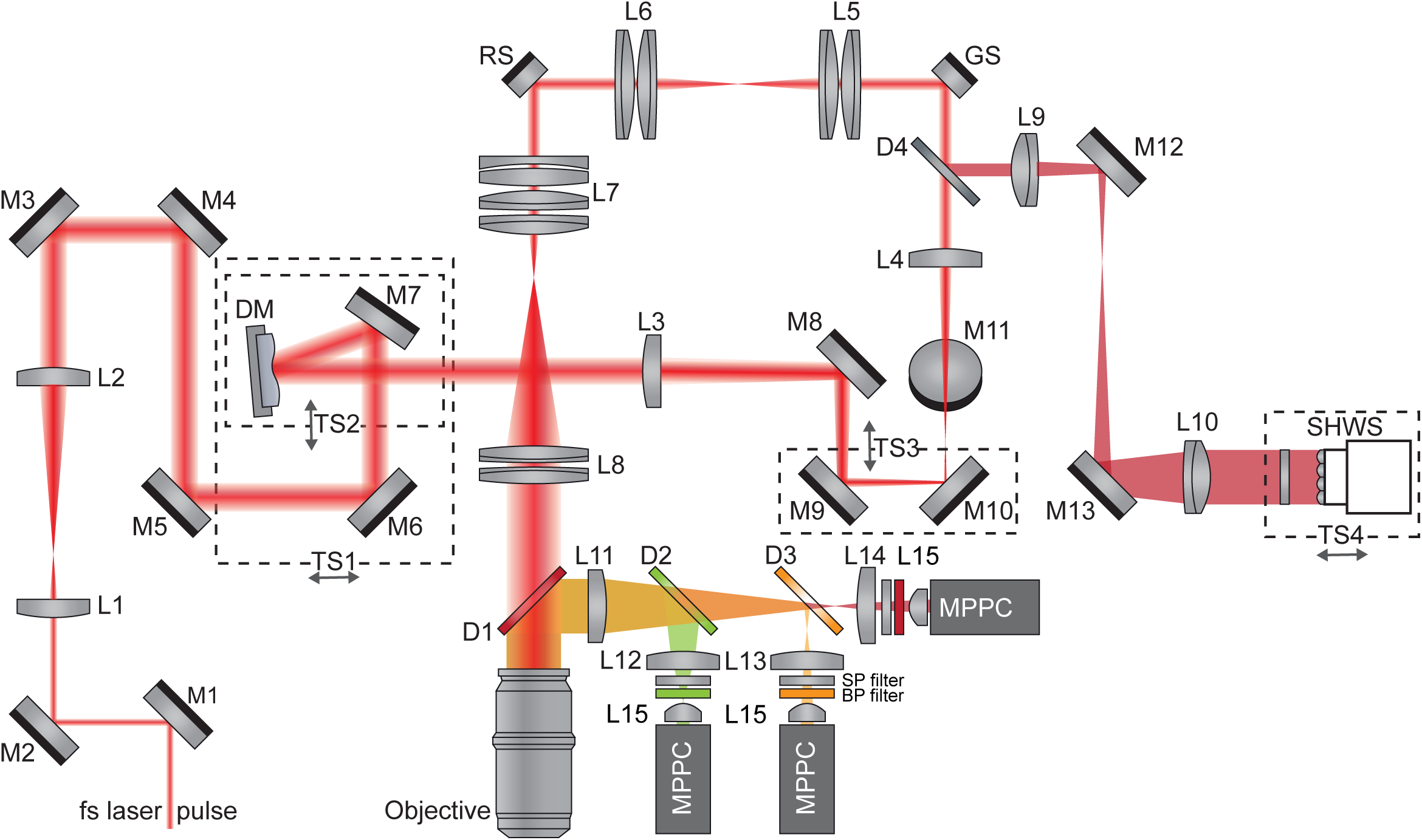
Schematic diagram of AO-TPLSM. Schematic diagram showing optics and beam path of the AO-TPLSM. DM, deformable mirror; SHWS, Shark-Hartmann wavefront sensor; TS, translational stage; M, mirror; L, lens; D, dichroic mirror; SP filter, shortpass filter; BP filter, bandpass filter; GS, galvo scanner; RS, resonant scanner; fs laser, femtosecond laser; MPPC, multi-pixel photon counters.

**Figure 4.**
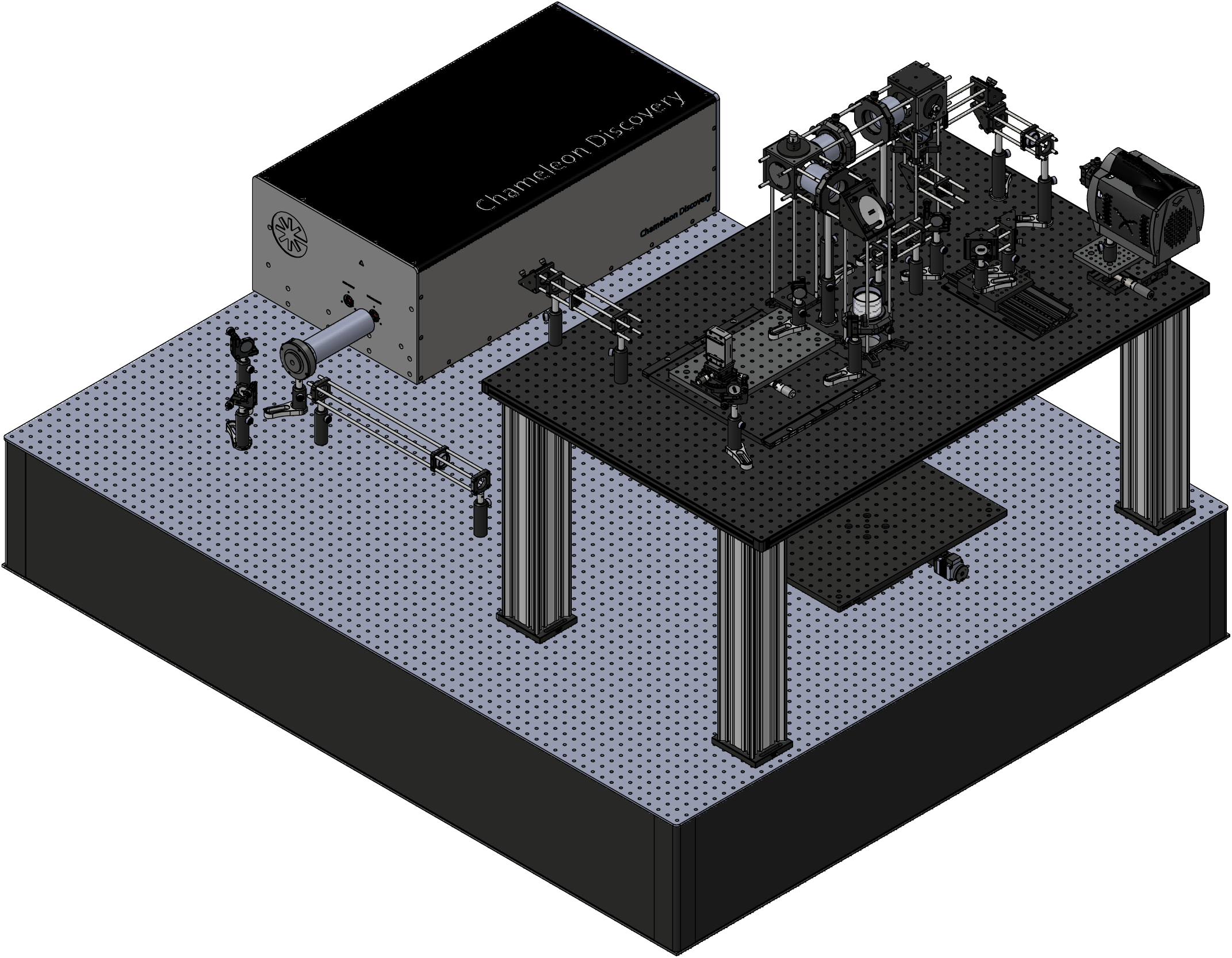
Implementation example of AO-TPLSM. Computer model of AO-TPLSM. Detailed technical drawings are provided in Supplementary Data 1.

**Figure 5.**
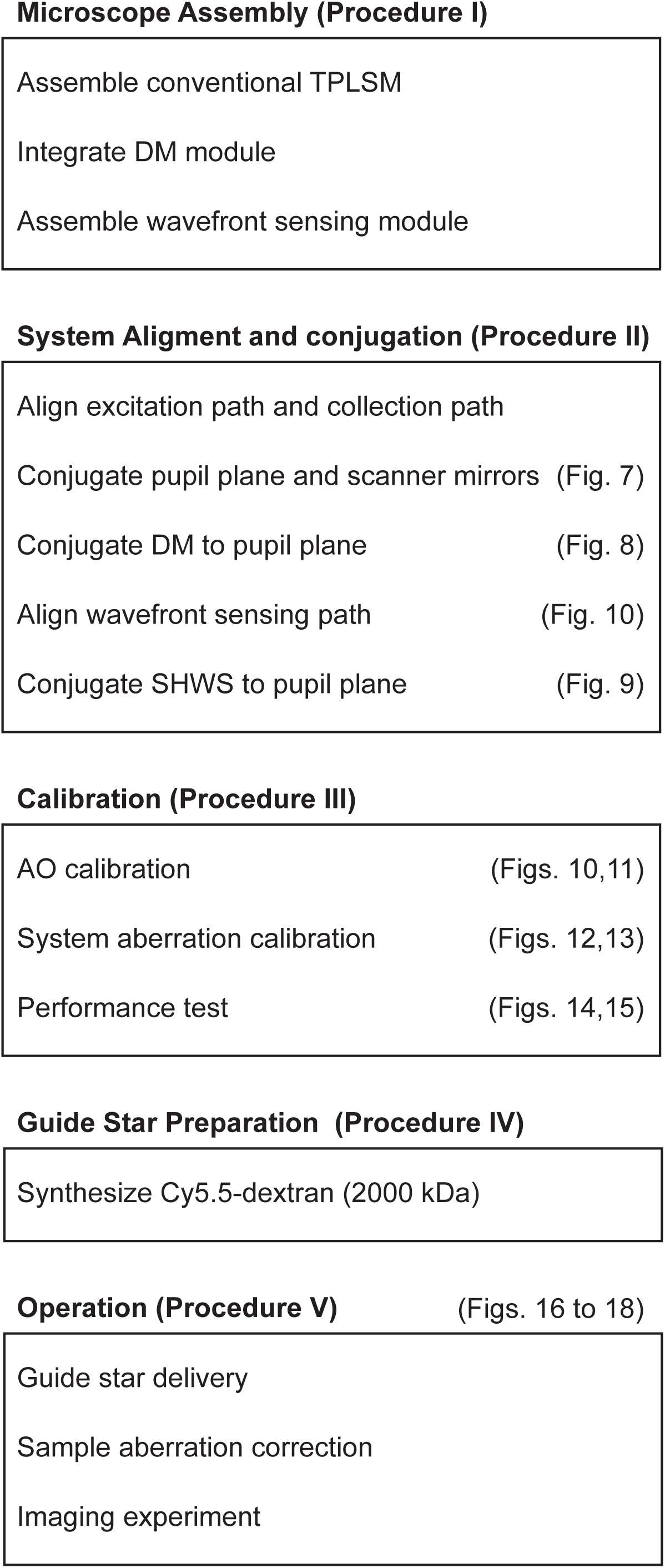
Overview of the procedure. Flowchart illustrating the essential steps for constructing, aligning, and calibrating the AO-TPLSM aw well as showing the main steps for conducting an imaging experiment with AO-TPLSM.

### System Design

Like a conventional TPLSM, the AO-TPLSM contains an excitation path to focus the excitation beam on the preparation, and a collection path that couples the light emitted at the focus to the photon-detectors, e.g., multi-pixel photon counters (MPPCs), or photomultiplier tubes (PMTs). A pair of the scanners that are conjugating to the pupil plane of the objective in the excitation path scans the focus across the preparation during the acquisition of an image. An image frame is constructed from the intensity signal as a function of the scanner angles.

Image quality, including the spatial resolution and signal-to-noise ratio (SNR), is largely determined by the quality of the focus, and is characterized by the point-spread-function (PSF). The goal of AO-TPLSM is to compensate for the aberrated wavefront of the excitation beam to achieve a non-aberrated PSF (**Figure 1c**). The shape of the aberrated wavefront, *W*, can be expressed as a sum of Zernike modes, the polynomials labeled 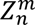 with azimuthal frequency *m* and radial order *n*, where

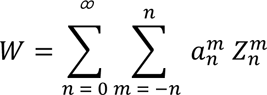

and 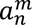 is the coefficient of mode 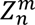. In our implementation of AO-TPLSM, we integrate a deformable mirror (DM) to the excitation path, whose reflective surface is a membrane that can be deformed as sum of Zernike modes 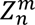, each with magnitude 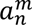, by a group of actuators. This changes the optical path length to introduce a known, corrective distortion to the wavefront. To fully utilize the surface of the DM, the excitation beam is expanded to fill its aperture by a pair of lenses (L1 and L2) between the laser and the DM. The pattern on the DM is imaged to the rear pupil plane of the objective, i.e., the two planes are conjugate (**Figure 6b**). This is achieved by a second pair of lenses (L3 and L4) between the DM and the Y-scanner, that form a four-times focal length (4-f) relay to conjugate the DM with the Y-scanner and demagnify the image of the DM; the scanners are conjugated to the pupil plane as in conventional TPLSM.

**Figure 6.**
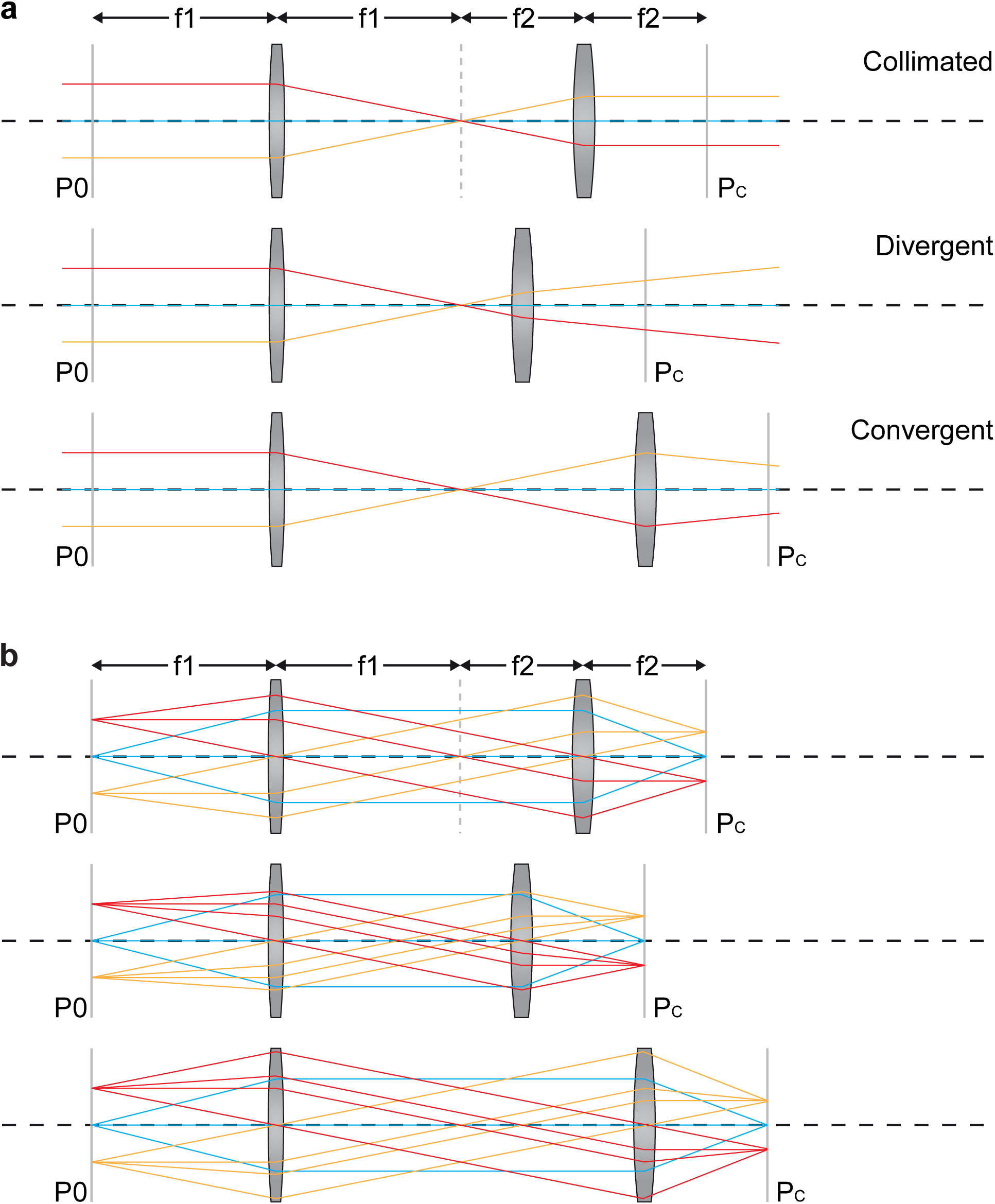
Concepts of collimation and conjugation. **a**, Diagram demonstrating that the divergence of the beam passing through a telescope can be controlled by adjusting the distance between the lens pair. **b,** Diagram illustrating the concepts of conjugation. The conjugate of a given plane P0 is defined as the plane Pc where the points on P0 are imaged. Upper panel shows a 4-f system that the front focal plane of the first lens and the back focal plane of the second lens are conjugate to each other. Middle and lower panel shows that adjusting the distance between the lenses of a telescope does not change the conjugation of the two focal planes. We applied this feature to the design the of relay between the DM and the scanner to orthogonalize the collimation and conjugation.

The wavefront of the descanned guide star signal is measured by direct wavefront sensing (**Figure 1d**) to determine the desired wavefront to apply by the DM to recover the non-aberrated PSF. This is achieved by a Shack-Hartmann wavefront sensor (SHWS), consisting of a microlens array and electron multiplying charge-coupled device (EMCCD) that is located at the focal plane of the microlens array. The local tilt of a subregion of an aberrated wavefront will deflect focused spots on the EMCCD away from the center of the field (**Figure 2a,b**) and is used to estimate the wavefront from the guide star. To separate the guide star signal from the excitation beam, a dichroic mirror (D4) is employed to reflect the guide star signal to the wavefront sensing path. The SHWS also needs to be conjugated to the rear pupil of the objective, like the case for the DM, and another 4-f set of relay lenses (L9 and L10) is introduced to conjugate the microlens array with the Y-scanner.

## Critical task

All told, it is essential to conjugate the X-scanner, Y-scanner, DM, and SHWS to the rear pupil of the objective for successful aberration correction (**Procedure II, Figures 7-9**). However, conjugating these components is not trivial since corrections to the position of one component may break the conjugation between other components. To minimize this difficulty, our system is designed to orthogonalize the conjugation procedure from alignment. For example, the DM is mounted on a translational stage (TS1) with a pair of mirrors (M6 and M7) so that sliding TS1 along the optical axis minimally affects the overall alignment of the excitation beam. A mirror pair on the TS3 are introduced to adjust the optical path between the L3 and L4 so that the collimation of the excitation beam can be adjusted without affecting the conjugation (**Figure 6**). The critical dimensions for conjugation, noted in the assembly drawing, should be carefully followed. Further, we optimized the assembly sequence so that conjugation at the final steps does not affect conjugation of components early in the assembly process.

**Figure 7.**
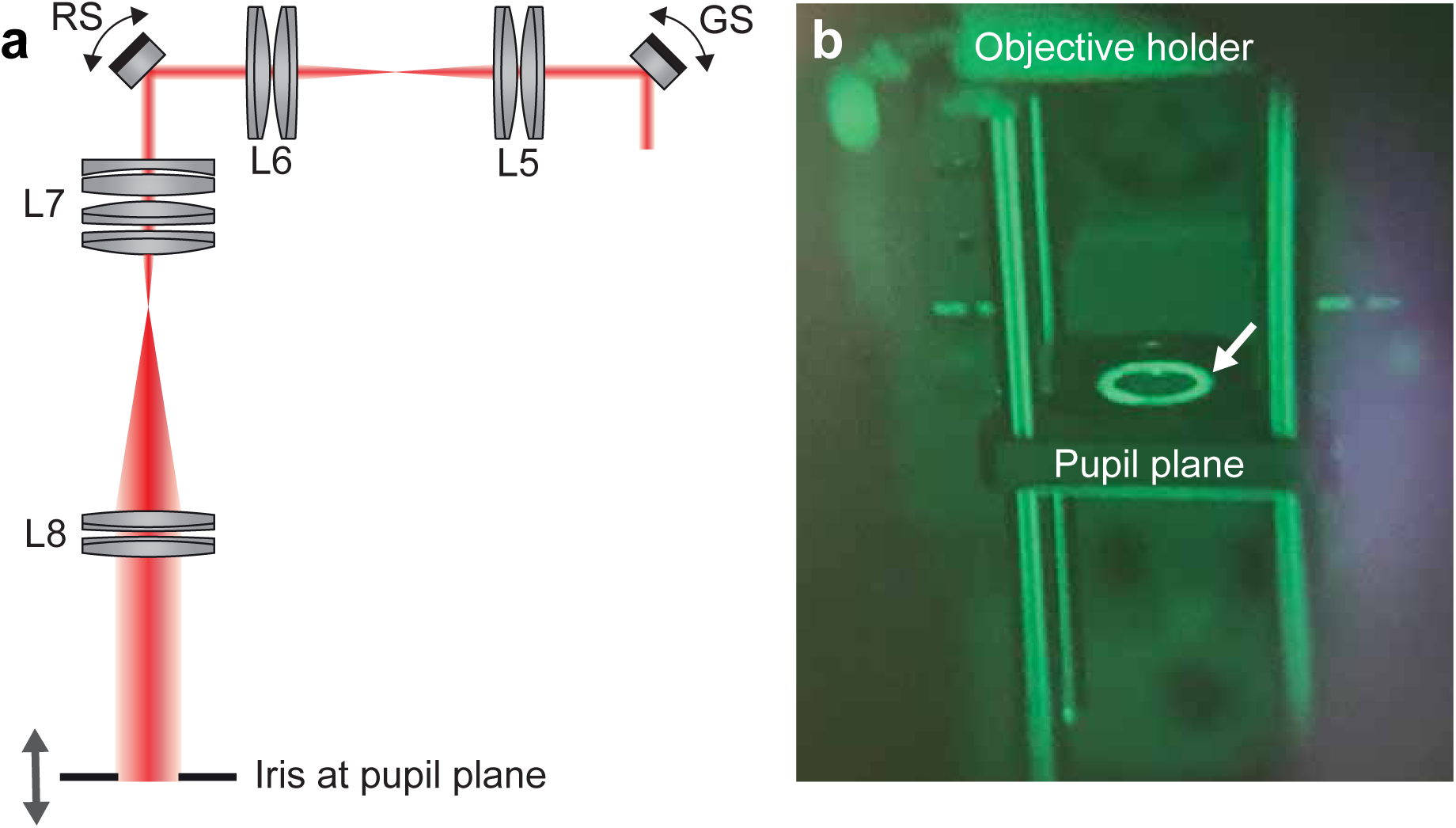
Conjugating the pupil plane with the scanners. **a,** Diagram for finding the objective pupil plane that conjugates to the scanners. **b**, Real item image of the iris at the pupil plane taken through the IR viewer. The beam passing through the iris center (arrow) shows that the excitation beam is well-aligned to the back aperture of the objective. While the scanner is on, little movement of the bright ring should be observed if the iris is conjugate to the scanners.

**Figure 8.**
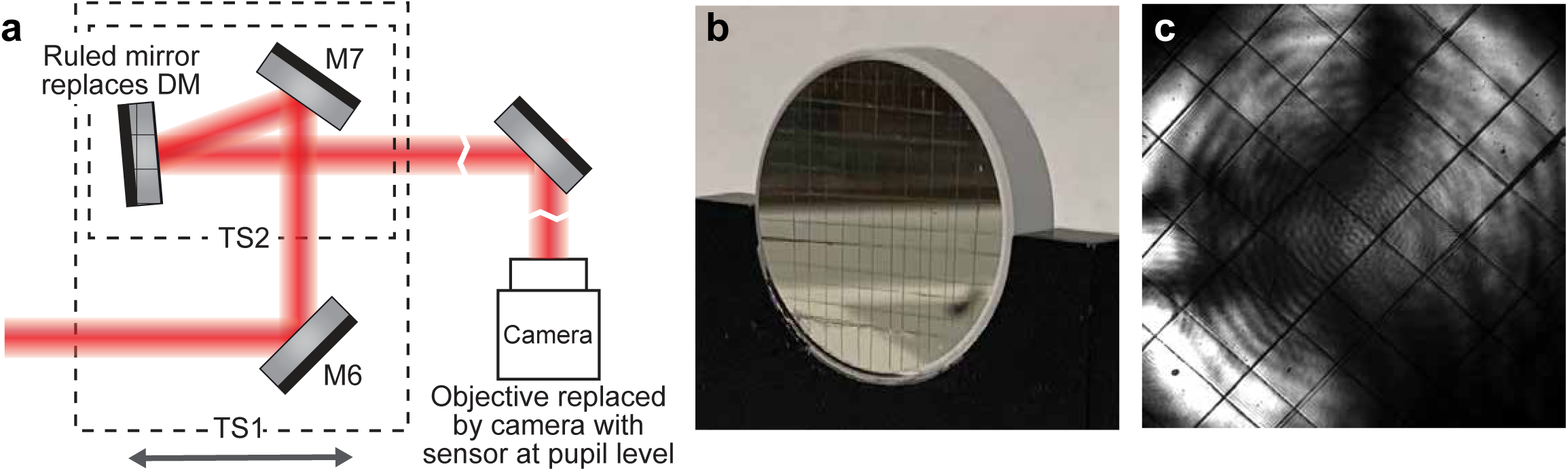
Conjugating the DM to the pupil plane. **a,** Schematics for finding the DM location along the optical axis that conjugates to the pupil plane. The objective is replaced by a camera with its sensor at the pupil plane. The deformable mirror is replaced by a mirror with scratch for the camera to image. **b**, Real item image of the scratched mirror we used to replace the DM. **c**, Image of the scratched mirror taken by the camera at the pupil plane. The sharp grid pattern indicates that the mirror is at the conjugates plane of the pupil.

**Figure 9.**
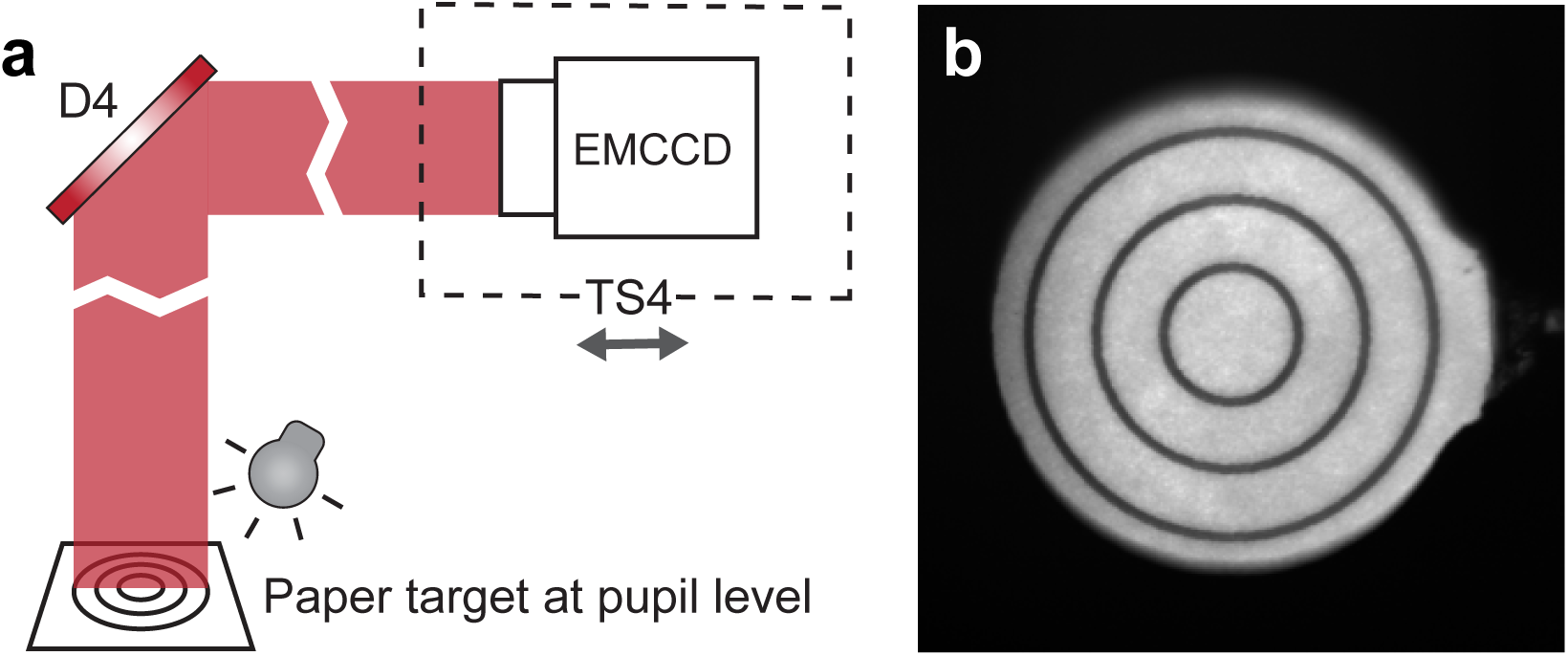
Conjugating SHWS to the pupil plane. **a**, Schematics showing the setup for finding the plane in the wavefront sensing path that conjugates to the pupil. The laser shutter was closed in this step. Another light source (e.g., flashlight) is needed to illuminate the paper target at the pupil plane. **b**, Sharp image of the target, indicating that the EMCCD sensor is conjugated to the pupil plane.

### Operational Logic

Besides aberrations that caused by brain tissue, the imaging system per se aberrates the excitation beams because of imperfections of the optics in the microscope and the aberrations induced by the cover glass of the cranial window. These are referred as system aberrations. To achieve the best image quality, as well as to maximize the imaging depth, both sample aberrations and system aberrations need to be corrected.

For the system aberration, an indirect wavefront sensing method is applied to search for the best pattern on the DM that recovers the PSF in the aberration-free sample (**Procedure IIIb**). The pattern on the DM can be considered as a sum of individual Zernike modes. We developed a gradient-descent algorithm^11^ (**Supplementary Software**) to optimize the coefficient for each Zernike mode by maximizing the emission intensity of a fluorescent solution, under a cover glass (**Figure 12**). The procedure starts with a flat pattern displayed on the DM. In each iteration, images are obtained with individual Zernike added to or subtract from the DM pattern that was obtained from prior iteration. The intensity gradient with respect to each Zernike coefficient is determined as the difference between the average intensity of the images taken with the corresponding Zernike modes added versus subtracted. At the end of each iteration, all Zernike coefficients are updated based on the result of the gradient-descent algorithm. After several rounds of iteration, the coefficients of all Zernike modes are seen to stabilize (**Figure 13**). This leads to a diffraction-limited PSF in the aberration-free sample (**Figure 14**).

To determine the wavefront pattern, or phase map, for correcting the sample aberration, we apply the direct wavefront sensing method and measure the wavefront of the descanned guide star via the SHWS. Conceptually, the guide star refers to an ideal point source of light in the sample whose wavefront will be aberrated as it propagates through the tissue to the rear pupil (**Figure 1d**). On basis of the reversibility of optical propagation, applying the same aberrated wavefront at the pupil generates a diffraction-limited focus in the tissue (**Figure 1c**). In practice, as a two-photon excited fluorescence naturally produces a confined light source, we used the emitted light of two-photon excited cyanine 5.5-conjugated 2000kDa dextran (Cy5.5–dextran) in the microvessels as the guide star^11, 17^. Cy5.5–dextran is a red-shifted dye that is bright, inexpensive to produce (**Procedure IV**), apparently nontoxic, and readily delivered to the vessels by either retro-orbital injection^18^ or tail vein injection. Unlike directly injecting a dye into a region of the brain^9^, vascular circulation constantly replenishes the guide star and makes it compatible with transcranial windows^19^ as well as cranial windows^20^. for chronic imaging.

Note that the guide star signal is also aberrated by the optics of the microscope, including the cover glass and the lenses and mirrors in the last part of the excitation path and wavefront sensing path, which needs to be nulled by a reference pattern of the spot taken from an aberration-free sample, e.g., a fluorescent solution under a cover glass (**Figure 10c**).

**Figure 10.**
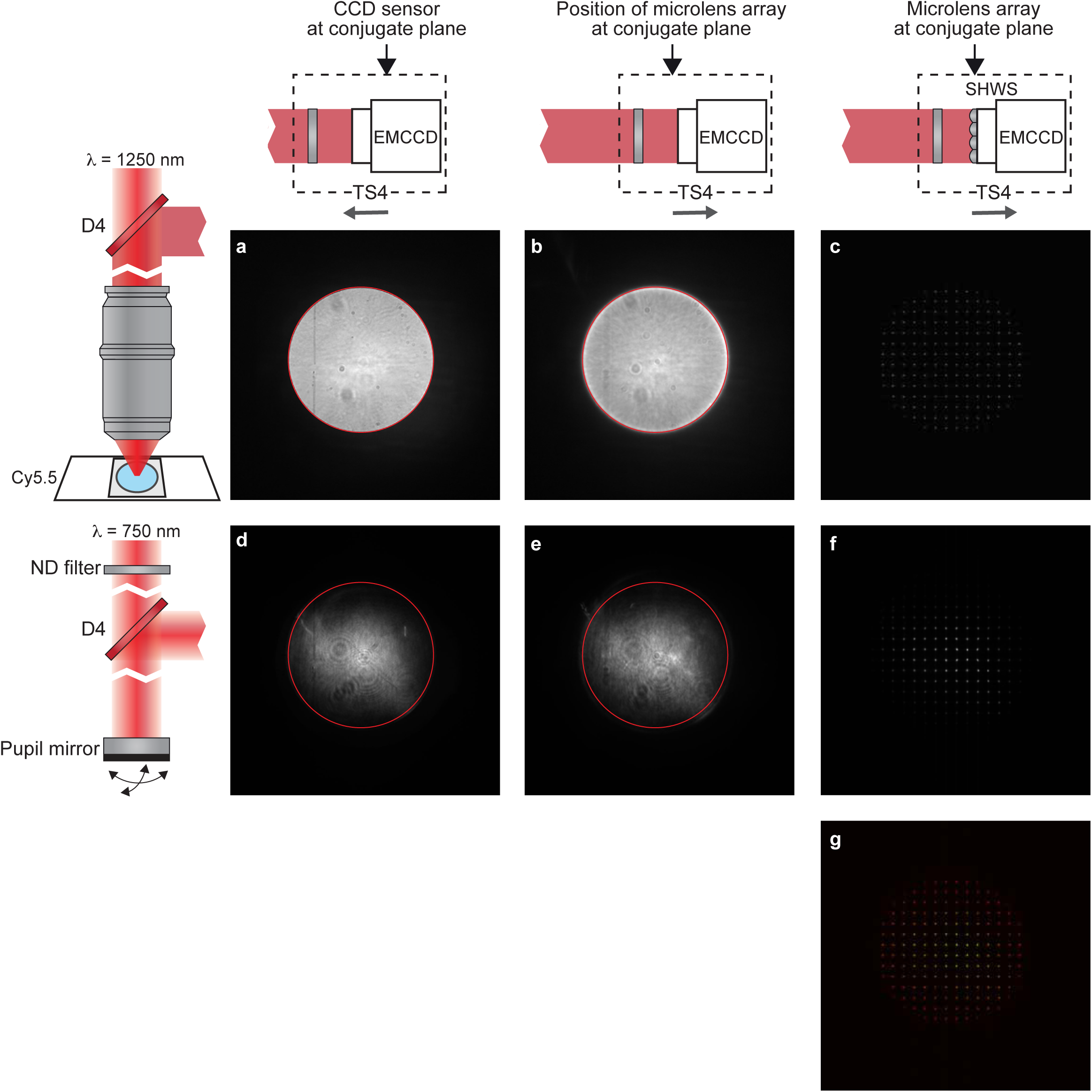
AO Calibration preparation. **a** to **f**, Images obtained on the EMCCD before the AO Calibration. Left column, diagram showing the light source for the EMCCD images. **a** to **c** was taken with the Cy5.5 fluorescence, **d** to **f** used the femtosecond laser beam reflected by a mirror at the pupil plane. Upper row, diagram showing the position of the EMCCD along the optical axis. **a** and **d** were taken by the EMCCD at the conjugate plane. **c** and **f** were taken by the SHWS of which the microlens array is at the conjugate plane. **b** and **e** were taken with EMCCD at the same position as **c** and **f** but no microlens array was placed on the light path. The edge of bright disk in **a** was indicated by a red circle in image **a**, **b**, **d,** and **e**, showing that the bright disks in the four images are concentric. **g**, Merge of **c** and **f**. It is critical to the AO calibration that **a**, **b**, **d,** and **e** are concentric, and that **c** and **f** are overlapped.

To compute the DM pattern based on the SHWS images from the guide star, it is critical to establish the relation between the DM pattern in terms of its Zernike modes to the SHWS in term of the deflection of the spots from each lens in the microarray. This is achieved by AO calibration (**Procedure IIIa**, **Figure 11**), in which we place a mirror at the pupil level to map the DM to the SHWS and record the corresponding pattern of the deflected spots on SHWS when each of the individual Zernike modes is displayed by the DM.

**Figure 11.**
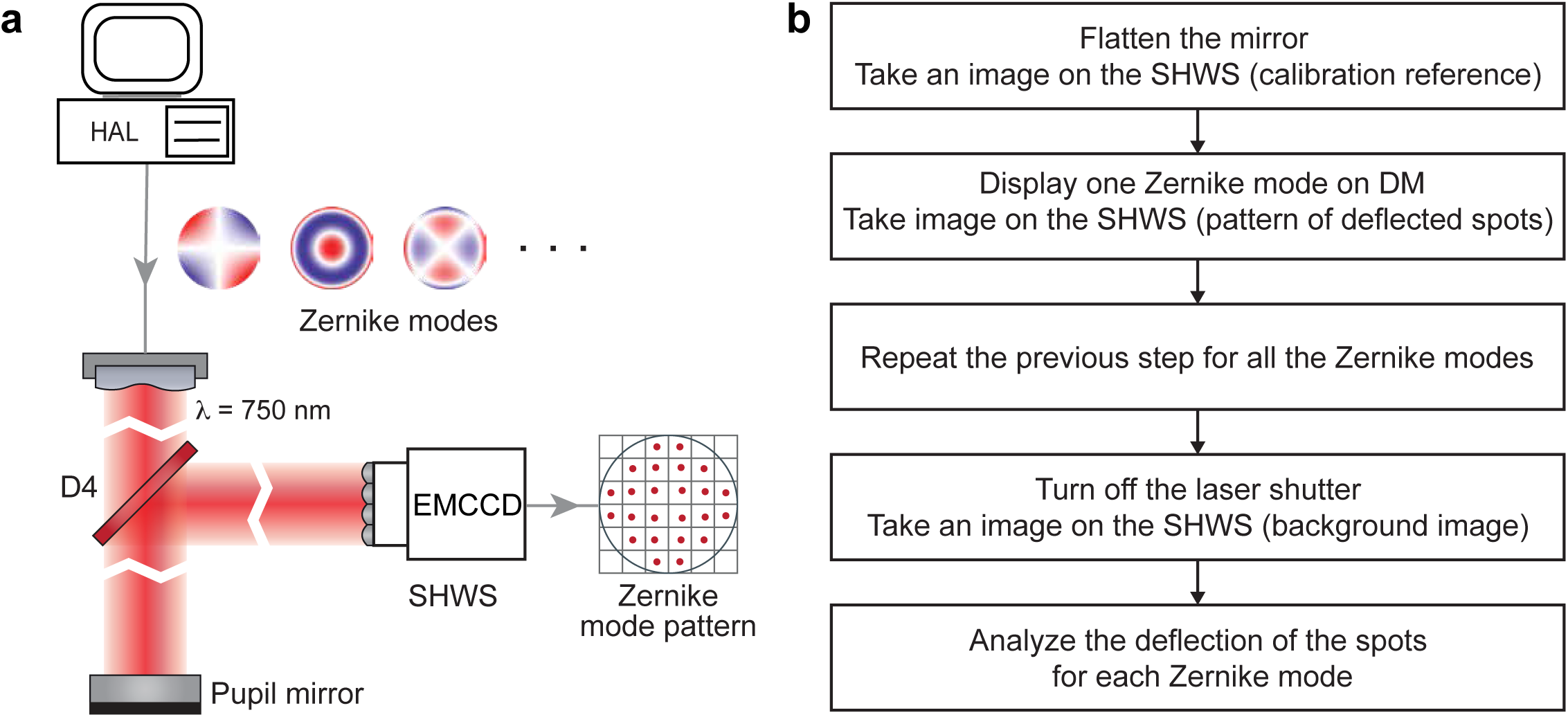
AO Calibration. **a,** Schematics illustrating the AO calibration. **b,** Flowchart of the AO calibration procedure. It corresponds to the MATLAB codes “DM_SHWS_calibration_Ctr.m” and “DM_SHWS_calibration.m”.

**Figure 12.**
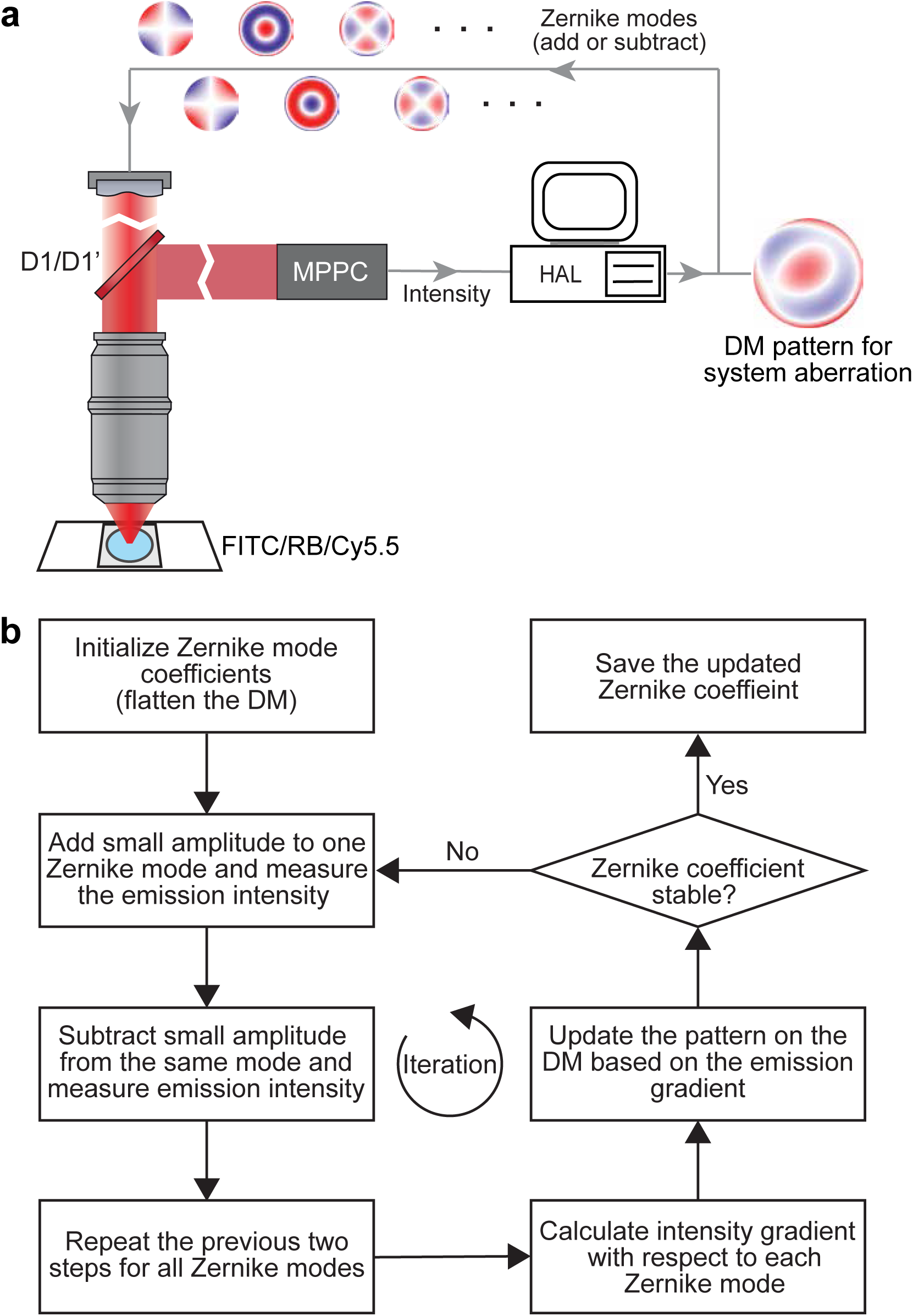
Configuration for System aberration calibration. **a,** Schematics illustrating the configuration for system aberration calibration. **b,** Flowchart of the system aberration calibration procedure. It corresponds to the MATLAB codes “sensorlessWF_sys_abr_descend.m”.

**Figure 13.**
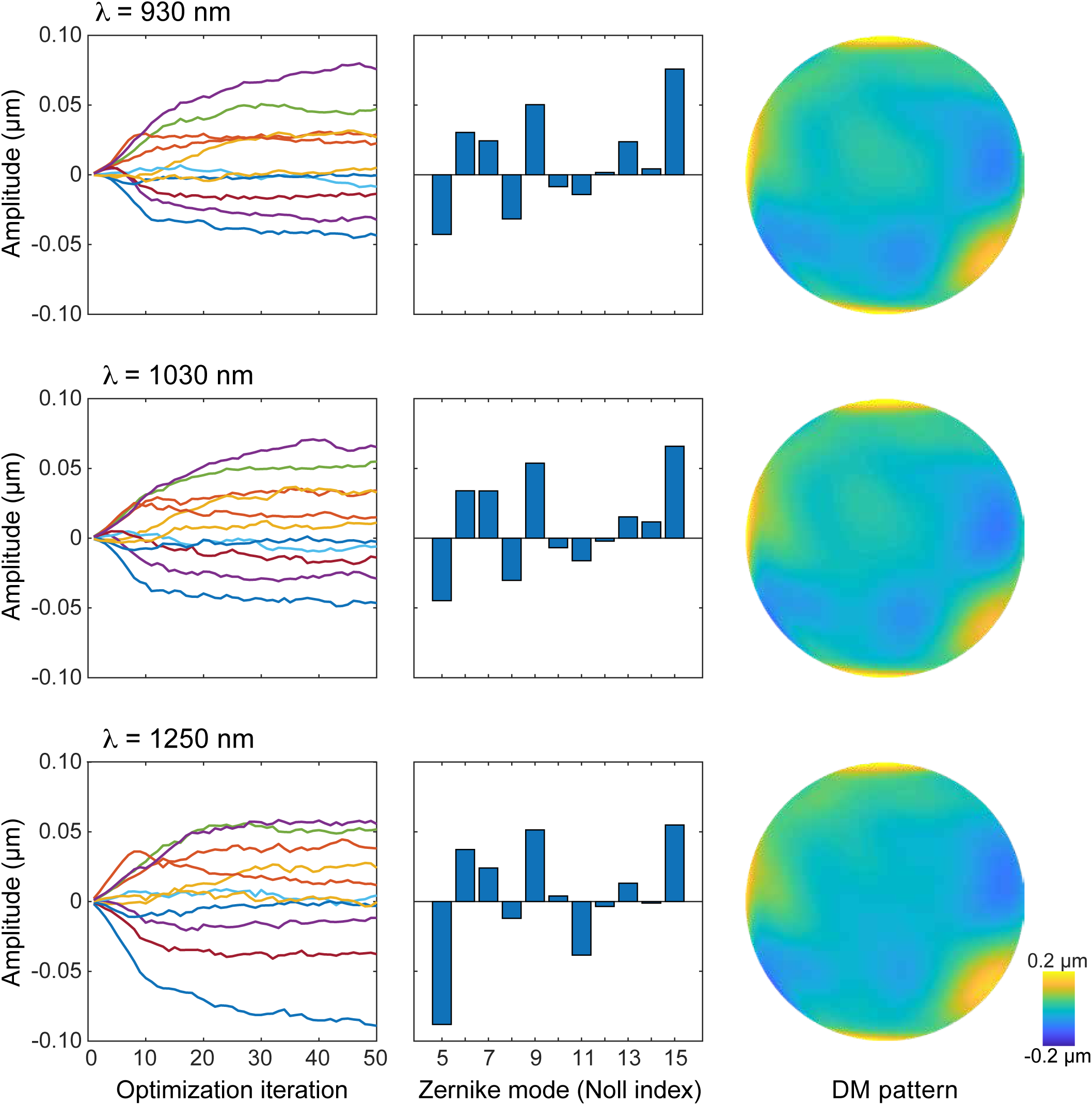
Results for system aberration calibration. Representative outcomes of system aberration calibration for the excitation beam in 930 nm (upper row), 1030 nm (middle row), and 1250 nm (bottom row). Left column, showing the changes of Zernike coefficients as a function of optimization cycles. Middle column, Zernike coefficients after 50 optimization cycles. Right column, wavefront phase map on the DM that compensates for the system aberration.

**Figure 14.**
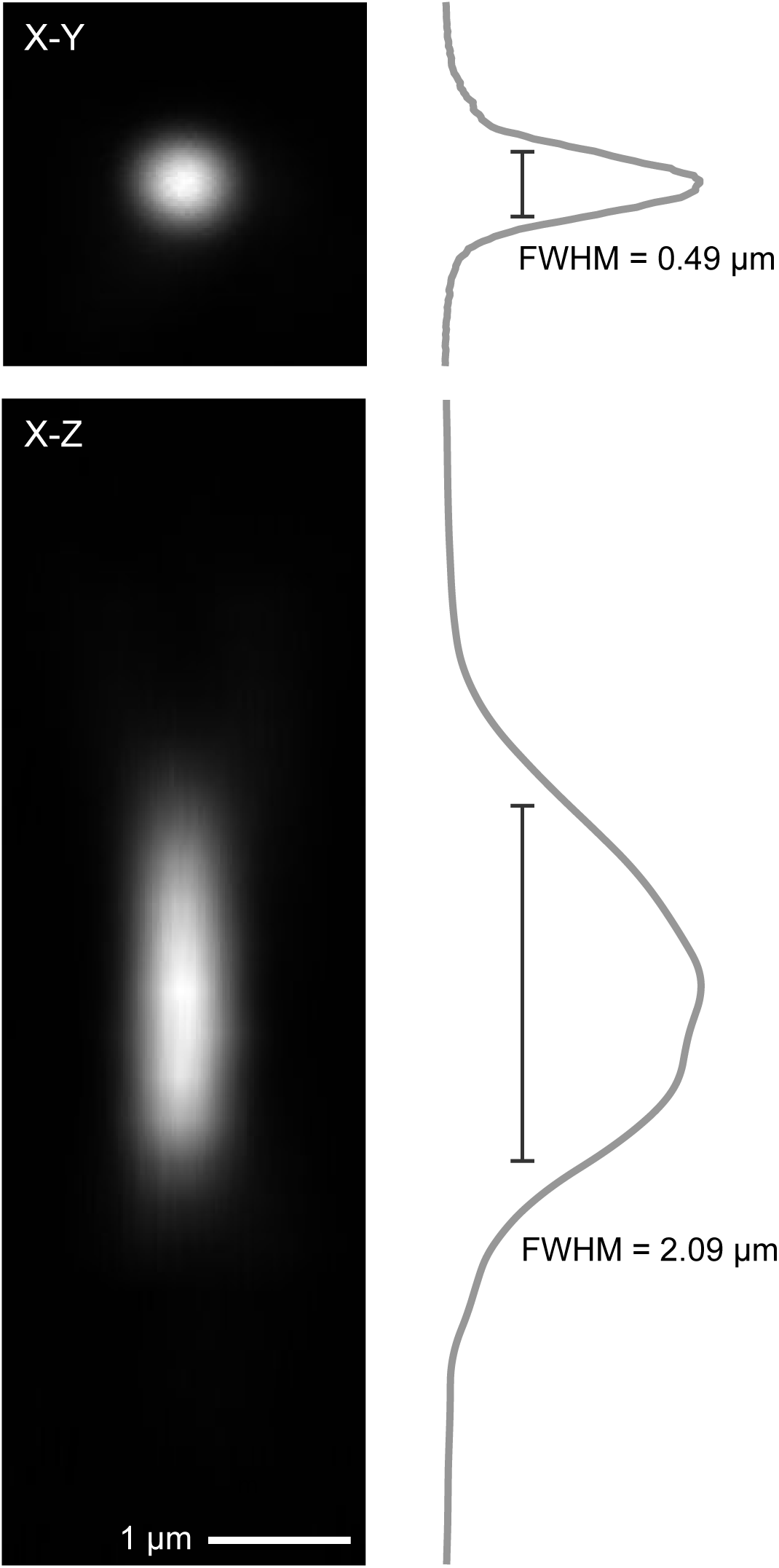
Point spread function after system aberration correction. Point spread function measured from a 200-nm diameter fluorescent bead after the correcting the system aberration. Scale bar, 1 μm. Upper row, image of the X-Y plane with the signal profile showing the lateral resolution. Bottom row, image of the X-Z plane with the signal profile, showing the axial resolution.

## Critical

We encourage the user to calibrate the system regularly, e.g., twice a year for the best performance, including the AO calibration (**Procedure IIIa**), system aberration calibration (**Procedure IIIb**). We also recommend that the user verify the imaging system by correcting the aberration for a fiber test sample after the calibration to ensure the system operates properly (**Procedure IIIc**).

The sample aberration, as it varies regions by region, should be corrected for every field of view during experiment (**Procedure V**).The region in which a single correction achieves substantial improvement in image quality is defined as the isoplanatic patches. The isoplanatic patches size of the mouse brain is about hundreds of micrometers^9, 11, 21–23^. In practice, to achieve the best imaging quality, we recommend the user to perform sample aberration correction for each 100 μm x 100 μm, or 50 μm x 50 μm, field of view and apply the resulting pattern within ±50 μm axially.

### Applications

With the capability of correcting the aberration of a given sample, AO-TPLSM could be widely applied to improve the image resolution and SNR during *in vivo* imaging. We previously demonstrated near-diffraction-limited images using direct wavefront sensing from descanned guide star in the brain microvessels in awake behaving mice. We resolved soma and fine structures, such as dendrites and individual spines, up to 850 μm below the pia^11^. Together with the expression of genetically encoded sensors, this approach has been used in the study of cortical deep layers, including the recording of the calcium transients in spines of layer 5b basal dendrites^11^ and imaging the glutamate signal from presynaptic boutons and postsynaptic spines during tactile sensing^3^. Besides the cortical region, AO-TPLSM has been employed in correcting the optical aberrations of the mouse eye for noninvasive retinal imaging at subcellular resolution^24^.

Endoscopic GRIN lens extends the optical access to regions that deeply buried in the brain^7, 25, 26^. However, due to the intrinsic aberrations of the GRIN lens, the imaging resolution decays dramatically radially and axially, resulting in a small field of view and narrow depth of field. AO-TPLSM with direct wavefront sensing can also counteract the aberration of the GRIN lens by treating it as sample aberration but with 30 μm x 30 μm isoplanatic patches^27^.

In addition to the improvement of imaging quality, the DM in the AO-TPLSM also serves as a convenient tool for manipulate the focus spot of the microscope. For example, the third Zernike mode of the DM (defocus, Noll Index 4) moves the focus along the optical axis without introducing extra aberration. With the fast settle time of the DM (∼ 0.5 ms), defocus can be used for multiplane imaging. The first two Zernike modes of the DM (Tip, and tilt, Noll Index 2 and 3) moves the focus along the X, Y axis, which can be potentially used for online motion correction.

### Alternative methods

The essential task for adaptive optics is the aberration measurement that finds the phase map for the DM to play for compensating the optical aberration (**Figure 1c,d**). Besides the direct wavefront sensing method we detailed in the protocol, indirect sensing method has also been widely implemented in TPLSM^14, 21^ and even three-photon microscopy^15, 16^. Instead of guide star signal, the indirect method inferred the desired wavefront from the acquired images. Since the aberrated wavefront usually leads to degraded image, one can find the desired wavefront by tuning it based on certain metrics of the image. It is easier to implement as it does not have the wavefront sensing light path and as no guide star needs to be delivered before the experiment. However, the disadvantage of the indirect approach is that it is takes more time than sensor-based direct methods because they typically require a collection of many images and need multiple iterations. On the other hand, the direct method we introduced in this protocol has better signal-to-noise ratio (SNR) and imaging depth comparing to all the indirect methods that were implemented in TPLSM^11^.

### Limitations

Limited by the scattering of the tissue, the maximum image depth of AO-TPLSM is about 800 ∼ 900 μm under the pia, where the guide star pattern on the EMCCD degrades severely for computing the wavefront. The emission of Cy5.5 is peak at 703 nm; therefore, the wavefront distortion of the guide star differs from the one of the excitation beams. A dye with longer emission is desired for matching with the excitation wavelength and for better scattering tolerance for deep wavefront sensing. Reducing the expansion ratio of the descanned guide star, can potentially increase the signal to noise for the image on EMCCD by concentrating the light intensity on the microlens array. However, this will reduce the number of the focus spots for wavefront sensing; therefore, reducing the precision of the wavefront estimation.

We designed an entire system from scratch rather than modify an existing plan or commercial system for ease of flexibility and because the bulk of the cost is AO components and tunable femtosecond laser.

### Expertise needed to implement the protocol

We put our effort in the design of AO-TPLSM and in the protocol detailed here to make it the easy to setup, maintenance, and use for laboratories with limited previous imaging expertise. Despite this, basic knowledge in optics, electronics and programming is required for the primary user, who is responsive for assembling, aligning, and calibrating the system. Prior experience in setting up imaging systems can largely facilitate the project but is not required. The users that are involved in the *in vivo* imaging experiment should have the ability to perform the craniotomy (or thin-skull surgery) and operate basic imaging software.

## MATERIALS

### Hardware

- Optical table (at least 4’ by 6’ in extent)
- Tunable femtosecond laser (e.g., Coherent, Chameleon Discovery)

#### Critical

The wavelength of the laser must cover 1250 nm (at least 1200 nm) with reasonable output power (at least 1000 mW) for wavefront sensing with Cy5.5). The imaging depth can be achieved by AO-TPLSM is largely determined by the maximum depth of wavefront sensing, which is constrained by the laser power. The power efficiency of the AO-TPLSM at 1250 nm is about 10 - 15 % (power measured at the laser/ power measured at the objective output). And previous study reported that no photodamages was observed when using average laser powers of approximately 250 mW for *in vivo* imaging. Therefore, ∼2000 mW laser output at 1250 nm should be ideal to generate up to 200 to 300 mW of excitation beam power for wavefront sensing without photodamaging.

- Optical components including the microlens array (Edmund, #64-483) are listed in **Table 1**.
- Optical electronics and opto-electronics including the DM (ALPAO, DM97-15) and EMCCD (Andor, DU-888U3-CS0-BV) are listed in **Table 2**.

**Table 1.**
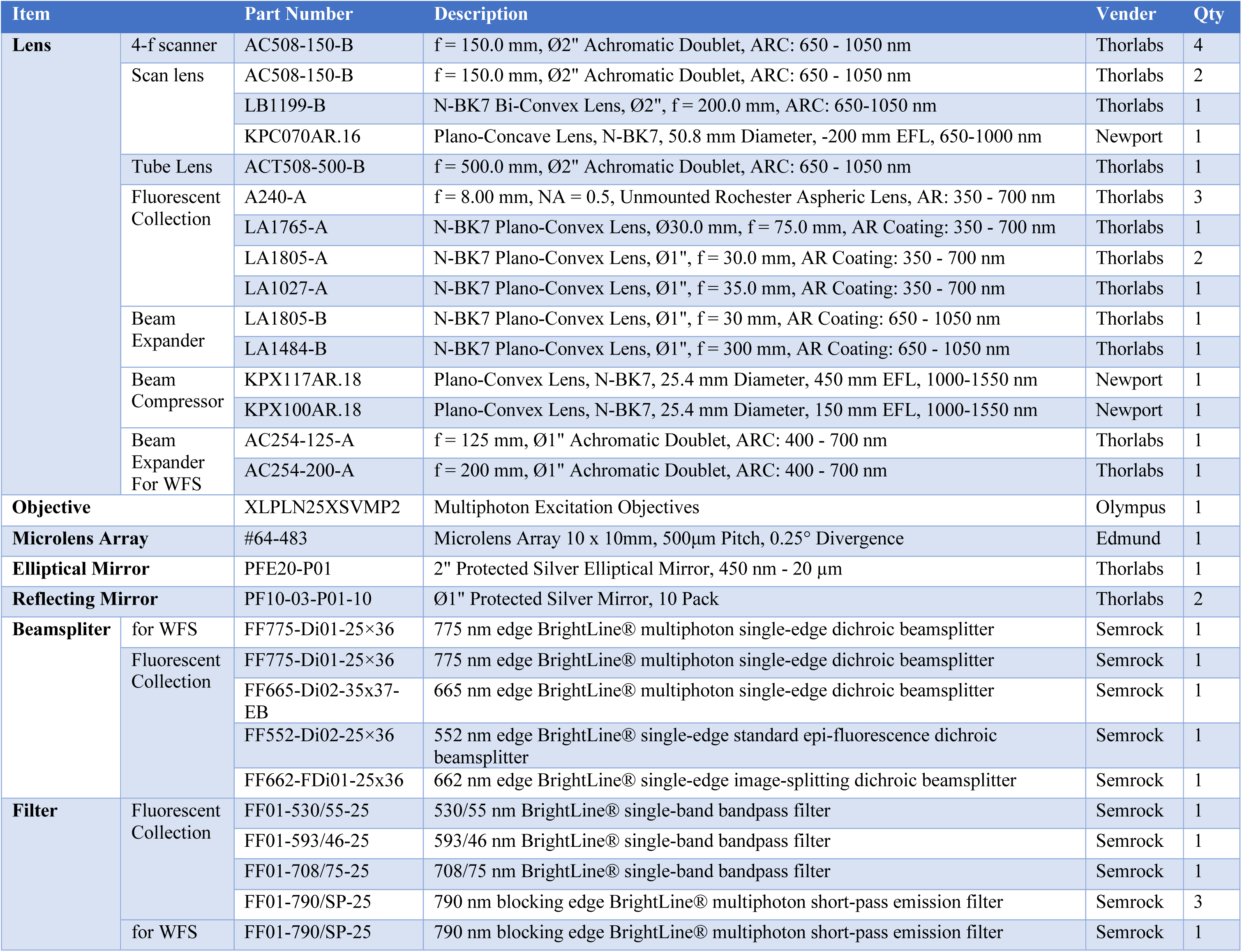
Optical components for AO-TPLSM.

**Table 2.**
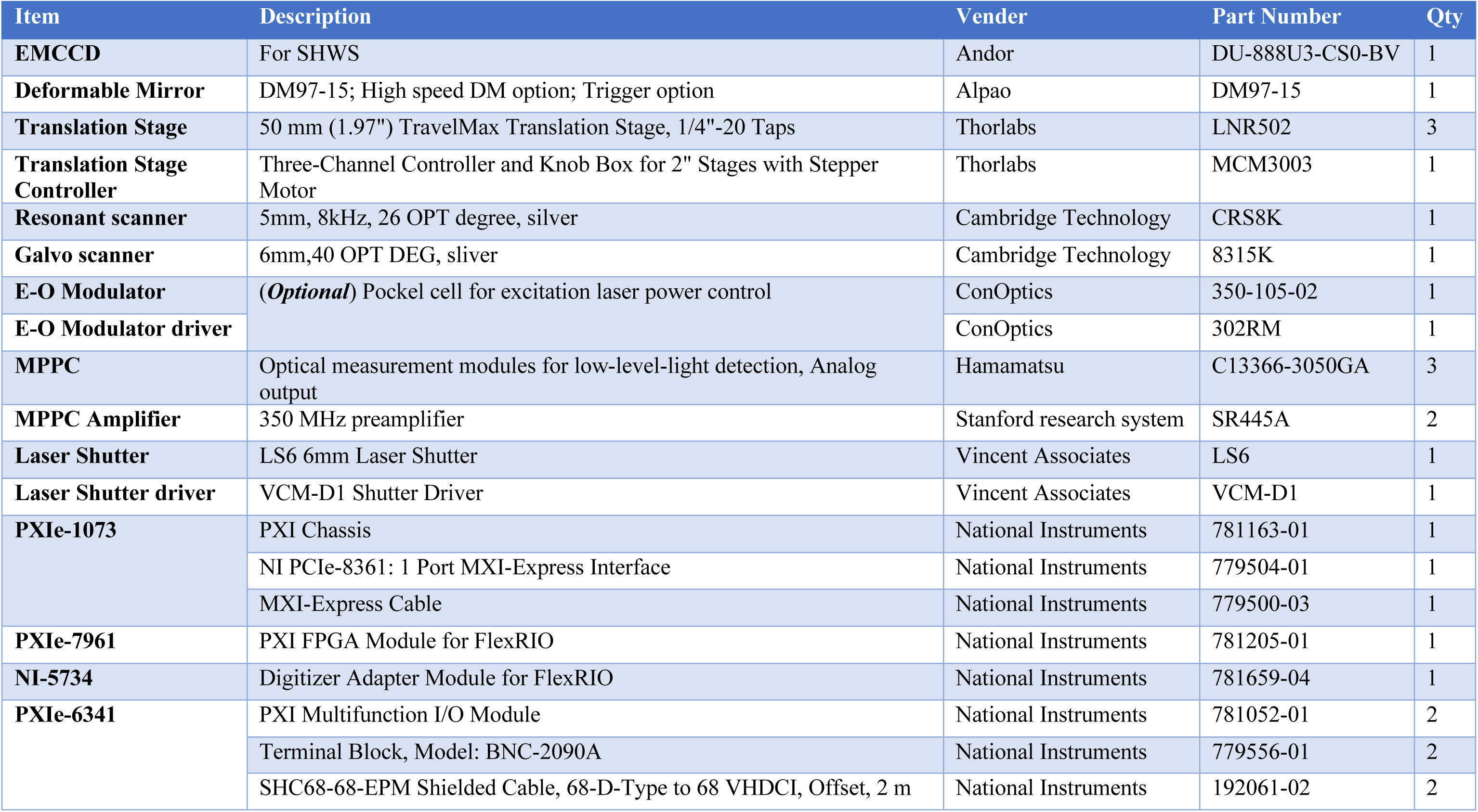
Electronics and opto-electronics for AO-TPLSM.

#### Critical

The trigger-option of the deformable mirror controller is required for synchronization between the DM and the Scanimage. We used High-speed option of the DM97-15 in our implementation, which has quick settling time but less stroke (dynamic range) as trade-off. High stability option should be more suitable for general user.

- Commercial opto-mechanics are listed in **Table 3**.

**Table 3.**
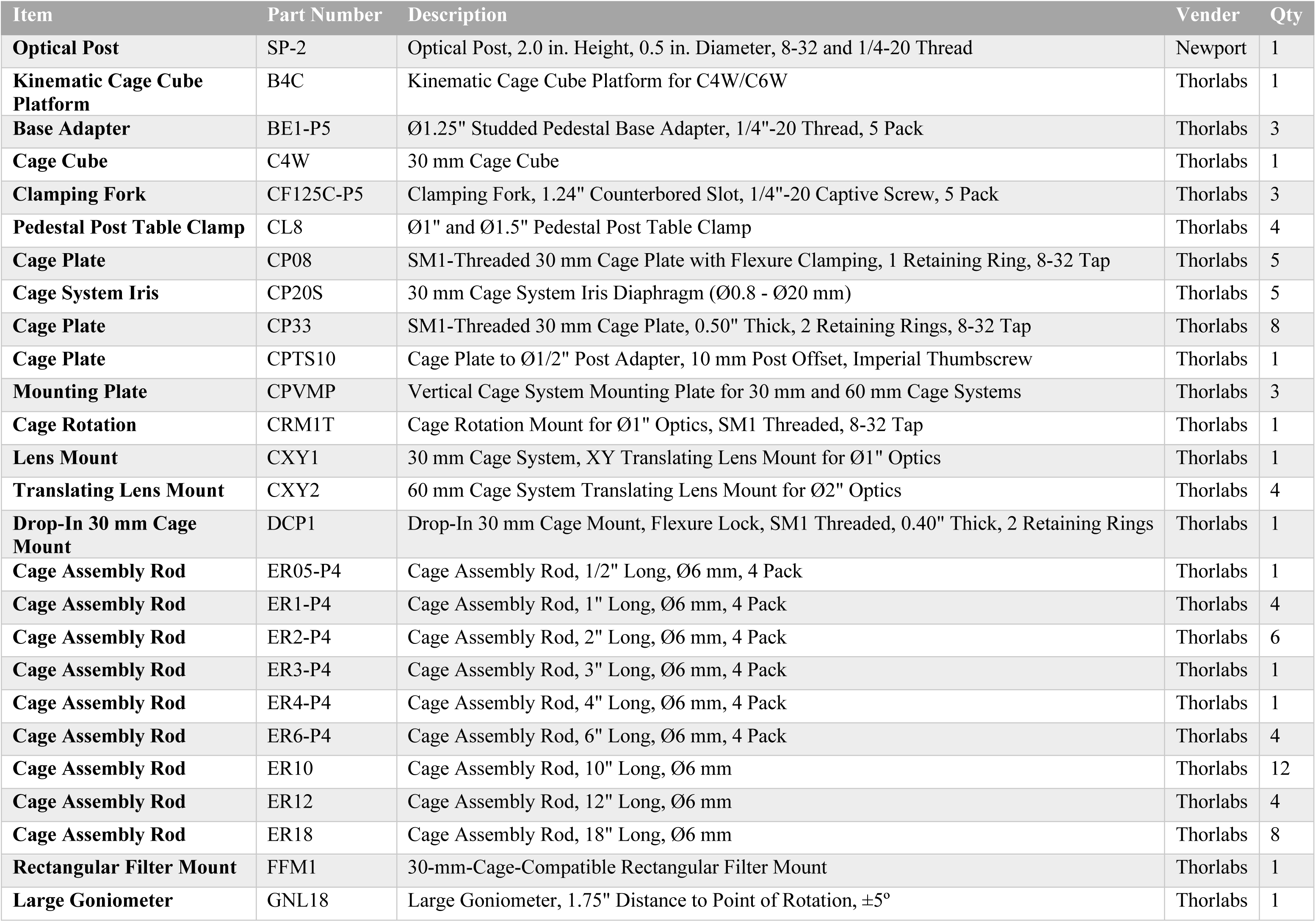
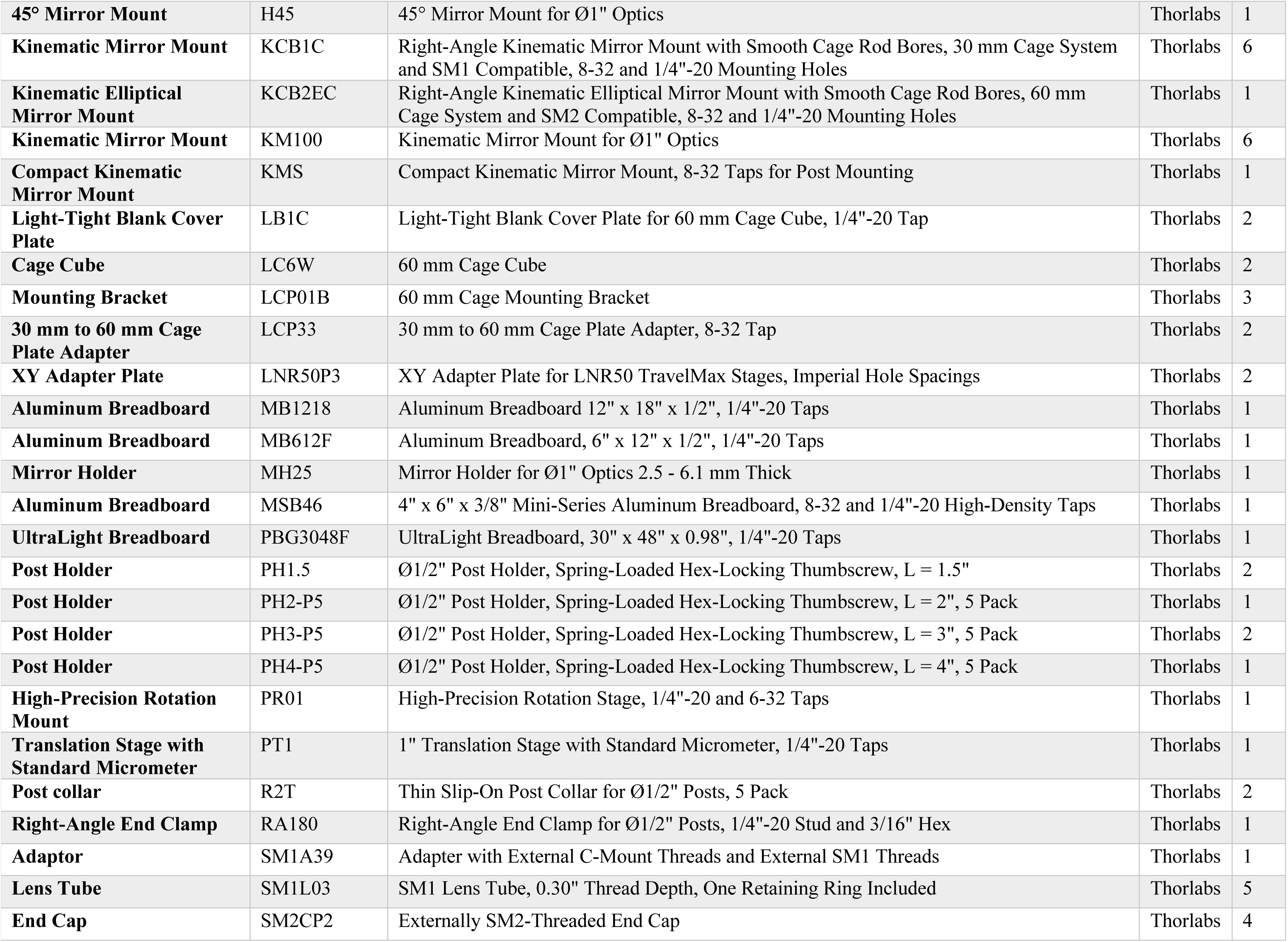
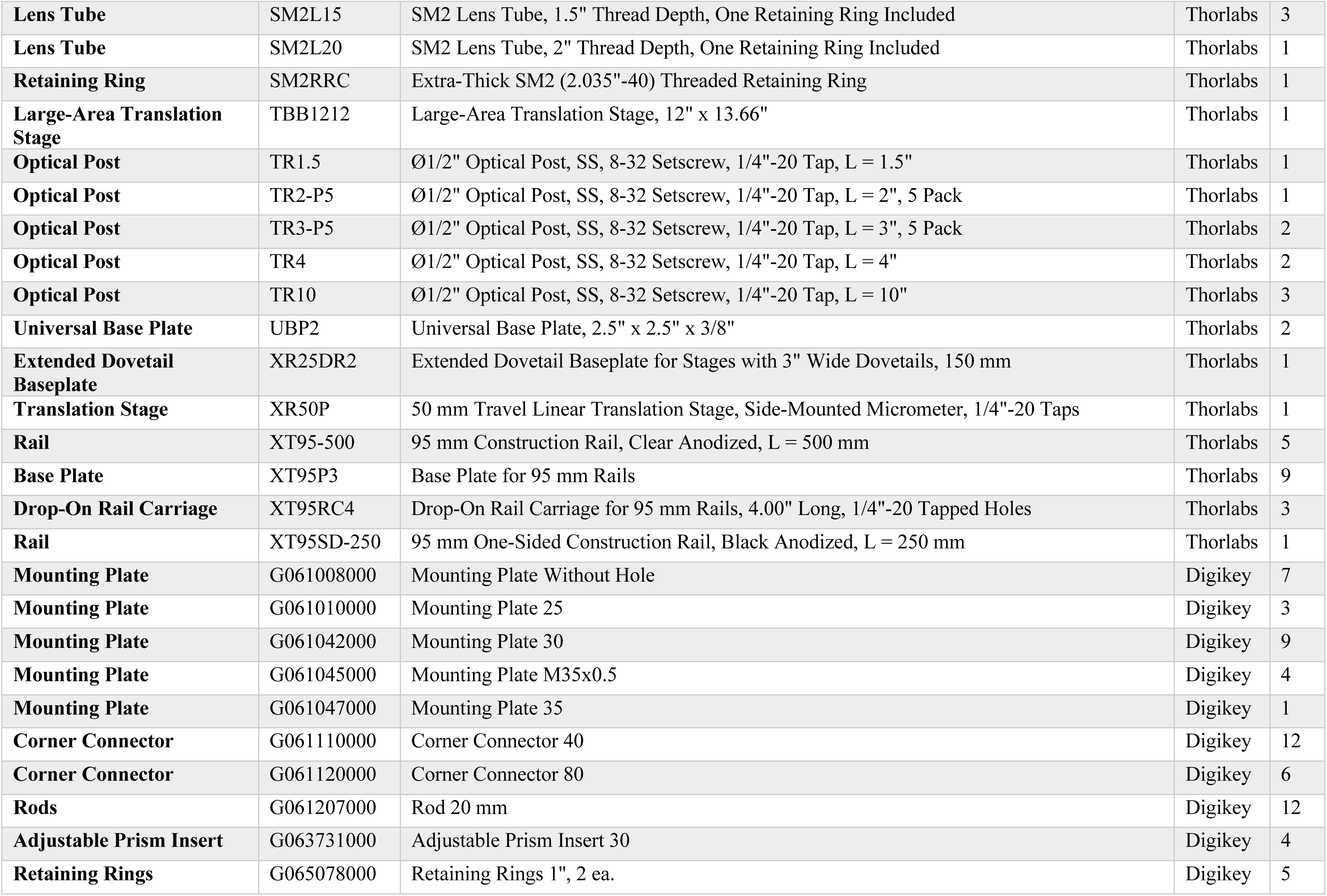
Opto-mechanics for AO-TPLSM.

**Table 4.** Custom opto-mechanics for AO-TPLSM.

| No. | Part Name | Qty | Original Part | Note |
| --- | --- | --- | --- | --- |
| 1 | main breadboard | 1 | Thorlabs, PBG3048F |  |
| 2 | z-motor adaptor | 1 | from scratch |  |
| 3 | XY stage breadboard | 1 | Thorlabs, MB1218 |  |
| 4 | Galvo mounting plate | 1 | from scratch | Need 1 for the galvo-resonant scanner system; Need 2 for a galvo-galvo scanning system |
| 5 | Galvo holder (6-mm 8135K X-galvo) | 1 | from scratch | Let the machinist know that Part5/Part5 backup must be able to fit into Part4 |
| 5 - backup | (Optional) Galvo holder (6-mm 8135K Y-galvo) | 1 | from scratch | Need it for the galvo-galvo system, but not galvo-resonant scanner system |
| 6 | resonant scanner mounting plate | 1 | from scratch | The information about the **SM1** threads can be found here: https://www.thorlabs.com/newgrouppage9.cfm?objectgroup_id=1524 |
| 7 | resonant scanner holder | 1 | from scratch | Let the machinist know that the resonant scanner must be able to fit into Part No.7 |
| 8 | extra-thick retaining ring | 1 | Thorlabs, SM2RRC |  |
| 9 | vertical mounting plate | 1 | Thorlabs, CPVMP |  |
| 10 | ER11.23 | 4 | Thorlabs, ER12 |  |
| 11 | objective mounting plate | 1 | LINOS Microbench, G061008000 |  |
| 12 | MPPC Cover | 3 | Hamamatsu, C13366-3050GA | Take the cover off from the MPPC body |
| 13 | MPPC mounting plate | 3 | LINOS Microbench, G061010001 |  |
| 14 | dichroic mount (short) | 2 | from scratch |  |
| 15 | dichroic mount (long) | 2 | from scratch |  |
| 16 | adjustable prism insert | 4 | LINOS Microbench, G063731000 |  |
| 17 | L11 mounting plate | 1 | LINOS Microbench, G061042000 |  |
| 18 | retaining ring (L12, L13, L14) | 3 | LINOS Microbench, G065078000 |  |
| 19 | mounting plate (D1) | 1 | LINOS Microbench, G061042000 |  |
| 20 | deformable mirror adaptor | 1 | from scratch |  |
| 21 | deformable mirror breadboard | 1 | Thorlabs, MB612F |  |
| 22 | EMCCD breadboard | 1 | Thorlabs, MSB46 |  |
| 23 | lens array adaptor | 1 | from scratch | Round corner for the square center counterbore |
| 24 | deformable mirror conjugation tool | 1 | from scratch |  |

The price and link for the commercial parts in **Table 1 – 3** are provided in **Supplementary Table 1**.

- Custom opto-mechanics for AO-TPLSM is listed in **Table 4**. The CAD models and ancillary drawings for the customized parts, are shared in **Supplementary Data 2**.
- Computer

#### Critical

The computer should have an extra PCI-e slot, x1 or more, for the deformable mirror controller, besides meeting the requirement for the ScanImage PXI-based system (https://archive.scanimage.org/SI2019/Supported-Microscope-Hardware_28377190.html#SupportedMicroscopeHardware-ComputerHardware).

### Software

- MATLAB (Version 2017b)
- NI DAQmx (Version 19.5.x)
- ScanImage (Version 2018b)
- Andor SOLIS
- Customized AO-TPLSM software (**Supplementary Software**)

### Chemical

These are used as 0.2 to 0.5 % (w/v) aqueous fluorescent solutions captured between a glass slide and a #1 cover slip.

- Fluorescein sodium salt (FITC) (Sigma-Aldrich, 46960, https://www.sigmaaldrich.com/US/en/product/sigma/46960)
- Sulforhodamine B (RB) (Invitrogen, S1307, https://www.thermofisher.com/order/catalog/product/S1307)
- Sulfo-cyanine5.5 (Sulfo-Cy5.5) (Lumiprobe, 27320, https://www.lumiprobe.com/p/sulfo-cy55-nhs-ester)

The Fluorescent beads are used for measuring the point spread function.

- 0.2 μm Yellow-green Carboxylate-Modified Microspheres (Invitrogen, F8811, https://www.thermofisher.com/order/catalog/product/F8811)
- 0.2 μm Red Carboxylate-Modified Microspheres (Invitrogen, F8810, https://www.thermofisher.com/order/catalog/product/F8810)

These are used to conjugate cyanine 5.5 to 2000 kDa dextran (cy5.5-dextran) as the in vivo fluorescent agent to form the guide star.

- Cy5.5 NHS ester (Lumiprobe, 27020, https://www.lumiprobe.com/p/cy55-nhs-ester)
- Amino-dextran 2000 kDa (Finabio, AD2000x250, https://www.finabio.net/product/amino-dextrans/)
- Bicine (Sigma-Aldrich, B3876, https://www.sigmaaldrich.com/US/en/product/sigma/b3876)
- Econo-Pac® chromatography columns (Bio-Rad, 7321010, https://www.bio-rad.com/en-us/sku/7321010-econo-pac-chromatography-columns-pkg-50?ID=7321010)
- DMSO, anhydrous (Thermo-Fisher, D12345, https://www.thermofisher.com/order/catalog/product/D12345)
- N-succinimidyl-acetate (TCI Chemicals, S0878, https://www.tcichemicals.com/US/en/p/S0878)
- Spin columns 100,000 Da MWCO (Sartorius Vivaspin™ 6, VS0641, https://www.sartorius.com/shop/ww/en/usd/applications-laboratory-filtration-ultrafiltration/vivaspin-6%2C-100%2C000-mwco-pes%2C-25pc/p/VS0641#)

### Biologicals

- Wild-type mouse for thalamocortical bouton imaging. (C57BL/6J, Jackson labs, Stock Number: 000664, https://www.jax.org/strain/013044)
- SST-tdTomato mouse generated by crossing SST-IRES-Cre mice (JAX, Stock Number: 013044, https://www.jax.org/strain/013044) with Ai14 mice (Jackson labs, Stock Number: 007914, https://www.jax.org/strain/007914).
- Rbp4-Cre mouse for layer 5 imaging. (MMRRC ID: 037128, https://www.mmrrc.org/catalog/sds.php?mmrrc_id=37128)
- AAV1.Syn.FLEX.NES-jRGECO1a.WPRE.SV40(Addgene, 100853-AAV1, https://www.addgene.org/100853/)
- AAV1.hSyn.FLEX.iGluSnFR3.v857.PDGFR (Addgene, 175180-AAV1, https://www.addgene.org/175180/)
- Behavioral training for head fixation and surgical procedures to make cranial windows and to inject viruses were in accordance with the animal use protocol approved by the Institutional Animal Care and Use Committee (IACUC) at the University of California San Diego.

## PROCEDURE

### I. Assembly of the AO-2P microscope with direct wavefront sensing path Timing. 5 to 10 days

This section describes the assembly of the adaptive optics two-photon microscope with direct wavefront sensing (Steps 1 to 22), as shown in **Figure 3**. Steps 1 to 16 demonstrate the procedure to build a conventional two-photon microscope with a 4-f scanning system. Steps 17 to 19 and 20 and 21 emphasize the assembly for the DM module and SHWS module, respectively. Step 22 describes the wiring for the electronics. All the assembly drawings referenced in the instructions are provided in **Supplementary Data 1**. The system should be assembled in the order and directions shown in the assembly drawings, denoted “AD”, and in **Figure 3**.

1. Mount the customized main breadboard (Thorlabs, PBG3048F, part drawing PD-01) onto the optical table (e.g., Newport, RS2000-56-12, 5’ by 6’ in extent), via four attached rails (Thorlabs, XT95-500 and XT95P3). Mount the center rail (Thorlabs, XT95-500 and XT95P3) vertically on the optical table as shown in assembly drawing AD-01.
2. Assemble the Z-motor unit (AD-02) and XY-sample stage (AD-03). Mount these components on the center rail and the optical table respectively, as shown in AD-04.
3. Assemble the lens pair L5 and L6, both of which consist of two achromatic doublets (Thorlabs, AC508-150-B), as shown in AD-05.
4. Insert the galvo scanner (Cambridge Technology, 6-mm 8315K X-galvo) and the resonant scanner (Cambridge Technology, CRS 8 kHz) into their holders (PD-05 and PD-07, respectively), as shown in AD-06 and AD-07. If a galvo-galvo scanning system is used instead of the resonant scanner, use the alternate holder (PD-05-backup) for the Y-galvo (Cambridge Technology, 6-mm 8315K Y-galvo).
5. Mount the lens pair L5 and L6, as well as the scanners, on the 4-f scanner module. Adjust the distance among each component as shown in AD-08.
6. Assemble the scan lens L7 (Newport, KPC070AR.16; Thorlabs, LB1199-B, and two of AC508-150-B; AD-09) and the tube lens L8 (Thorlabs, two of ACT508-500-B; AD-10). Mount theses assemblies on the scan module, as shown in AD-11.
7. Mount the scanning module on the main breadboard, aligning the cutout hole on the custom vertical mounting plate (Thorlabs, CPVMP, PD-09) with the cutout hole on the main breadboard. Attach a mirror (M11, Thorlabs, PF10-03-P01) to the scanning module as the entrance for the excitation beam, as shown in AD-12.
8. Mount the microscope objective (Olympus, XLPLN25XSVMP2, ×25, 1.0 NA and 4-mm working distance) to the Z-motor via an objective mounting module (AD-13), as shown in AD-14.
9. Mount a 100-femtosecond pulsed laser with tunable wavelength, e.g., Coherent, Chameleon Discovery with a wavelength range of 680 to 1300 nm, on the optical table. The main output port is followed by a shutter (Uniblitz, Shutter: LS6; Driver: VCM-D1), as shown in AD-15. Reserve the space on the optical table between the laser and the main breadboard as shown in AD-15, for a possible Pockels cell power control unit, the excitation beam expander, and the periscope.
10. Depending on whether the laser has an internal laser power control, an external power control unit such as Pockels cell (ConOptics, Cell: 350-105-02; Driver: 302RM) may be required for modulation of the excitation power. Mount the Pockels cell on the optical table (ConOptics, Mount: 102A Adjustable Mount) and align with the laser output (AD-15).
11. Assemble a telescope, formed by a pair of plano-concave lenses (L1: Thorlabs, LA1805-B, AD-16; L2: Thorlabs, LA1484-B, AD-17), as shown in AD-18. This telescope is used to expand the laser beam to fill, or slightly overfill, the aperture of the deformable mirror (ALPAO, DM75-15, pupil diameter: 13.5 mm). ***Critical step.*** The beam size varies according to the specific model of laser. Thus, the user should choose the L1 and L2 lens pair according to:

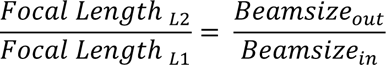

where the *Beamsize_out_* represents the diameter of collimated beam output from telescope, which should match the pupil size of the deformable mirror, and the *Beamsize_in_* represents the size of collimated beam input to the telescope, which may be approximate by diameter of the beam at the output of the power control. It is best to measure the beam diameter at the entrance of the beam expander with a beam profiler, e.g., Thorlabs, BP209IR1, and select the lens pair accordingly. For our specific realization, the beam is expanded ten-fold by lens pair L1 and L2, which are chosen to have 30- and 300-mm focal lengths, respectively. **Trouble shooting.** The expanded beam may larger than expected. We assume the laser output is collimated in the previous discussion. However, as a Gaussian beam, the laser output naturally has a slight divergence. This is compensated by adjusting the distance between lenses L1 and L2 to achieve a collimated beam at the output of the beam expander (step 24). This procedure may slightly change the magnification of the telescope, which is negligible since this application does not require a precise output beam size. Another consequence of the divergence is that *Beamsize_in_* is underestimated if it is taken as the same diameter as the laser output. The optical path from the laser output to the telescope should be minimized to reduce the effect.
12. Mount a pair of mirrors, M1 and M2 (Thorlabs, PF10-03-P01), on the optical table followed by the telescope, as shown in AD-15. The mirror pair will be used for telescope alignment.
13. Assemble the periscope, formed by a pair of mirrors, M3 and M4, as shown in AD-19. Mount M4 on the edge of the main breadboard (AD-15) and mount M3 on the optical table vertically aligned with M4.
14. Assemble the emitted photon detection unit with the following steps (AD-20).

i. Attach a custom mounting plate (LINOS Microbench, G061010000, PD-13) to the modified the cover of MPPCs (three of Hamamatsu, C13366-3050GA, PD-12).
ii. Affix an aspheric lens (L15: Thorlabs, A240-A) to the center hole of the MPPC cover with glue (Momentive, RTV103).
iii. Insert the bandpass filter (Semrock, FF01-530/55-25 for the green/yellow channel, Semrock FF01-593/46-25 for the red channel, or Semrock FF01-708/75-25 for the far-red channel) followed by a short pass filter (Semrock, FF01-790-SP-25) into the MPPC mounting plate.
iv. Fix the position of the filters with set screws. ***Critical step*.** Keep some space between the aspheric lens and the filter, as shown in AD-20. Recall that the arrow on the edge of the bandpass filter indicates the direction for the light path. Block the unused thread hole with black RTV silicone cement (Momentive, RTV103) to prevent stray photons from leaking into the MPPCs.
15. Mount the dichroic filters to the customized dichroic mount (D1: Semrock, FF665-Di02-35x37-EB, AD-21; D1’: Semrock, FF775-Di01-25×36, AD-22; D2: Semrock, FF552-Di02-25×36, AD-23; D3: FF662-FDi01-25x36, AD-23). Face the reflective coating side towards the midline of the dichroic mount; information about how to orient the dichroic filter is found at https://www.semrock.com/technical-faq.aspx.
16. Assemble the acquisition module, consisting of lenses L11 (Thorlabs, LA1765-A, AD-24), L12 (Thorlabs, LA1027-A, AD-25), L13 and L14 (Thorlabs, LA1805-A, AD-26), as shown in AD-27. Mount the acquisition block to the Z-motor. Plug in the main dichroic (D1) and three MPPC modules, as shown in AD-28. ***Critical step.*** Fill the corners and small gaps between the plates with black RTV cement to prevent stray photons from leaking to the MPPCs.
17. Assemble the deformable mirror translation stages (TS1 and TS2) on the main breadboard. Mount the deformable mirror (ALPAO, DM97-15) as well as a pair of mirrors M6 and M7 on the stage, as shown in AD-29. Depending on the placement of the periscope, one extra mirror (M5) may be required to feed the excitation beam from the periscope to the deformable mirror module. ***Critical step.*** To minimize the alignment effort when moving the TS1 (step 37), mirrors M5 and M6 should be located such that the beam between M5 and M6 is near parallel to the direction of travel of TS1. Similarly, locate mirrors M6 and M7 such that the beam is parallel to the direction of travel of TS2. To avoid mirror M7 blocking the reflected light from deformable mirror, a small incident angle, i.e., < 5°, is introduced at the deformable mirror by adjusting the position of mirror M7 on the stage TS2 (Figure 3, **AD-29**).
18. Mount lens L3 (Newport, KPX117AR.18, f = 450 mm, AD-31) on the main breadboard and attach L4 (Newport, KPX100AR.18, f = 150 mm, AD-32) to the 4-f scanner module (AD-33), as shown in AD-30. Lens pair L3 and L4 form a telescope that demagnifies the beam, three-fold, from the DM to fit the aperture of the scan mirrors.
19. Mount a mirror M8 followed by a translation stage (TS3) for the mirror pair M9 and M10 (Thorlabs, PF10-03-P01, AD-34) on the main breadboard. These are used for fine adjustment of the optical path between lenses L3 and L4, as shown in AD-30. ***Critical step.*** To minimize the alignment effort, the mirror pair M9 and M10 should be placed so that the beam between M10 on the TS3 and M11 in the scanning module is about parallel to the direction of travel of TS3. One means to achieve this is to temporarily link the M10 and M11 mirror mounts with long rods and locate the position for the M9 and M10 mirror pair before mounting them on the TS3. Similarly, place M8 in a way such that the incident beam to the M9 travels parallel to the direction of travel of TS3.
20. Attach a dichroic (D4: Semrock, FF775-Di01-25×36, AD-35) to the scanning module followed by a 4-f relay for wavefront sensing (AD-36), formed by a pair of achromatic doublet lenses (L9: Thorlabs, AC254-125-A, f = 125 mm, and L10: Thorlabs, AC254-200-A, f = 200 mm) and a pair of reflective mirrors (M12 and M13), as shown in AD-38.
21. Mount the SHWS module, consisting of an EMCCD (Andor, iXon Ultra 888) and a microlens array (Edmund, #64-483), on the main breadboard via a translation stage (TS4), as shown in AD-37 and AD-38.
22. Connect the electronics and setup the computer data file for the ScanImage; information about wiring and machine data file for the ScanImage PXI-based system is found at https://archive.scanimage.org/SI2019/28377185.html. ***Critical step.*** The frame clock of ScanImage (Primary DAQ, Port: PFI6) will be used as the trigger signal for the deformable mirror. A NOT-gate is required since the PCI-E interface of the DM controller (PEX-292144) supports trigger on falling edge only.

### II. Alignment for the AO-2P microscope

This section describes the alignment for the adaptive optics two-photon microscope with direct wavefront sensing (steps 23-42). Steps 23-37 describe the basic alignment for the microscope. Steps 38 and 39-41 emphasize the fine adjustment on the excitation path and direct wavefront sensing path respectively, to conjugate scanners, DM and SHWS to the pupil plane of the objective.

For the basic alignment and the DM conjugation (steps 23-36 and 38), we recommend tuning the laser to the most used wavelength for your application, e.g., 930 nm for excitation of GCaMP, and visualize the beam with an infrared viewer (e.g., Newport, IRV2-1300). An infrared detector card (e.g., Thorlabs, VRC4) is handy to localize the beam during alignment. We recommend you use low output power, e.g., ∼100 mW, during the alignment for safety and to minimize accidental damage to the Pockels cell.

***Timing.*** 2 - 5 days

23. Open ScanImage. Click the “Direct Mode” checkbox and adjust the power on the Power Controls Panel. Click the “Point” button on the Main Controls Panel. Flatten the DM by clicking the “Flatten DM” bottom on the AOGUI. ***Critical step***After starting the ScanImage, the scanners will be oriented towards the park angles, as set in the machine data file. Clicking the “Point” button will bring them back to 0 degrees and open the shutter for the alignment; information about the “Point” button is at https://archive.scanImage.org/SI2019/Main-Controls_28377140.html.
24. Align the laser output with the center of the beam expander. An alignment plate with a pinhole (Thorlabs, CPA1) can be used to indicate the center of the beam expander. Adjust the positioning knobs on mirrors M1 and M2 so that the beam is centered at the pinhole on the alignment plate while moving the plate back and forth along the rail to check for parallel alignment. ***Critical step.*** We recommend you temporarily remove lenses L1 and L2 from the cage system during the alignment to make the beam path more intuitive.
25. Adjust the distance between lenses L2 and L1 to collimate the output from the beam expander. ***Critical step.*** Introduce a temporary mirror after the telescope to project the expanded beam to a distant location and visualize the beam by the infrared detector card. You should not see significant changes in beam size along the beam path if the beam is well collimated. If the beam size increases along the projection, you should move lens L2 away from L1 along the cage system, and vice versa.
26. Adjust the directions of mirrors M3 and M4 to align the beam to the center of the periscope cage system indicated by the iris (Thorlabs, CP20D) that moves along the rail.
27. Adjust the orientation of mirrors M5, M6, and M7 to align the periscope output beam to the deformable mirror and tune the orientation of the DM rotation stage (Thorlabs, PR01) and TS2 to align the DM output beam to the lens L3 cage system. ***Critical step.*** Like the case in Step 24, we recommend that you temporarily remove lens L3 off from the cage system and use the alignment plate (Thorlabs, CPA1). Ensure that the beam is not blocked by M7, as discussed in step 17.
28. Adjust the distance between the DM and lens L3 to be around 450 mm by moving lens L3 along the cage rails.
29. Align the incident beam to the M9 mirror by tuning the direction the beam reflecting from mirror M8. ***Critical step.*** As we discussed in step 19, the ideal direction of the incident beam to the M9 mirror should be parallel to the direction of travel of TS3. Place an iris in front of the M9 mirror via temporary rods and adjust M8 until the beam stays on the center of the iris while the iris is moved along the rods.
30. Orient mirrors M9 and M10 to align the reflective beam to the center of iris (Thorlabs, CP20S) sliding along the rods at the entrance of the scanning module.
31. Tune the orientation of mirror M11 to make the beam go up vertically at the center of the cage system. You could take off X-galvo as well as the caps on of X-galvo cube temporally to mount an iris (e.g., Thorlabs, SM2D25D) on the top of the cube via rods for the alignment. ***Critical step.*** The dichroic mirror (D4) whose function is to reflect the emission light coming back from the sample to the SHWS, may slightly shift the excitation beam in this section of the beam path. We recommend you adjust the dichroic approximately towards the same direction as we shown in AD-38 during the alignment in this step to minimize the influence on the excitation path while aligning the SHWS path in step 39.
32. Rotate the galvo holder in the mount plate to align the excitation beam to the center of the 4-f cage system indicated by an alignment plate (Thorlabs, LCPA1) sliding along the rail. ***Critical step.*** We recommend you take leans pair L5 andL6 off from the cage system for the initial alignment. Without L5 and L6, this segment of beam path could also be used to check the collimation of the output from the beam compressor lens pair L3 and L4. Reduce the optical path between the L3 and L4 by moving the M9-M10 module along the rail if you found the beam shrinking as it travels between the two scanners, and vice versa. ***Critical step.*** After you restore the lens pair L5 and L6, double check whether the beam stays at the center of the cage system. If not, tune the L5 and L6 translation mounts (Thorlabs, CXY2) to be bring the beam back to the center of the cage system and the center of the resonant scanner, respectively.
33. Rotate the resonant scanner in the holder to align the beam to the center of the cage system between the scanner and the folding mirror. ***Critical step.*** We recommend you remove scan lens L7 for the initial alignment and tune the position of the translation mounts (Thorlabs, CXY2) of the lens tube to recenter the beam as needed.
34. Adjust the folding mirror (Thorlabs, PFE20-P01, Mount: KCB2EC) and the tube lens (L8) translation mounts (Thorlabs, CXY2) to align the beam to the center of lens L8. ***Critical step.*** Mount an iris (Thorlabs, SM2D25D) on the L8 lens tube temporarily to assist with the alignment.
35. Align the beam to the back aperture of the objective (pupil). ***Critical step.*** We recommend that you replace the objective by an iris (Thorlabs, CP20S) mounting at the pupil plane (AD-39), as shown in **Figure 7**. Adjust the non-axial position of the back aperture by sliding the objective module along the rail and changing the position of the z-motor (Thorlabs, LNR502) horizontally on its mount (Thorlabs, XT95RC4). You can check this alignment with the EMCCD as discussed later in step 44.
36. Check the conjugation between the scanners and the pupil. ***Trouble shooting.*** Ideally, the X scanner, Y scanner and the back focal plane are conjugate given the dimensions in the assembly drawing (AD-05, AD-08, AD-09, AD-10, AD-11, AD-14). However, we do recommend the users to verify the conjugation by observing the laser spot on the iris at the pupil level while scanning. You should observe little movement of the spot if they are well conjugated. If you observe large movement of the spot, translate the pupil level along the axial direction by the Z-motor to the level where you see little movement of the spot during scanning.
37. Align the collection path. Adjust the main dichroic mirror (D1 or D1’) to maximize the image brightness while imaging the sulfo-Cy5.5 solution (excitation beam: 1250 nm) with the far-red channel. Similarly, adjust D2 and D3 while imaging FITC (excitation wavelength: 930 nm) and sulfo-rhodamine B (excitation wavelength: 1030 nm) with green-yellow channel and red channel, respectively.
38. Conjugate the deformable mirror to the back focal plane of the objective. ***Critical step.*** To find the conjugate plane for the DM, temporarily replace the deformable mirror by a scratched mirror (AD-41) and image it by a camera whose sensor is at the level of the objective back focal plane (AD-40). Slide stage TS1 until the camera obtains the clear and sharp image of the scratched mirror, as shown in **Figure 8**. Put the DM back after you find the conjugate plane and adjust the alignment if necessary.
39. Excite the sulfo-Cy5.5 solution with 1250 nm excitation beam and align the emission beam to the wavefront sensing path from dichroic mirror D4 to L10 by tuning the orientation of D4 and mirrors M12 and M13. ***Critical step.*** Mount a camera, the same one you used for the DM conjugation, in the cage system for the initial alignment if you find the emission light too dim to be visualized by the IR viewer or IR card. ***Critical step.*** Use dichroic mirror D1, and not D1’, as the main dichroic so that most of the emitted light will go to the wavefront sensing path rather than the collection path.
40. Temporarily remove the lens array from the SHWS. Open the Andor SOLIS software. Adjust the lateral and vertical position of the EMCCD to locate the emission beam at the center of the EMCCD. Adjust mirror M12 and M13 until the beam position on the EMCCD stays fixed as the stage TS4 travels along the axial direction (**Figure 10a,b**). Adjust the distance between lens pair L9 and L10 until the beam size does not change as the EMCCD travels axially. **!Caution** Always keep the short pass filter (Semrock, FF01-790-SP-25) in front of the highly sensitive EMCCD to protect it from scattered light from the excitation laser.
41. Mount the lens array back in the SHWS and slide it along the rail until it focuses the emission light on the EMCCD. ***Critical step.*** Mount a stopper on the rail (Thorlabs, ERCPS) to mark the relative position for the lens array, as the array will be removed and replaced several times for testing of conjugation and for calibration. Measure and calculate the distance between to lens array and the EMCCD sensor (AD-37), marked as Dist1.
42. Find the plane that conjugates to the objective back focal plane on the SHWS path. ***Critical step.*** Mount a paper target at the level of the objective back focal plane (AD-42) and illuminate it with a flashlight (laser off), as shown in **Figure 9**. Remove the lens array and move the EMCCD axially to the conjugate plane, where it obtains the sharp image of the paper target. Read the micrometer on stage TS4 and mark this position as P1. Restore the lens array and move stage TS4 axially for Dist1 to bring the lens array to the conjugate plane (marked as P2).

### IIIa. AO calibration

This section describes the calibration of the AO-TPLSM system (steps 43 to 48). The purpose for the AO calibration is to map the phase shifts for each Zernike mode at 1-μm root-mean-square amplitude on the DM to the spot shifts on the SHWS, which will be used to calibrate and calculate the command signals to the DM module to correct for sample aberration.

*Timing.* 1 day.

43. Map the DM to the field of the SHWS as follows

i. Remove the lens array from the SHWS, keeping the short pass filter, and move the EMCCD to P1.
ii. Replace the objective by a mirror mounted at the back focal plane (AD-43).
iii. Mount an ND filter at the end of the lens pair L1 and L2 beam expander (e.g., Thorlabs, ND30A, Optical Density: 3.0) to protect EMCCD by reducing the laser power.
iv. Flatten the DM.
v. Tune the laser wavelength to 740 – 750 nm.
vi. The calibration beam incident on the DM will be reflected by the pupil mirror and project to the EMCCD, where the DM will be seen as a bright disk with sharp edge in the SOLIS software after adjusting the contrast and brightness, as shown in **Figure 10d**.
vii. Use the ROI tool in SOLIS to draw circle on the edge of the DM. **!Caution** Always double-check the ND filter for the steps that directly reflect the laser to the EMCCD. ***Trouble shooting.*** The sharp edge of bright disk is the edge of the DM because the DM is conjugated to the EMCCD in this setting.

i. If you cannot see the sharp edge, it is likely that the incident beam to the DM is not sufficiently expanded or the DM, Pupil, and EMCCD are not conjugated to each other.
ii. If the edge is not circular, it means the beam was cropped by something in the optical path, often the pupil mirror or the scanners. Move the pupil mirror horizontally to test if it fixes the problem before checking the alignment of the system.
iii. If the brightest part of the laser is not centered at the disk, it means the incident beam to the DM is not well aligned. Readjust mirrors M6 and M7 to center the laser to the DM.
44. It is critical for the AO calibration to map the DM and the objective back aperture at the same location on the SHWS. These steps describe a process.

i. Temporarily remove the lens array and keep the EMCCD at P1. Remove the ND filter and put the objective back.
ii. Excite the Cy5.5 solution with the 1250 nm excitation beam, as shown in **Figure 10a**. The emission light collected by the objective will be seen a disk with sharp edge (back aperture disk) in the SOLIS software.
iii. Adjust the objective mounting module non-axially until the back aperture disk is concentric with the DM edge, as shown in **Figure 10a,d**.
iv. Double check the alignment after the adjustment by moving the EMCCD to P2 to confirm whether the beam position on EMCCD the stays still (**Figure 10a,b**). If not, adjust the system as described in step 40.
v. Replace the lens array and obtain an image with the SOLIS software as a sample reference pattern (**Figure 10c**).
45. Set up the system in the same manner as in step 43. Move the EMCCD to P2 and tune the direction of the pupil mirror to fit the calibration beam into the edge you have drawn at P1, as shown in **Figure 10e**. Mount the lens array back. Finely tune the pupil mirror to overlap the current pattern on the EMCCD (calibration reference pattern, **Figure 10f**) to the sample reference pattern as much as possible (**Figure 10g**). ***Trouble shooting.*** The mismatch of the calibration reference pattern with sample reference pattern will reduce the dynamic range for wavefront sensing. Use the following sequence if the patterns cannot be overlapped.

i. Adjust the correction ring on the objective while acquiring the sample reference pattern until the new pattern best matches the calibration reference pattern. Fix the correction ring at this position for the rest of the procedure.
ii. If mismatch still occurs, it may indicate that the excitation beam is not collimated, especially if you see the grid size differs in the two-reference patterns. Move stage TS3 along the rail to change the divergence of the emitted light. Then recollimate the wavefront sensing beam as described in step 40.
46. Close the SOLIS software to release control of the EMCCD and initialize the DM and SHWS from MATLAB by clicking “initialize DM” and “initialize SH” bottom on the AOGUI. Click “Camera config” to set the temperature as -60 °C for the EMCCD and turn it on. ***Critical step.*** Acquire images before the calibration to find a laser power and exposure time for clear images after the EMCCD reaches its target temperature.
47. Run the first section of the code “DM_SHWS_calibration_Ctr.m”, which will display each of the 60 individual Zernike modes on the DM and acquire an image of the deflected spots on the SHWS (**Figure 11**). The code will also download a flat pattern to the DM to acquire a wavefront spot pattern for calibration reference. After the first section of the code is complete, turn off the shutter and run the second section of “DM_SHWS_calibration_Ctr.m” to obtain a background image. Save “DM_calibration_shdata”, “DM_calibration_shdata_ref” and “DM_calibration_shdata_ref_bg” for the analysis in next step.
48. Run “DM_SHWS_calibration.m”, which will analyze the deflection of the spots on the SHWS for each Zernike modes. ***Critical step***. Before running the code, adjust the parameter for the center and the radius of the field-of-view (FOV) (lines 65 to 67) based on the back aperture disk on EMCCD as previously obtained in step 44. ***Trouble shooting.*** If random dots and/or small objects are detected as focused spots on the EMCCD image, the detection threshold can be adjusted (shstruct.thresh_course_img, line 133) to excludes these spots. ***Trouble shooting.*** This part of the code will show the putative pattern on the DM versus the ideal Zernike modes. The putative pattern should look like the ideal one, except that it may be rotated or inverted. If it appears very different from the ideal pattern and there is nothing wrong with the calibration, i.e., no extra spots, perfect overlap of the pupil and DM, and well-aligned wavefront sensing pathway, you should contact the DM manufacturer to calibrate the DM influence function.
49. Click “Camera config” to set the temperature as 20 °C for the EMCCD and release control of the EMCCD from MATLAB by clicking “deinitialize SH” after it reaches 20 °C. The temperature of the EMCCD can be checked by click “check temp”. **!Caution**. Always use the temperature control to return EMCCD back to the room temperature before turning it off.

### IIIb. System aberration calibration

The purpose for the system aberration calibration is to find a pattern on the DM that compensates for the aberration introduced by the imaging system itself, which includes the imperfections of the microscope as well as aberrations induced by the cover glass of the cranial window (steps 50 to 52). We suggest the system aberration should be corrected for the all the wavelengths that are commonly used during experiments, as some components of the aberration, e.g., spherical aberration, vary with the wavelength (**Figure 13**). We recommend the use of aqueous solutions of the fluorescent dyes fluorescein, sulforhodamine B, or cyanine 5.5 to correct the system aberration for 930, 1030, and 1250 nm excitation, respectively.

***Timing.*** 2-3 hours.

50. Flatten the DM by clicking the “Flatten DM” bottom on the AOGUI. Image the central 100 μm × 100 μm field of the fluorescent sample at the depth of 50 μm under the cover glass (thickness number 1). ***Critical step.*** Use dichroic mirror D1 as the main dichroic for FITC, sulforhodamine B imaging and use the D1’ for of sulfo-Cy5.5.
51. Run “sensorlessWF_sys_abr_descend.m” to implement a gradient-descent algorithm that optimizes the amplitude of each Zernike mode to maximize the average intensity (**Figure 12**). In each iteration, the increments of the Zernike mode coefficients are computed based on the gradient of mean intensity changes to update the pattern on the DM. Tip, tilt and defocus are excluded. The coefficients of the Zernike modes usually became stable after 30 to 50 cycles of optimization (**Figure 13**); you can set the number of iterations at line 18 of the code. The resulting Zernike coefficients will be referred as system aberration correction coefficients.
52. Validate the system aberration correction coefficients by measuring the point spread function of the 200-nm fluorescent beads after loading the system aberration correction pattern on the DM (**Figure 14**).
53. Image the aqueous solution of sulfo-Cy5.5 after system aberration correction (main dichroic: D1’, 100 µm x 100 μm field, 50 μm under the cover glass). Switch the main dichroic to D1 to obtain a sample reference pattern on the SHWS, which will be used for sample aberration correction.

### IIIc. System verification

Improper calibration is a common cause for poor performance with AO in actual experiments. It can even lead to image degradation after aberration correction. To avoid this, we recommend that the performance of the AO-TPLSM system is tested on fiber test sample (steps 54 and 55 and the associated sub-steps) before use of AO-TPLSM with living preparations.

***Timing.*** 2-3 hours.

54. Prepare a double-layer cover glass sample.

i. Glue two number 1 cover glasses with optical adhesive (Norland optical adhesive 61).
ii. Cut a 5 mm × 5 mm piece of Kimwipe, place it on a slide.
iii. Add a drop of the 5 % fluorescein and a drop of the 5 % Cy5.5 solutions, wait till the solutions dry, add a drop of instant adhesive (Loctite 4014), and close with the double-layer cover glass.
55. Perform verification

i. Start ScanImage. Turn on the EMCCD and set the CCD temperature to -60 °C.
ii. Initialize the DM and flatten it. Acquire a 50 μm × 50 μm FOV image of a branch of a fiber (**Figure 15a**).
iii. Load the system aberration correction coefficients for acquisition at 930 nm to the DM.
iv. Acquire an image for the same branch of the fiber with the same laser power as step 55 (**Figure 15b**).
v. Use dichroic mirror D1’ as the main dichroic. Tune the laser to 1250 nm for cy5.5 and load the system aberration correction coefficients for acquisition at 1250 nm.
vi. Image the same FOV as in step 54.iv with the far-red channel to confirm the existence of cy5.5 in the FOV.
vii. Replace dichroic mirror D1’ with mirror D1 to transmit the far-red emission light of cy5.5 to the wavefront sensing path.
viii. Obtain the sample aberration pattern on the SHWS with the SOLIS software by a long exposure, i.e., 0.5 s, during scanning.
ix. Save the sample aberration pattern (**Figure 15e**). Run “sample aberration correction” to compute the coefficient for sample aberration correction (**Figure 15f,g**).

**Figure 15.**
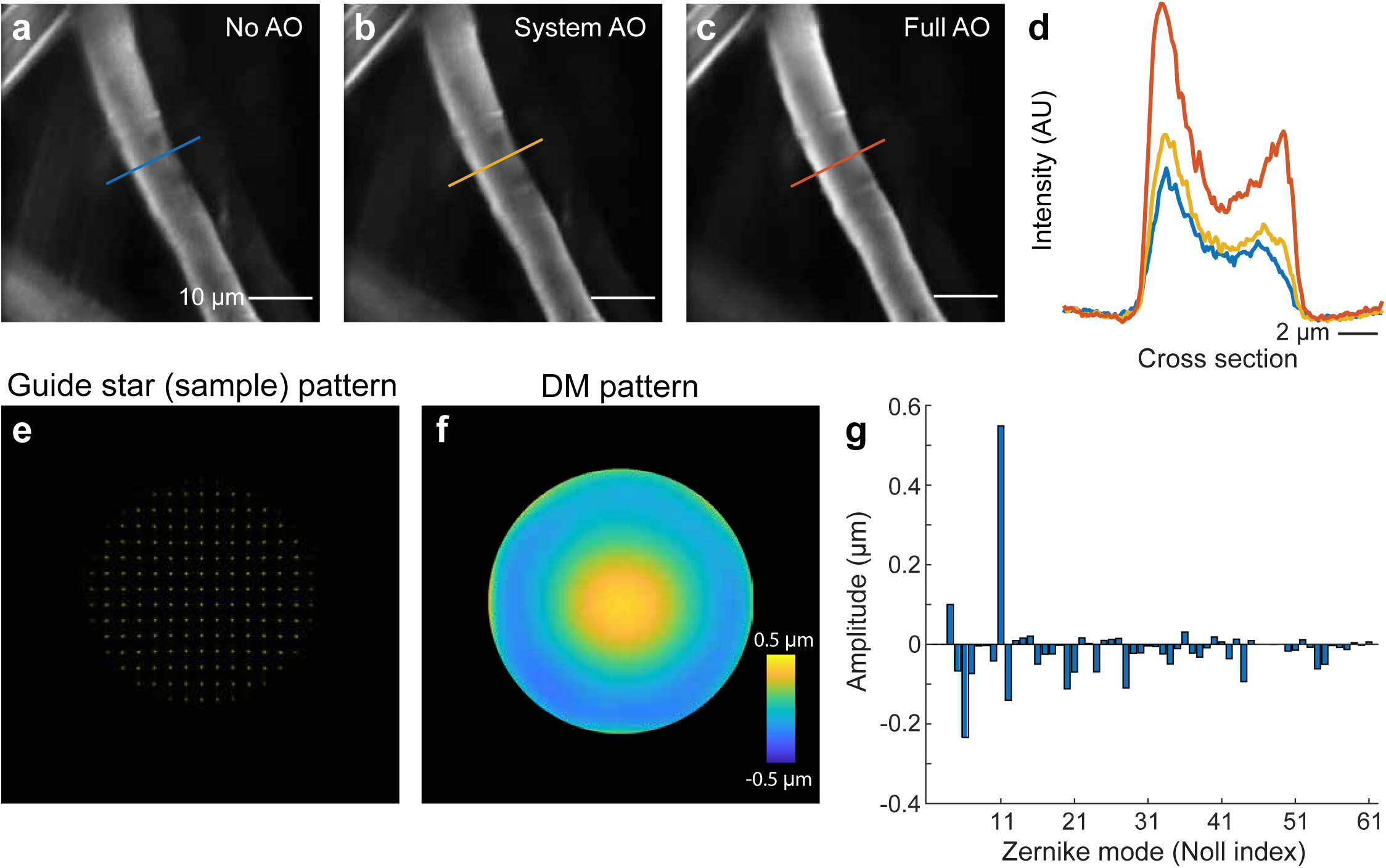
System verification by correcting the aberration of the double-layer cover glass. **a** to **c**, Images of a fiber stained with rhodamine B. Scale bar, 10 μm. **a**, Images taken without any AO correction. **b**, Images taken after correcting the system aberration. **c**, Images taken with both system aberration and sample aberration corrected. **d**, Signal profiles along the line in **a** to **c**. Scale bar, 2 μm. **e**, Spot patterns formed by the SHWS from the descanned guide star signal for inferring the sample aberration. **f,** Reconstructed wavefront that applied to the DM. **g**, Zernike coefficients of the reconstructed wavefront. ***Critical step.*** The pattern for the double-layer cover glass should be dominant by the spherical aberration (Noll index 11) in a well calibrated system.
x. Add the sample aberration correction coefficient to the system aberration correction coefficient for 930 nm and send it to the DM.
xi. Acquire an image for the fiber using the same FOV and same power as in steps 55.iv and 55.vi (**Figure 15c**).
xii. Compare the three images. A good calibration should give you about two-fold increase of the SNR in this setting (**Figure 15d**).

### IV. Synthesis of Guide Star (2000 kDa Cy5.5-dextran)

This section (steps 56 to 68) describes the procedure to synthesize 3 % (w/v) 2000 kDa Cy5.5-Dextran which will be used for generating the guide star signal for sample aberration correction *in vivo*.

*Timing.* 2 days.

56. Prepare 50 mM bicine buffer at pH 8.5; adjust with NaOH. Store at 4 °C.
57. Dissolve 166.3 mg of amino-dextran in 1039.4 μL Bicine buffer; add up to an extra 200 μL buffer if it does not dissolve well as the solution is close to the saturation limit. Use a 50 ml Falcon tube for this process.
58. Dissolve 5 mg Cy5.5 NHS ester in anhydrous DMSO by adding 651.3 µl of DMSO to the 5 mg Cy5.5 NHS ester in the Falcon tube. Gently swirl and then centrifuge down for 30 s. **!Caution.** Thus solution is unstable and should be used immediately after preparation.
59. Mix dextran and Cy5.5 NHS ester, vortex, and incubate for 2 h at room temperature. The solution should get a little warm from the reaction.
60. Dissolve 32.9 mg of N-succinimidyl-acetate (quencher) in 140 µl of anhydrous DMSO in a 1.5 ml tube. **!Caution.** Thus solution is unstable and should be used immediately after preparation. N-succinimidyl-acetate reacts with H2O, so always store N-succinimidyl-acetate in N2 and work quickly after opening.
61. Add N-succinimidyl-acetate solution to the reaction and incubate overnight at 4 °C.
62. Suspend Sephadex G25 (in the separation column box) in PBS pH 7.4, for one batch use either 2 g or 3 g, add up PBS to 50 ml and allow some time to equilibrate (e.g., overnight at 4 °C).
63. Decant PBS and leave 5 ml PBS on top of the Sephadex G25, vortex and add to chromatography columns; wait for the Sepadex to compact and add the top filter. Slowly push it down until it touches the Sephadex. Don’t push to far this will compact the beads too harsh. Poor the top PBS and equilibrate the column with 10 ml fresh PBS. Never let the column run dry.
64. Add 2-3 ml of PBS pH 7.4 to the reaction mixture and load onto the chromatography column.
65. Wait for the Dye to completely enter the Sephadex.
66. Add some PBS (fill the column) and collect ‘blue’ fraction from the column (50 ml Tube). Stop collecting when the bottom of the column becomes transparent.
67. Load ‘blue’ fraction into spin column(s) into spin column(s), and centrifuge for 4 h at 4 °C @ 4000 g (swinging bucket rotor) or 5000 g (fixed rotor).
68. Collect the supernatant, which is Cy5.5-dextran; dilute it to 5 mL with saline, sterile filter, and store at -20 °C.

### V. AO-2P Imaging with microvessels based guide-star

This section (steps 69 to 75) describes the process (**Figure 16**) to imaging structure (**Figure 17**) and function (**Figure 18**) of the mice brain with AO-TPLSM. The animals should be prepared in the same manner as for conventional TPLSM, including the viral injection, craniotomy, or other special treatment that may be performed for a specific experiment. To prevent the incident light cone from being blocked by the edge of the optical window, for deep layer imaging, we recommend use of a cover glass that is larger than the area of interest.

**Figure 16.**
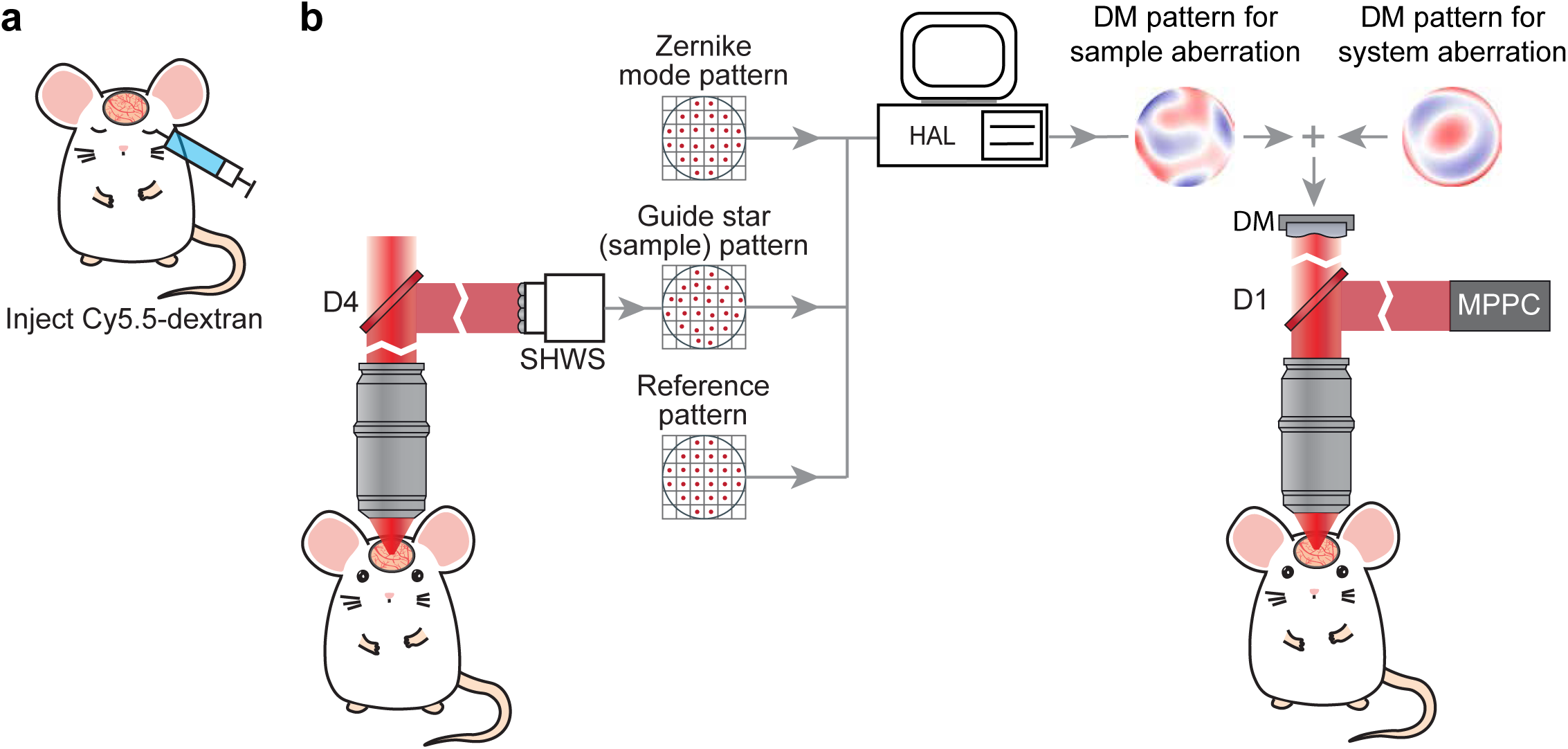
*In vivo* experiment workflow. **a**, Schematics of the guide star (Cy5.5-dextran) delivery by retro-orbital injection. **b**, Schematic diagram of the AO correction. Spot patterns formed by the SHWS from the descanned guide star signal in the capillary, together with the sample reference pattern and AO calibration results of the DM Zernike modes, are used for calculating the DM pattern for correcting the sample aberration. The sample aberration correction pattern is added to the system aberration pattern for full AO correction.

**Figure 17.**
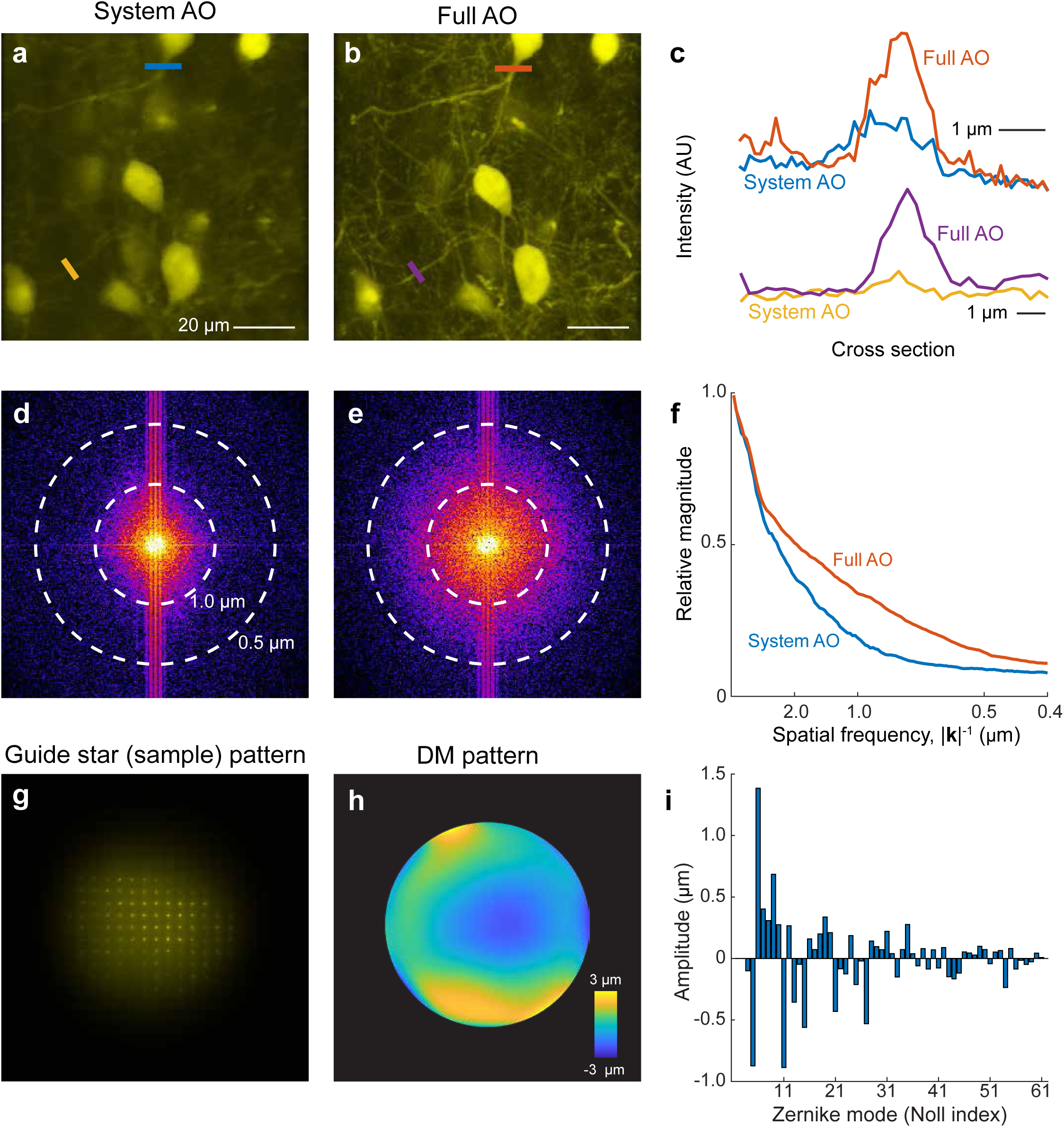
Morphological imaging with AO correction. **a**,**b**, SST^+^ neurons in mouse somatosensory cortex. Images were acquired at 675 - 690 μm below the pia with system (**a**) or full AO correction (**b**) using excitation wavelength λ = 1030 nm. The SST^+^ neurons were labeled with tdTomato. Scale bar, 20 μm. **c**, Signals profiles along the lines in **a** and **b**. Scale bar, 1 μm. **d** and **e**, Spectral power as a function of spatial frequency k for the images in **a** and **b**. **f**, |k|-space plot of the spatial frequency with system AO and full AO correction. **g**,**h**, Spot patterns on the SHWS (**g**) and reconstructed wavefront phase map on DM (**h**) for correcting the sample aberration in the **a**. **g**, Zernike coefficients of the reconstructed wavefront.

**Figure 18.**
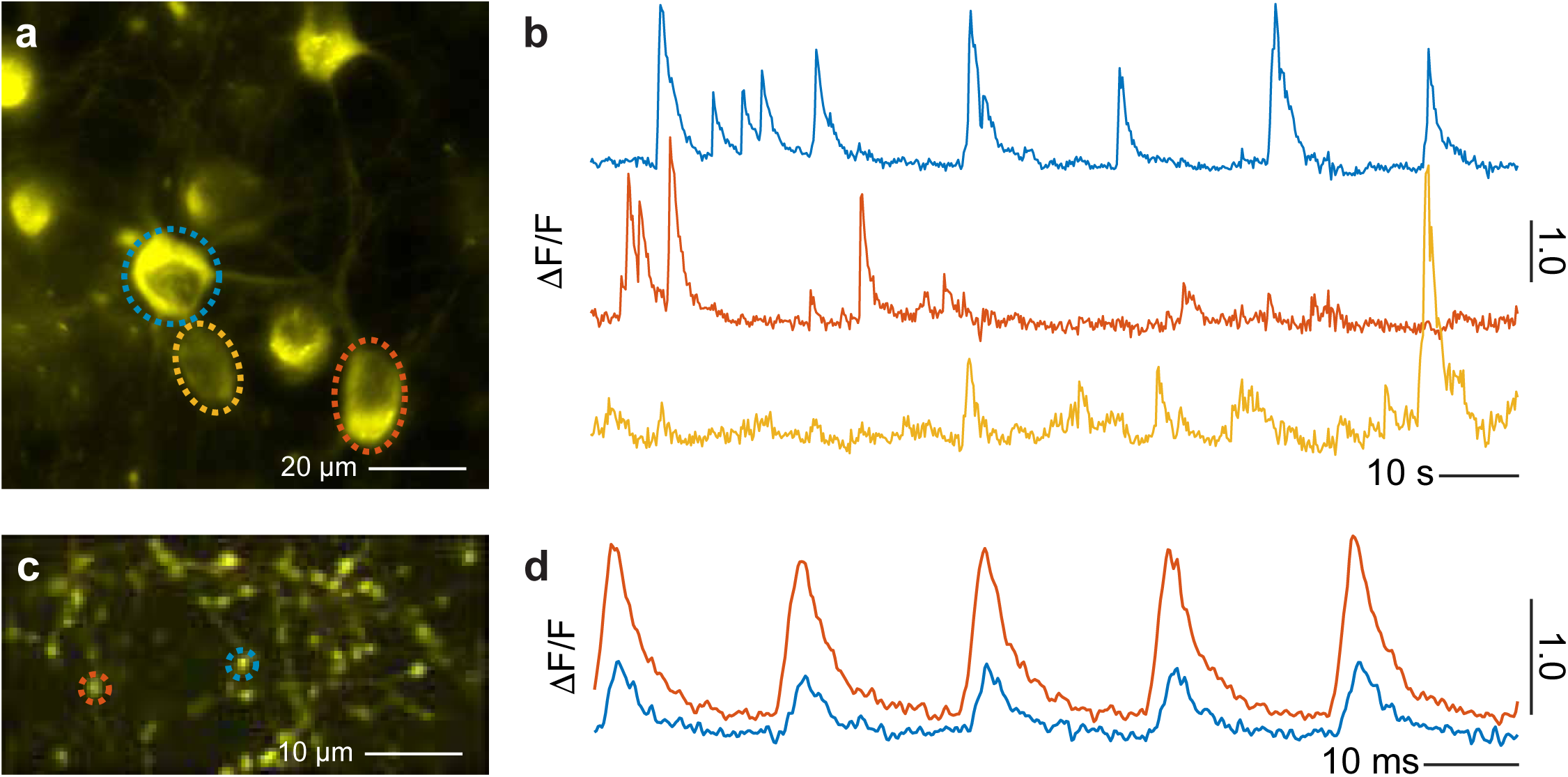
Functional imaging AO correction. **a**, *In vivo* two-photon imaging of the layer 5 neurons of vS1 cortex which were labelled with jRGECO1a. Data were obtained at 630 μm below the pia with full AO correction using excitation wavelength λ = 1030 nm. **b**, Calcium signal in the ROIs defined by the dashed circles in panel **a**. **c**, *In vivo* two-photon imaging of the thalamocortical boutons which were labelled with iGluSnFR3. Data were obtained at layer 4 of vS1 cortex with full AO correction using excitation wavelength λ = 970 nm. **d**, Glutamate dynamics responding to 5 Hz air puff in the ROIs defined by the dashed circles in panel **c**.

*Timing.* 1 day.

69. Briefly anesthetize the mice with isoflurane and deliver 50 µl of 3 % (w/v) solution in saline of Cy5.5-dextran (**Procedure IV**) into the lumen of blood vessels via retro-orbital intravenous injection^18^. The Cy5.5-dextran can circulate for more than 4 hours in the blood and provide a stable guide star signal during this period.
70. Start ScanImage. Initialize the DM and load the system aberration correction coefficients to the DM based on your imaging wavelength. Turn on the EMCCD and set the temperature as -60 °C. Find the FOV within the window formed by a craniotomy^20^ or thinned bone^19, 28^. ***Critical step.*** Make sure the cranial window is horizontal, i.e., perpendicular to the optical axis. A tilted window will alter the sample aberration correction such that the FOV is shifted after sample aberration correction.
71. Use dichroic mirror D1’ as the main dichroic. Tune the laser to 1250 nm for Cy5.5 and load the system aberration correction coefficients for 1250 nm. Image the microvessels in the region of interest with the far-red channel. ***Critical step.*** As the tissue aberration varies gradually as the spatial location, we recommend the user to perform the sample aberration correction for individual 100 µm × 100 µm FOVs, which is a phenomenological limit to the isoplanatic region^11, 22^.
72. Replace dichroic mirror D1’ with dichroic D1 so that the far-red emission light of Cy5.5 is mainly directed to the wavefront sensing path rather than the collection path. Obtain the sample aberration pattern on the SHWS by a long exposure, i.e., 1 to 30 s, with scanners scanning the FOV to correct depending on signal-to-noise (**Figure 17g**). Adjust the EM-CCD gain to achieve a good signal of the sample aberration pattern. Save the sample aberration pattern. ***Critical step.*** The SNR of the sample aberration pattern is critical to the AO correction. To improve the signal for deep tissue, we recommend use of the ScanImage microROI function. By selecting ROIs on the microvessel, the galvos will only scan on the vessel during the period of exposure, voiding collecting the noise from the location without a vessel in the field of view.
73. Run “sample aberration correction”, which computes the shifts of the center of individual spots in the sample aberration pattern from the reference pattern and decomposes the shifts to the Zernike mode pattern to obtain the coefficient for sample aberration correction (**Figure 17h,i**).
74. Add the sample aberration correction coefficient to the system aberration correction coefficient of wavelength that you are going to use and send them to the DM. At this point, both the system aberration as well as the sample aberration for that FOV are corrected. Start the recording (**Figure 17a-c**). ***Critical step.*** In a successful sample correction, the user should see significant increase in SNR (**Figure 17a-c**) and spatial resolution which can be visualized by a 2-D FFT of images (**Figure 17d-f**). ***Trouble shooting.*** If the image is not improved by the correction, it is usually caused by the low SNR of the sample aberration pattern. The following process can help.

i. Extend the exposure time during wavefront sensing.
ii. Increase the EMCCD gain in SOLIS; typically, 2 to 80.
iii. Use ScanImage microROI function to select ROIs on the microvessel to let the scanners only scan on the vessel during the exposure.
iv. Reduce the FOV to 50 mm × 50 mm.
v. Use a Zernike mode pattern in which the defected spots are weighted by their signal-to-noise ratio when calculating the Zernike coefficients.
vi. Check the AO calibration (**Procedures IIIa** and **IIIc**).
75. If you have another FOV to record, center at the new FOV, and repeat steps 69 to 74.

## Supporting information

Supplemental data, tables, and software

## Author contributions statements

DK and RL guided this project, RL and PY designed the microscope, which is based on a prior design by RL, PY carried out fabrication and testing of the hardware, TB, RL and MT developed the method for synthesize the cy5.5-dextran, RL and PY performed all in vivo measurements, and DK and PY wrote the manuscript. DK attended to the plethora of university rules and forms that govern research compliance, export control, and environmental health and safety, including the ethical use of animals as well as the use of chemicals, hazardous substances, lasers, and viruses.

## Acknowledgments

We thank Stephen Adams for assistance with cy5.5-dextran synthesis and Beth Friedman for assistance with animal preparation. This work was funded by the National Science Foundation, grant PHY153264, and the National Institutes of Health, grants U24 EB028942, R35 NS097265, and U19 NS107466.

## Competing interests

None

## Notes

### Competing Interest Statement

The authors have declared no competing interest.

