## Supplemental data, tables, and software for "Guide to the construction and use of an adaptive optics two-photon microscope with direct wavefront sensing": AO_TPLSM_Supplementart Information.docx

Supplementary Information should fit into one of three categories:

1. EXTENDED DATA: Extended Data is an integral part of the paper and only data that directly contribute to the main message should be presented. These figures will be integrated into the full-text HTML version of your paper and will be appended to the online PDF. There is a limit of 10 Extended Data figures, and each must be referred to in the main text/online methods ( cited as ‘Extended Data Figure 1’, ‘Extended Data Figure 2,’ etc.). Each Extended Data figure should be of the same quality as the main figures, and should be supplied at a size that will allow both the figure and legend to be presented on a single A4 page. Each figure should be submitted as an individual .jpg, .tif or .eps file with a maximum size of 10 MB each, using the ‘Figure File’ article type in our Manuscript Tracking system.

2. SUPPLEMENTARY INFORMATION: Supplementary Information is material that is essential background to the study but which it is not practical to include in the PDF version of the paper (for example, video files, large data sets and calculations). Each item must be referred to in the main manuscript/online methods and detailed in the attached Inventory of Supplementary Information. Tables containing large data sets should be in Excel format, with the table number and title included within the body of the table. All textual information and any additional Supplementary Figures (which should be presented with the legends directly below each figure) should be provided as a single, combined PDF. Please note that Supplementary Information is not copyedited and we cannot replace any Supplementary Information after the paper has been formally accepted unless there has been a critical scientific error. Supplementary Figures and other items are not required to be called out in your manuscript text, but should be numerically numbered, starting at one, as ‘Supplementary Figure 1’, etc., not SI1.

3. SOURCE DATA: We encourage you to provide Source Data for your Figures whenever possible. Full-length, unprocessed blots and gels must be provided as Source Data for any relevant Figures, and should be provided as individual PDF files (one for each figure) containing all supporting blots and/or gels with the linked figure noted within the PDF file. Data underlying any relevant graphical Figures should be provided as individual Excel files (one for each figure) with the linked figure noted within the Excel file. We encourage deposition of all types of Source Data into a relevant repository, for example figshare (<https://figshare.com/>) or the Image Data Resource (<https://idr.openmicroscopy.org>). Please see more information about data availability: <https://www.nature.com/nature-research/editorial-policies/reporting-standards#availability-of-data>.
