## Supplemental data, tables, and software for "Guide to the construction and use of an adaptive optics two-photon microscope with direct wavefront sensing": AD-01.pdf

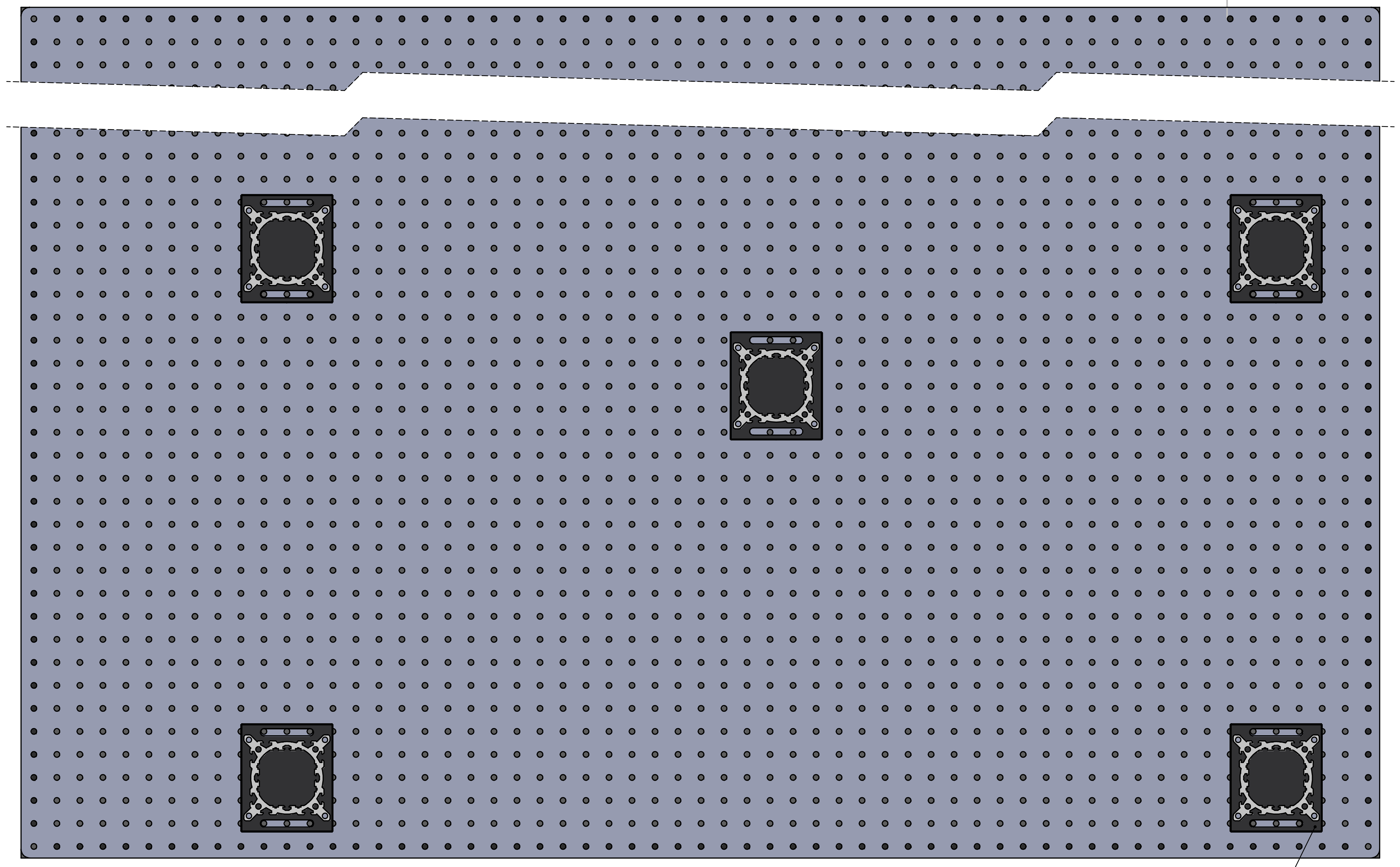

9 x Thorlabs,XT95P3

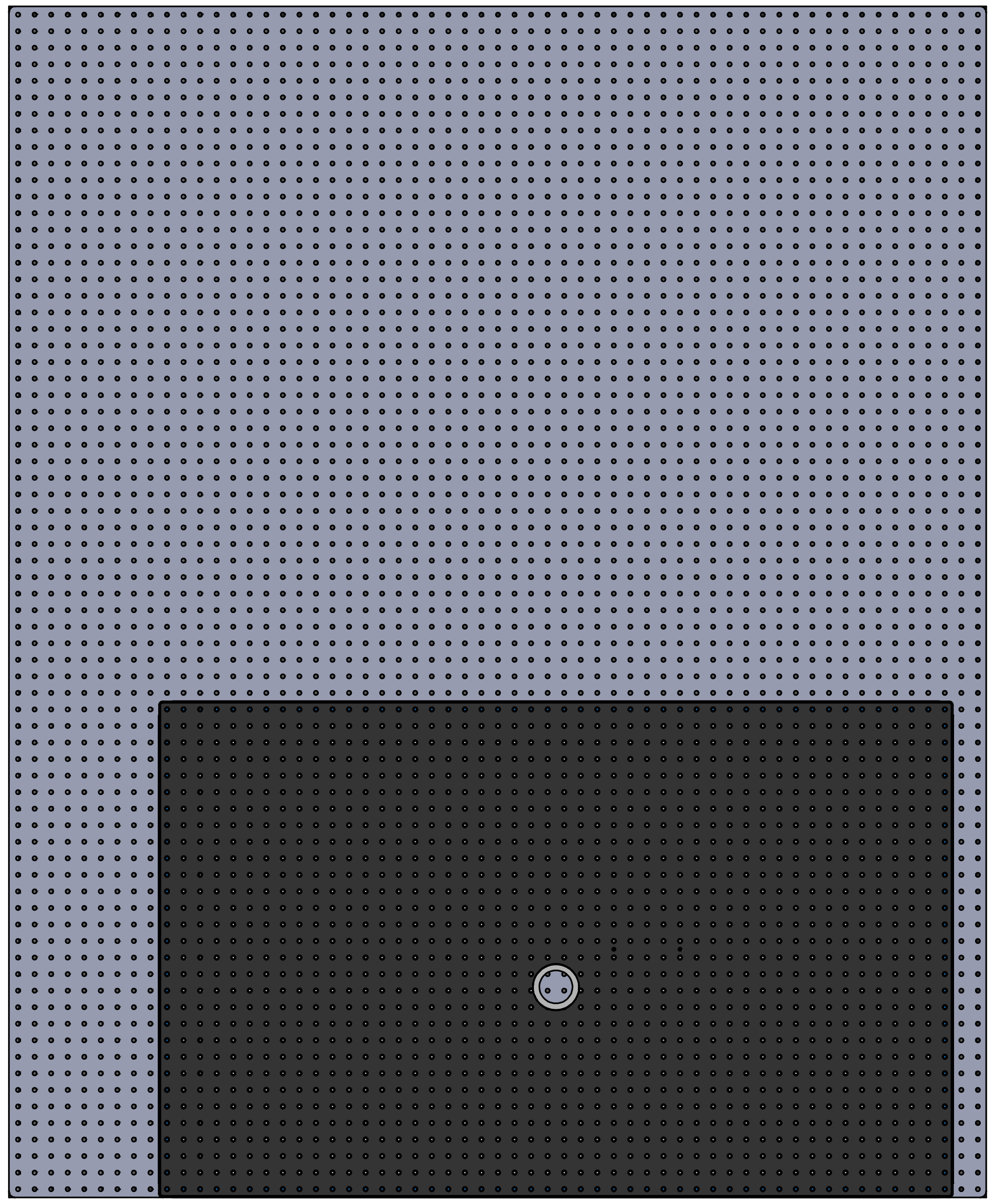

Thorlabs, PBG3048F, PD-01

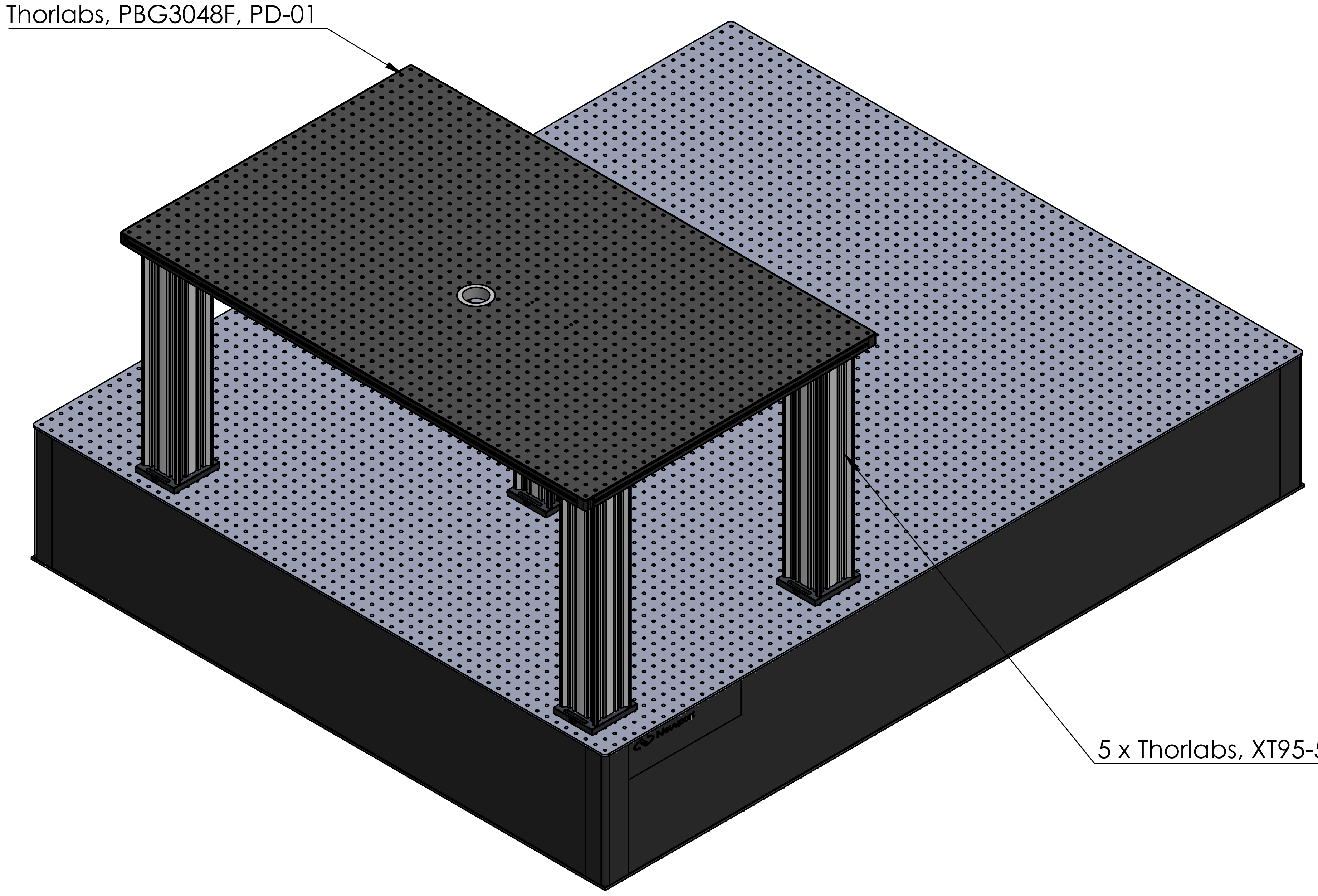

5 x Thorlabs, XT95-500

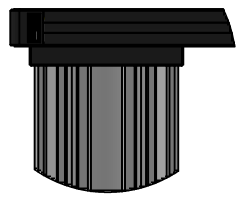

DETAIL A
SCALE 1 : 5

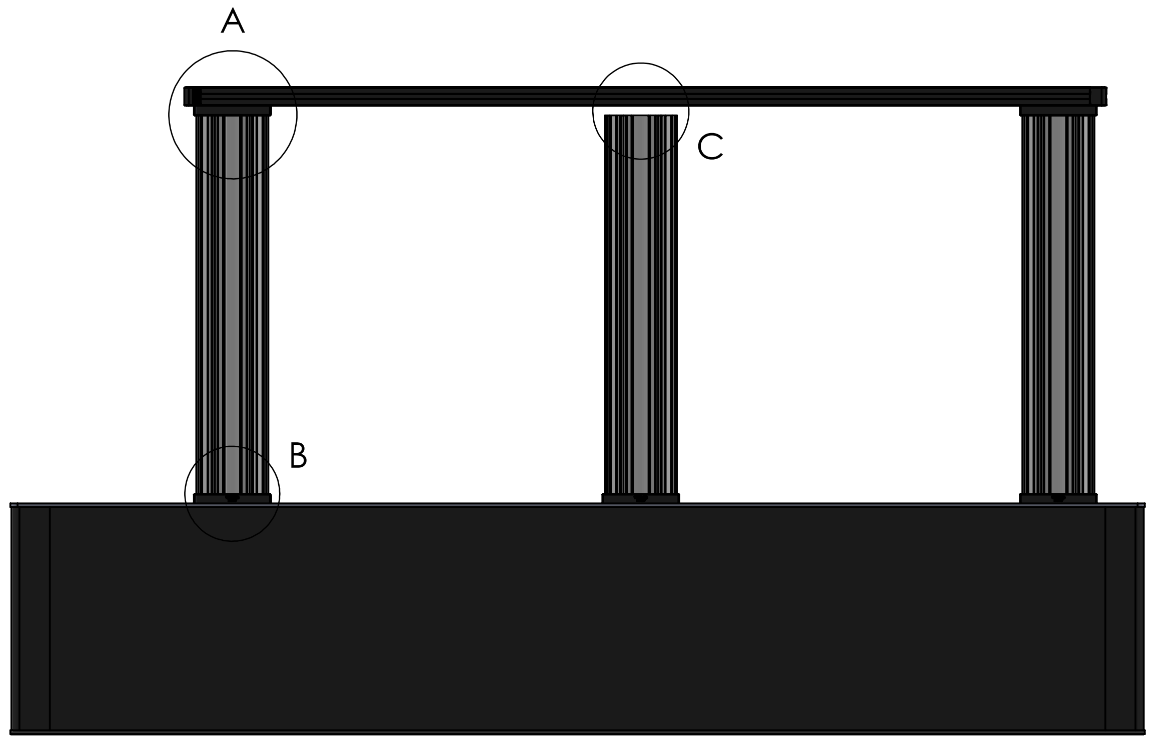


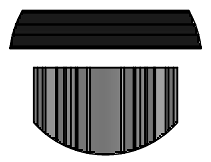

DETAIL C
SCALE 1 : 5

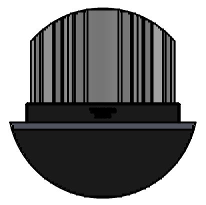

DETAIL B
SCALE 1 : 5

| UNLESS OTHERWISE SPECIFIED:<br>DIMENSIONS ARE IN MILLIMETERS<br>SURFACE FINISH:<br>TOLERANCES:<br>LINEAR:<br>ANGULAR: | | | | FINISH: | | DEBURR AND BREAK SHARP EDGES | DO NOT SCALE DRAWING | REVISION |
|---|---|---|---|---|---|---|---|---|
| | NAME | SIGNATURE | DATE | | | | TITLE: | |
| DRAWN | | | | | | | | |
| CHK'D | | | | | | | | |
| APPV'D | | | | | | | | |
| MFG | | | | | | | | |
| Q.A | | | | MATERIAL: | | | DWG NO. AD-01 | A2 |
| | | | | WEIGHT: | | | SCALE:1:20 | SHEET 1 OF 1 |
