## Supplemental data, tables, and software for "Guide to the construction and use of an adaptive optics two-photon microscope with direct wavefront sensing": AD-02 z-motor unit.pdf

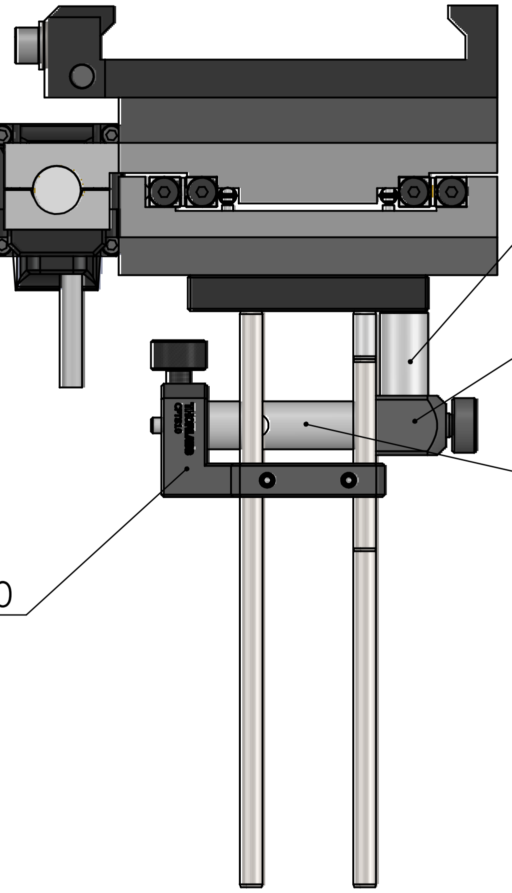


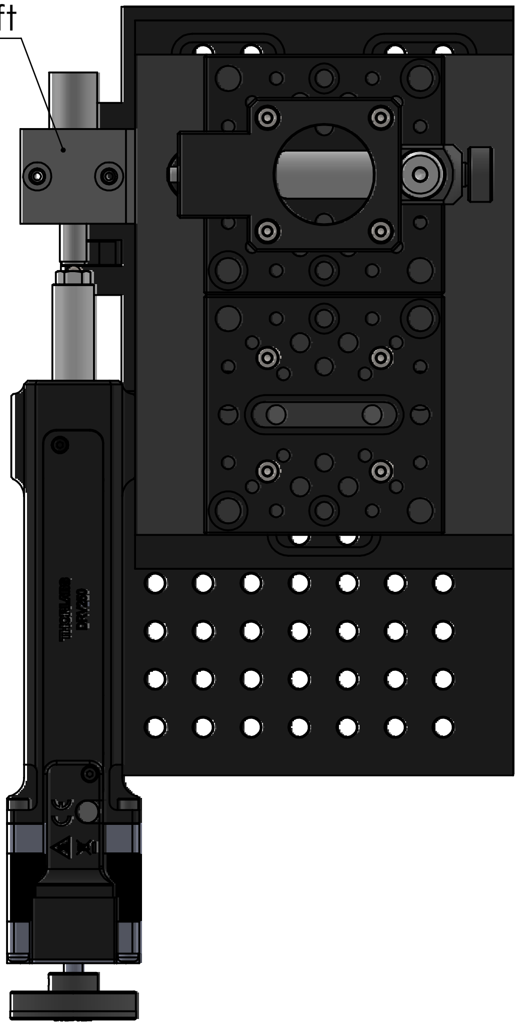


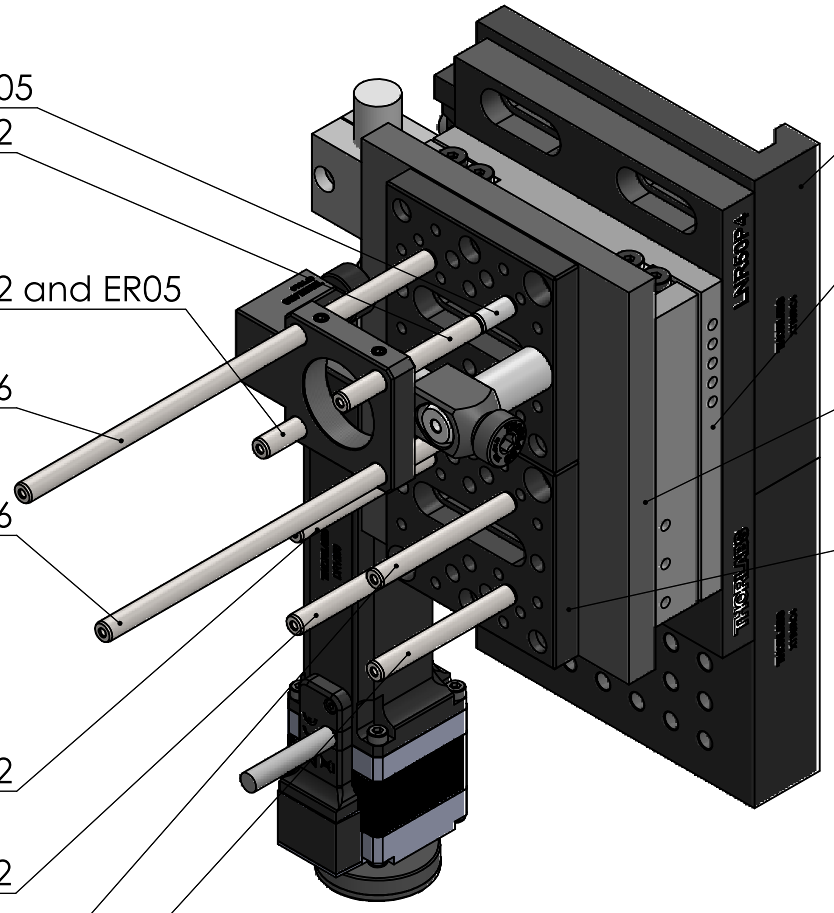


| | | | | | |
|---|---|---|---|---|---|
| UNLESS OTHERWISE SPECIFIED: DIMENSIONS ARE IN MILLIMETERS SURFACE FINISH: TOLERANCES: LINEAR: ANGULAR: | FINISH: | DEBURR AND BREAK SHARP EDGES | DO NOT SCALE DRAWING | REVISION | |
| | NAME | SIGNATURE | DATE | | TITLE: |
| DRAWN | | | | | |
| CHK'D | | | | | |
| APPV'D | | | | | |
| MFG | | | | | |
| Q.A | | | | MATERIAL: | DWG NO. AD-02 z-motor unit A3 |
| | | | | WEIGHT: | SCALE:1:5 SHEET 1 OF 1 |
