## Supplemental data, tables, and software for "Guide to the construction and use of an adaptive optics two-photon microscope with direct wavefront sensing": AD-11 scanner+tube lens.pdf

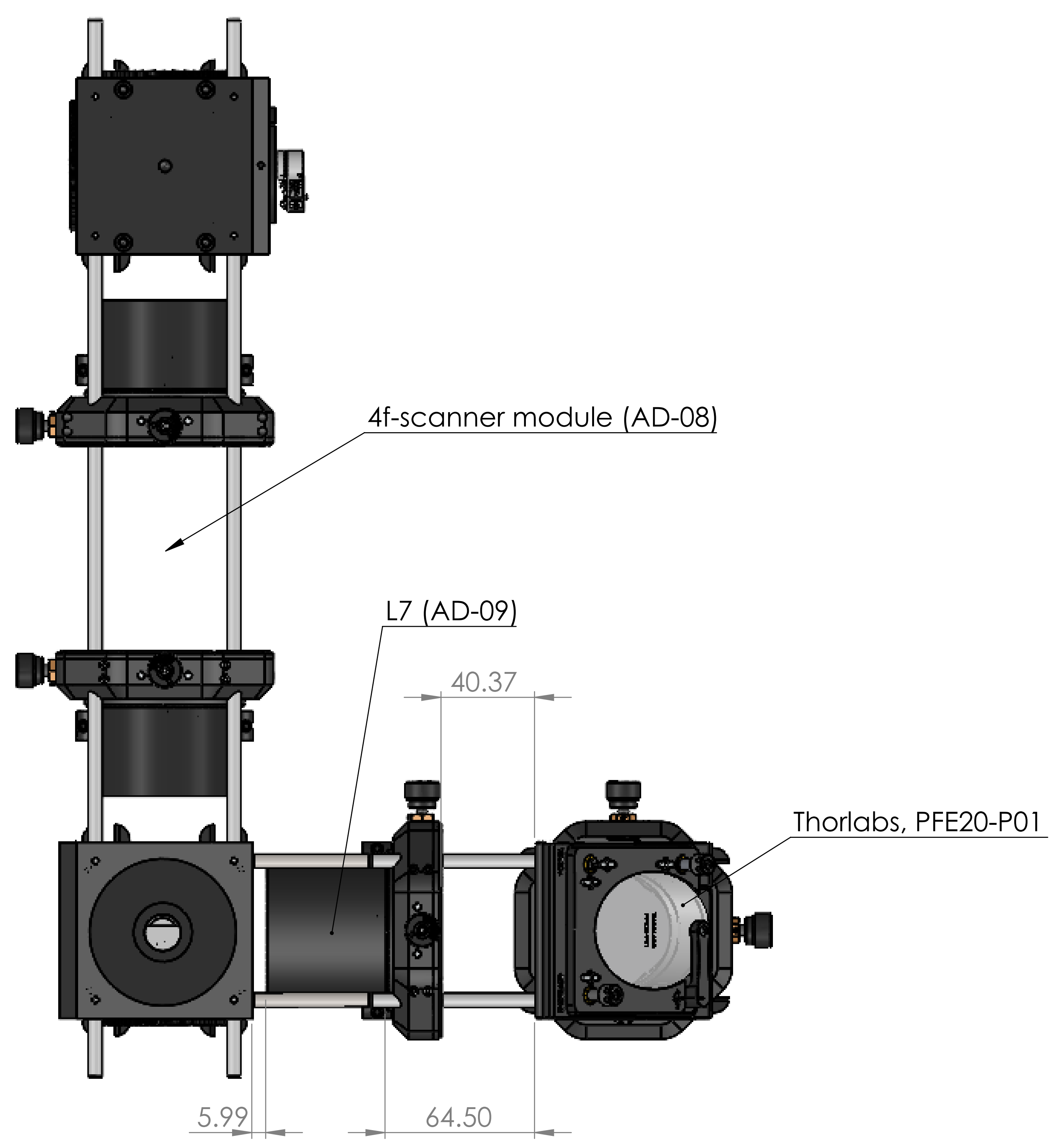


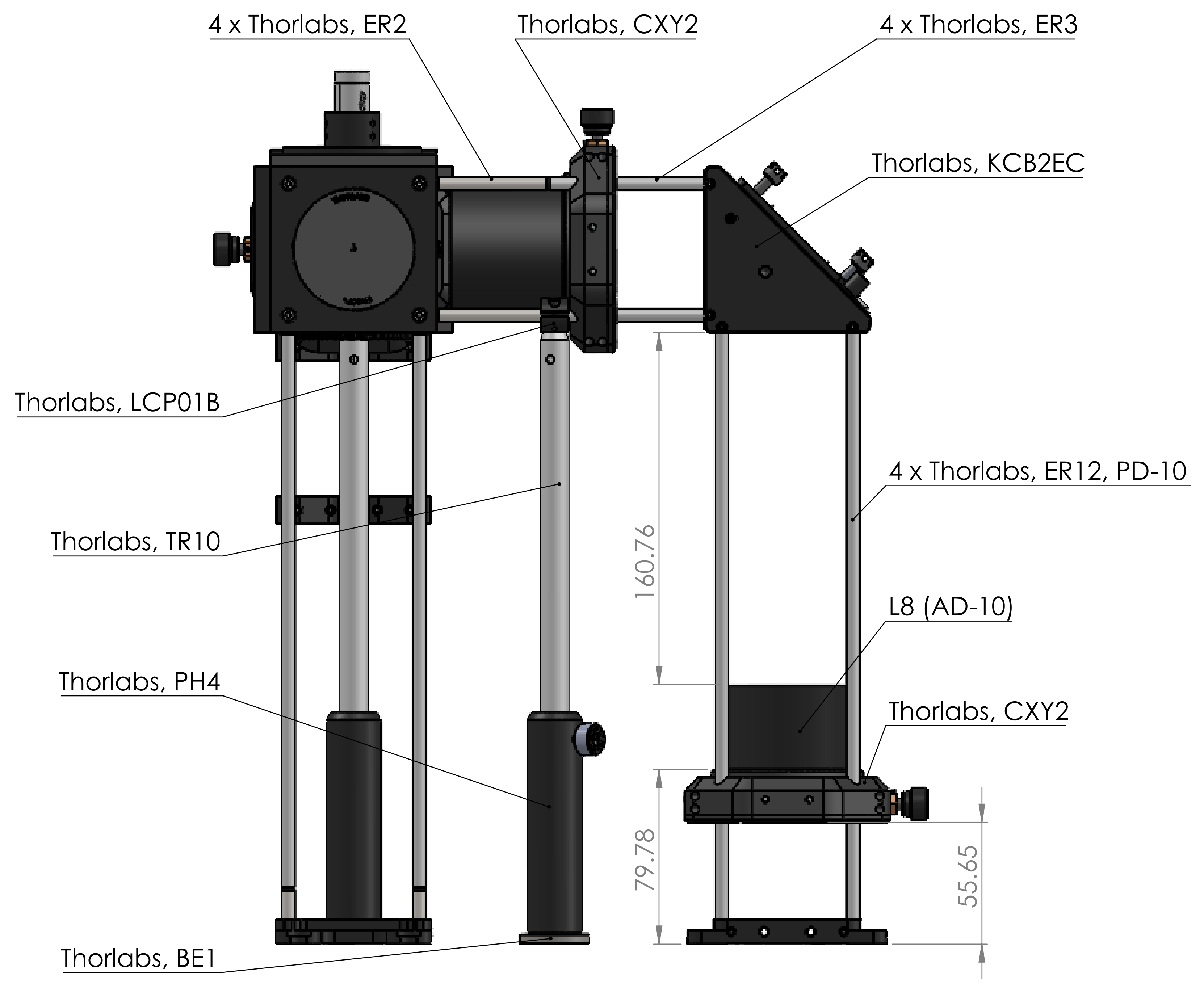


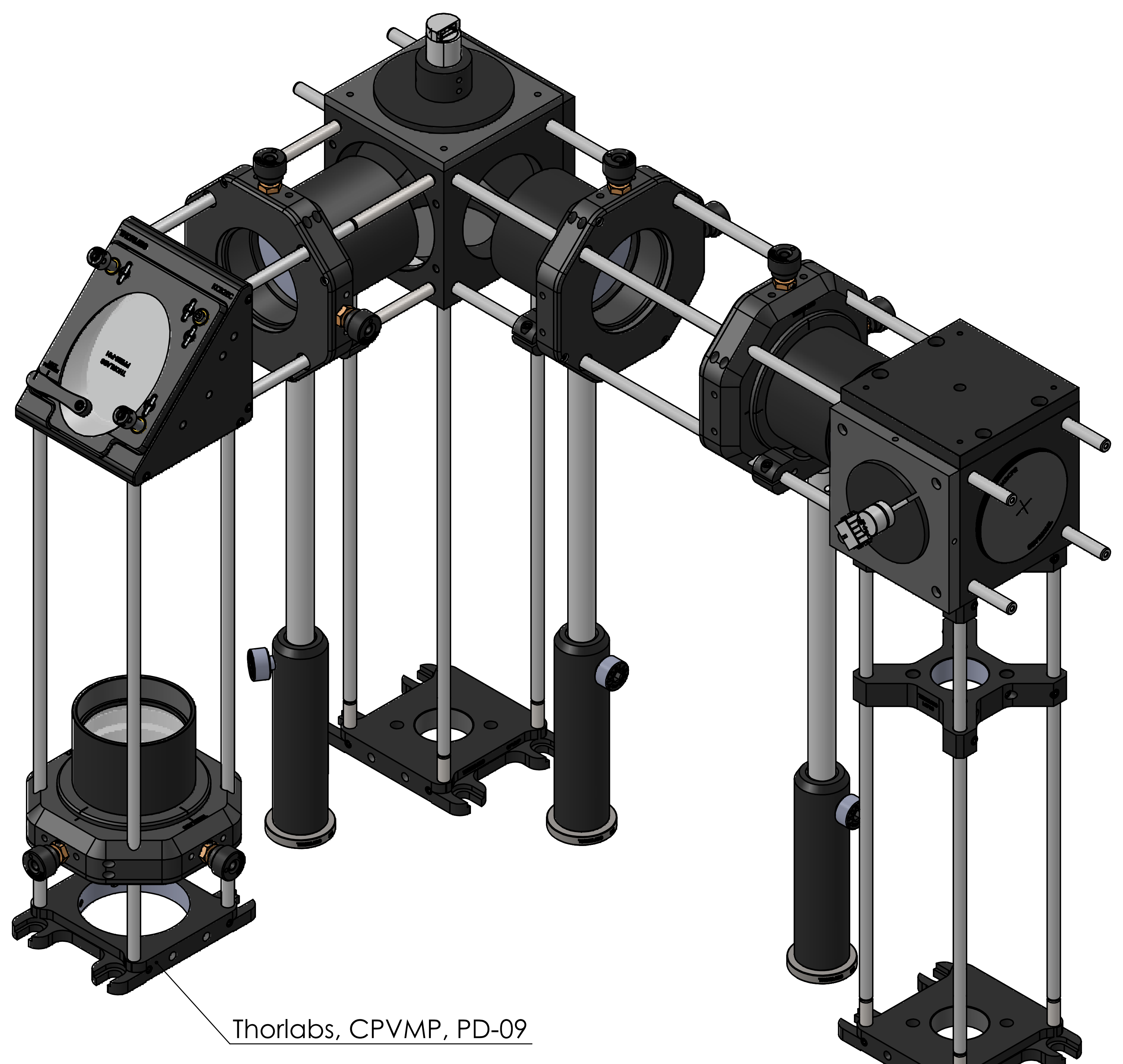


| UNLESS OTHERWISE SPECIFIED: DIMENSIONS ARE IN MILLIMETERS SURFACE FINISH: TOLERANCES: LINEAR: ANGULAR: | FINISH: | DEBURR AND BREAK SHARP EDGES | DO NOT SCALE DRAWING | REVISION |
|---|---|---|---|---|

| | NAME | SIGNATURE | DATE | | | TITLE: |
|---|---|---|---|---|---|---|
| DRAWN | | | | | | |
| CHK'D | | | | | | |
| APPV'D | | | | | | |
| MFG | | | | | | |
| Q.A | | | | MATERIAL: | | DWG NO. AD-11 scanner+tube lens |
| | | | | | | A2 |
| | | | | WEIGHT: | | SCALE:1:5 SHEET 1 OF 1 |
