## Supplemental data, tables, and software for "Guide to the construction and use of an adaptive optics two-photon microscope with direct wavefront sensing": AD-12 scanner module mount on main breadboard.pdf

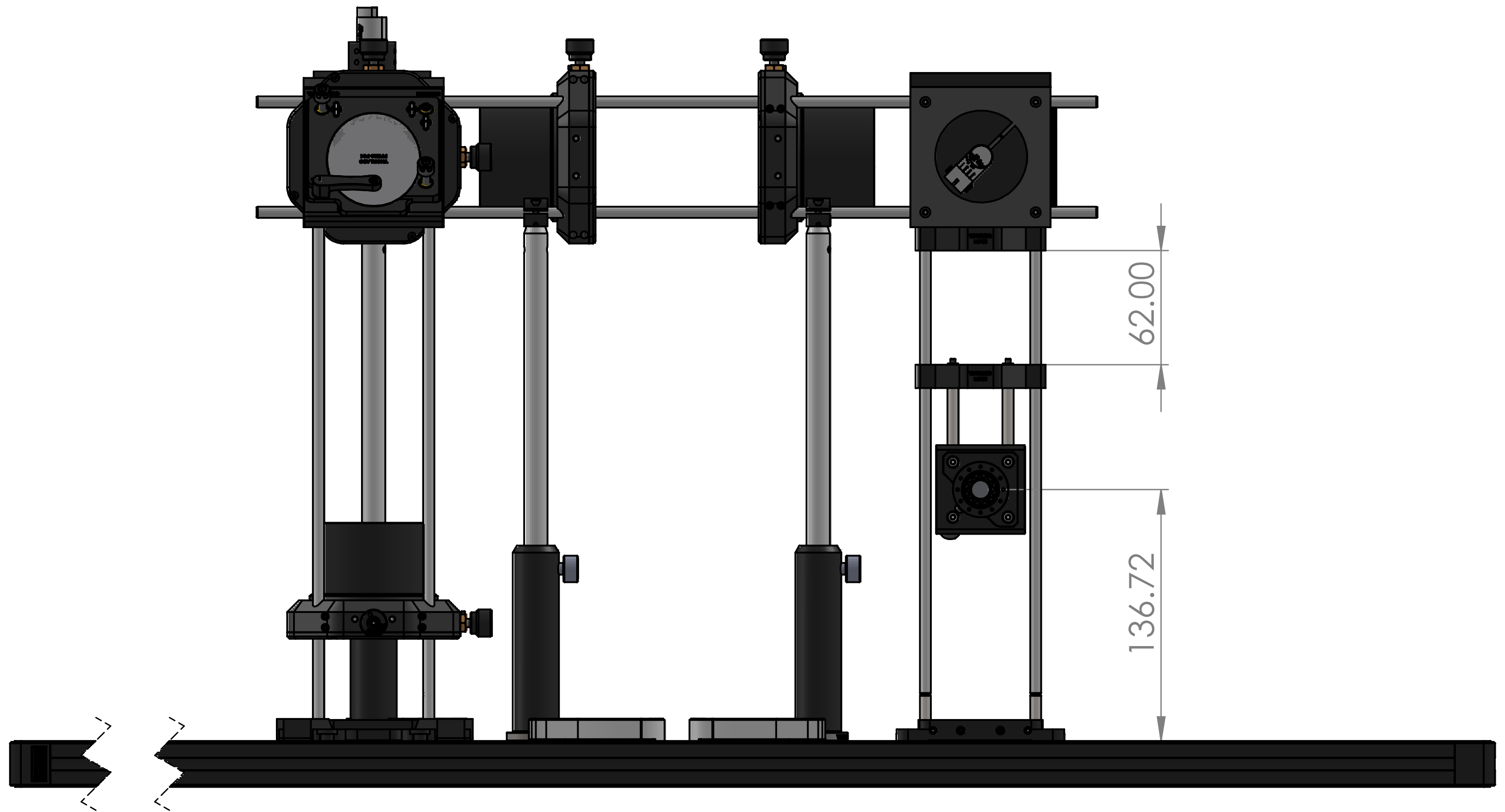


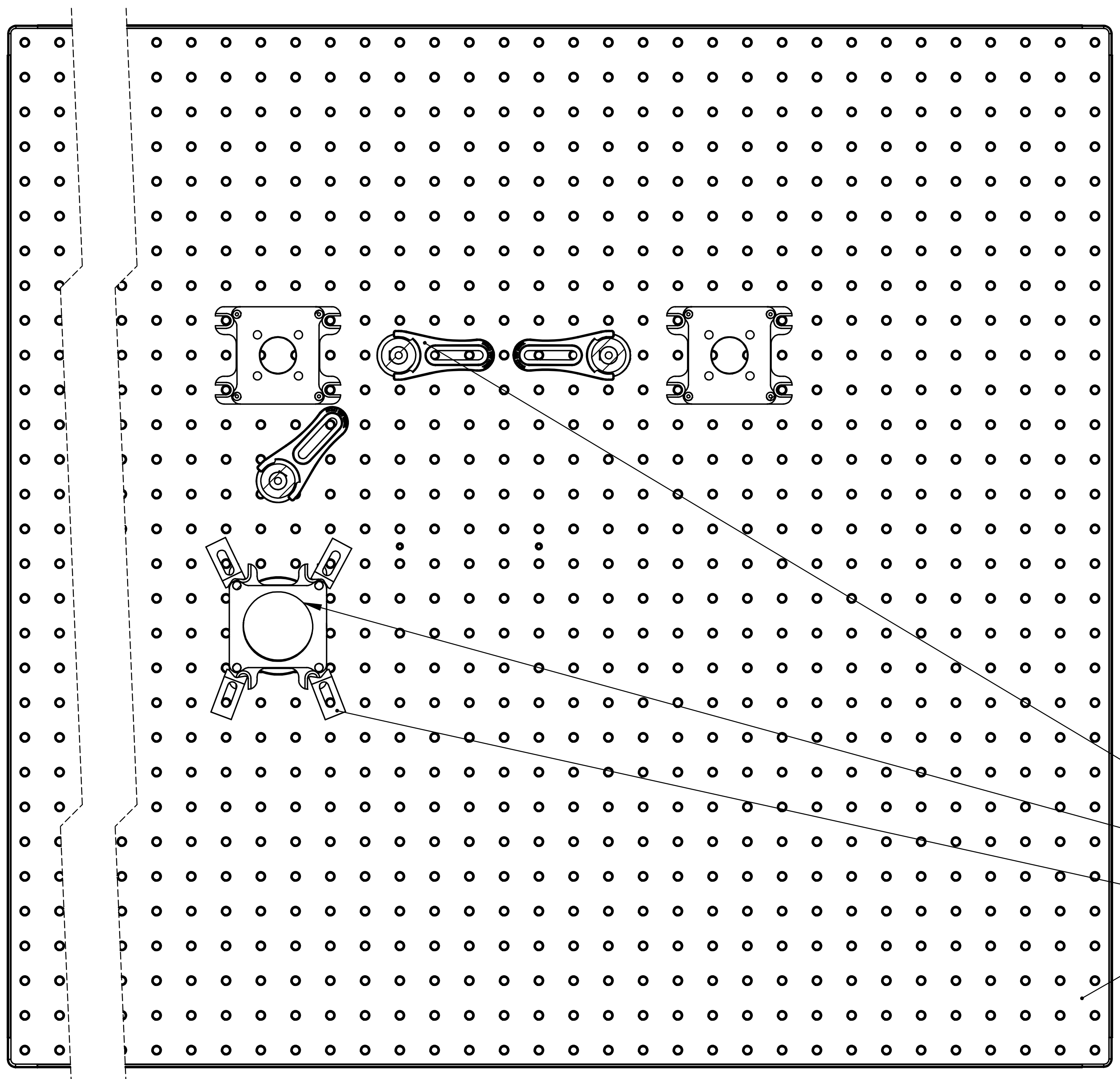


SECTION A-A
SCALE 1 : 4

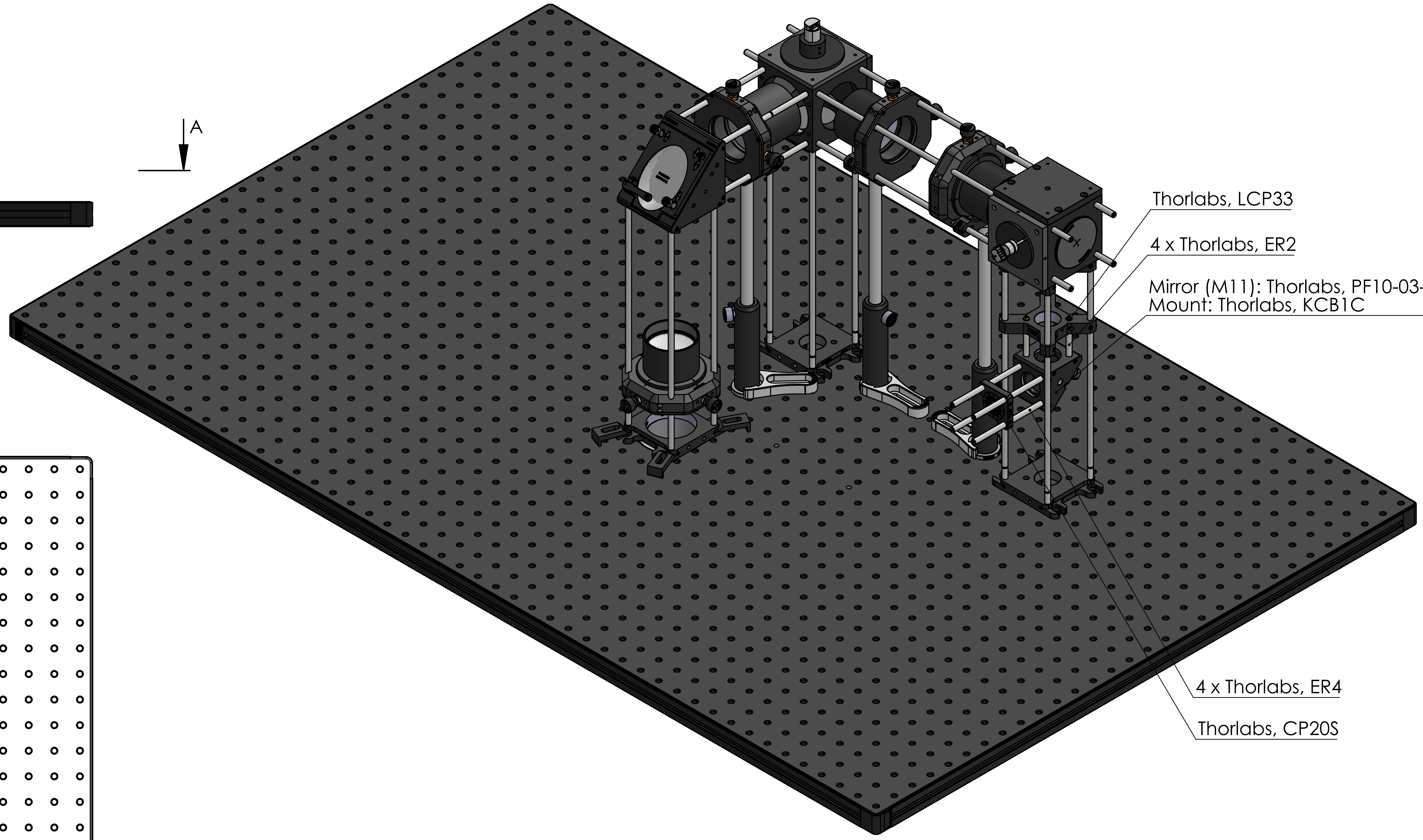


| | | | | | |
|---|---|---|---|---|---|
| UNLESS OTHERWISE SPECIFIED: DIMENSIONS ARE IN MILLIMETERS SURFACE FINISH: TOLERANCES: LINEAR: ANGULAR: | FINISH: | | DEBURR AND BREAK SHARP EDGES | DO NOT SCALE DRAWING | REVISION |
| NAME | SIGNATURE | DATE | | TITLE: | |
| DRAWN | | | | | |
| CHK'D | | | | | |
| APPV'D | | | | | |
| MFG | | | | | |
| Q.A | | | MATERIAL: | DWG NO. AD-12 scanner module mount on main b | A2 |
| | | | WEIGHT: | SCALE:1:10 | SHEET 1 OF 1 |
