## Supplemental data, tables, and software for "Guide to the construction and use of an adaptive optics two-photon microscope with direct wavefront sensing": AD-16 L1.pdf

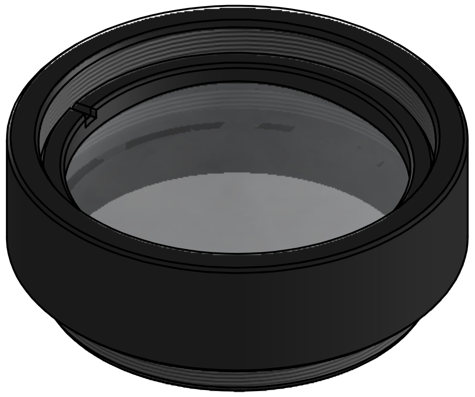

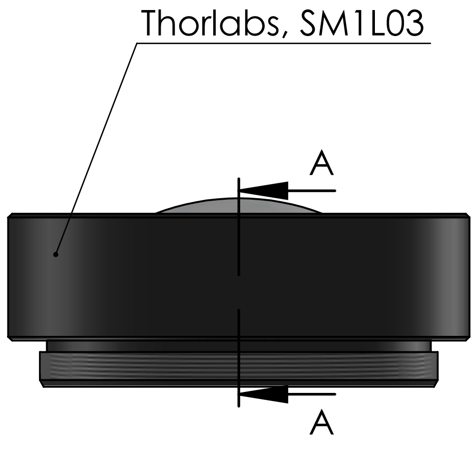


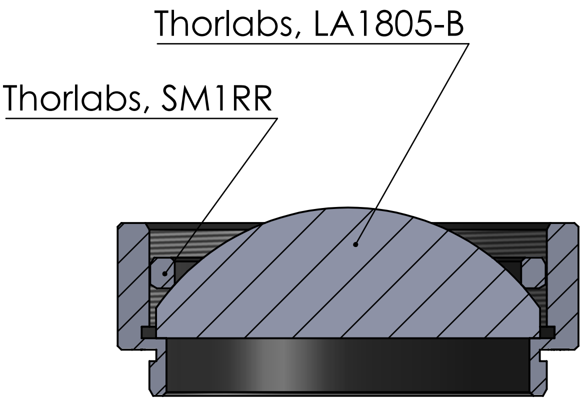


SECTION A-A

| | | | | | | |
|---|---|---|---|---|---|---|
| UNLESS OTHERWISE SPECIFIED:<br>DIMENSIONS ARE IN MILLIMETERS<br>SURFACE FINISH:<br>TOLERANCES:<br>LINEAR:<br>ANGULAR: | FINISH: | | | DEBURR AND BREAK SHARP EDGES | DO NOT SCALE DRAWING | REVISION |
| | NAME | SIGNATURE | DATE | | TITLE: | |
| DRAWN | | | | | | |
| CHK'D | | | | | | |
| APPV'D | | | | | | |
| MFG | | | | | | |
| Q.A | | | | MATERIAL: | DWG NO. AD-16 L1 | A4 |
| | | | | WEIGHT: | SCALE:2:1 | SHEET 1 OF 1 |
