## Supplemental data, tables, and software for "Guide to the construction and use of an adaptive optics two-photon microscope with direct wavefront sensing": AD-23 D2D3.pdf

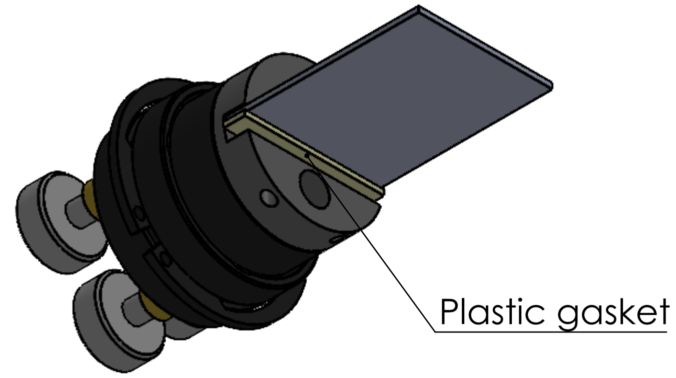


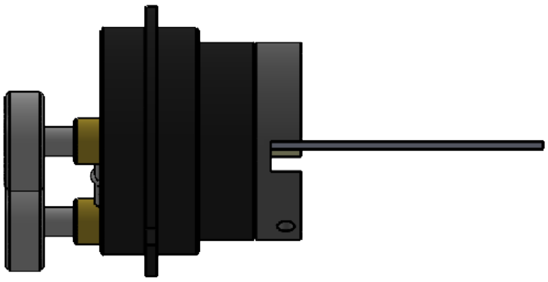

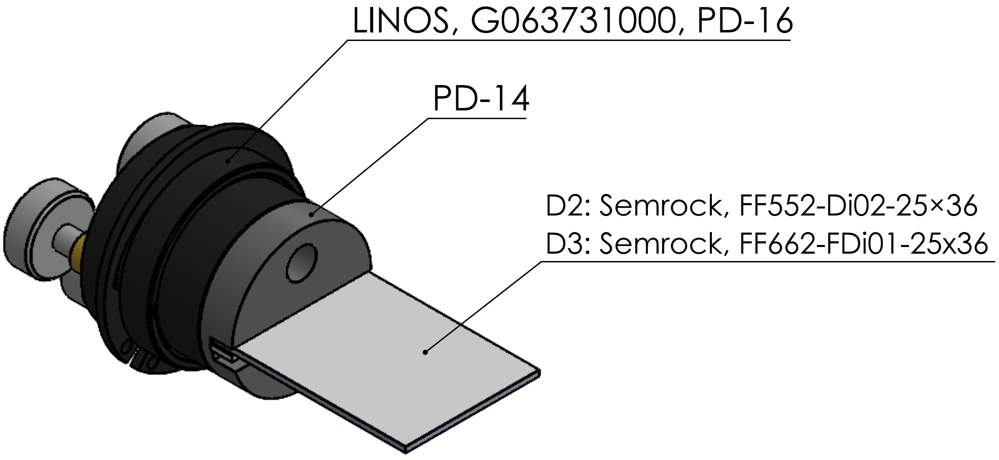


UNLESS OTHERWISE SPECIFIED:
DIMENSIONS ARE IN MILLIMETERS
SURFACE FINISH:
TOLERANCES:
LINEAR:
ANGULAR:

FINISH:

DEBURR AND BREAK SHARP EDGES

DO NOT SCALE DRAWING

REVISION

| | NAME | SIGNATURE | DATE |
|---|---|---|---|
| DRAWN | | | |
| CHK'D | | | |
| APPV'D | | | |
| MFG | | | |
| Q.A | | | |

TITLE:

MATERIAL:

WEIGHT:

DWG NO.

# AD-23 D2D3

A4

SCALE:1:2

SHEET 1 OF 1
