## Supplemental data, tables, and software for "Guide to the construction and use of an adaptive optics two-photon microscope with direct wavefront sensing": AD-26 L13L14.pdf

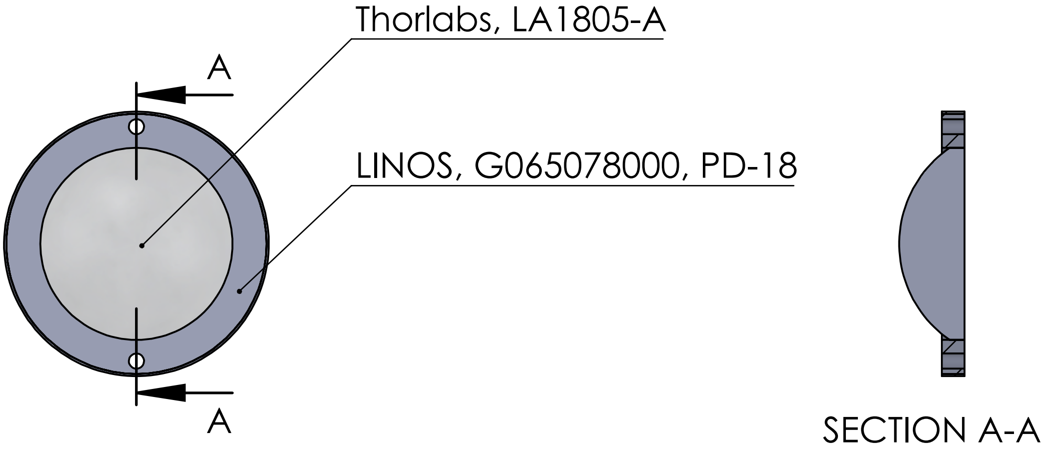


| | | | | | | | | |
|---|---|---|---|---|---|---|---|---|
| UNLESS OTHERWISE SPECIFIED: DIMENSIONS ARE IN MILLIMETERS SURFACE FINISH: TOLERANCES: LINEAR: ANGULAR: | | | FINISH: | | | DEBURR AND BREAK SHARP EDGES | DO NOT SCALE DRAWING | REVISION |
| | NAME | SIGNATURE | DATE | | | | TITLE: | |
| DRAWN | | | | | | | | |
| CHK'D | | | | | | | | |
| APPV'D | | | | | | | | |
| MFG | | | | | | | | |
| Q.A | | | | MATERIAL: | | | DWG NO. AD-26 L13L14 | A4 |
| | | | | WEIGHT: | | | SCALE:1:1 | SHEET 1 OF 1 |
