## Supplemental data, tables, and software for "Guide to the construction and use of an adaptive optics two-photon microscope with direct wavefront sensing": AD-37 SHWS (with stage).pdf

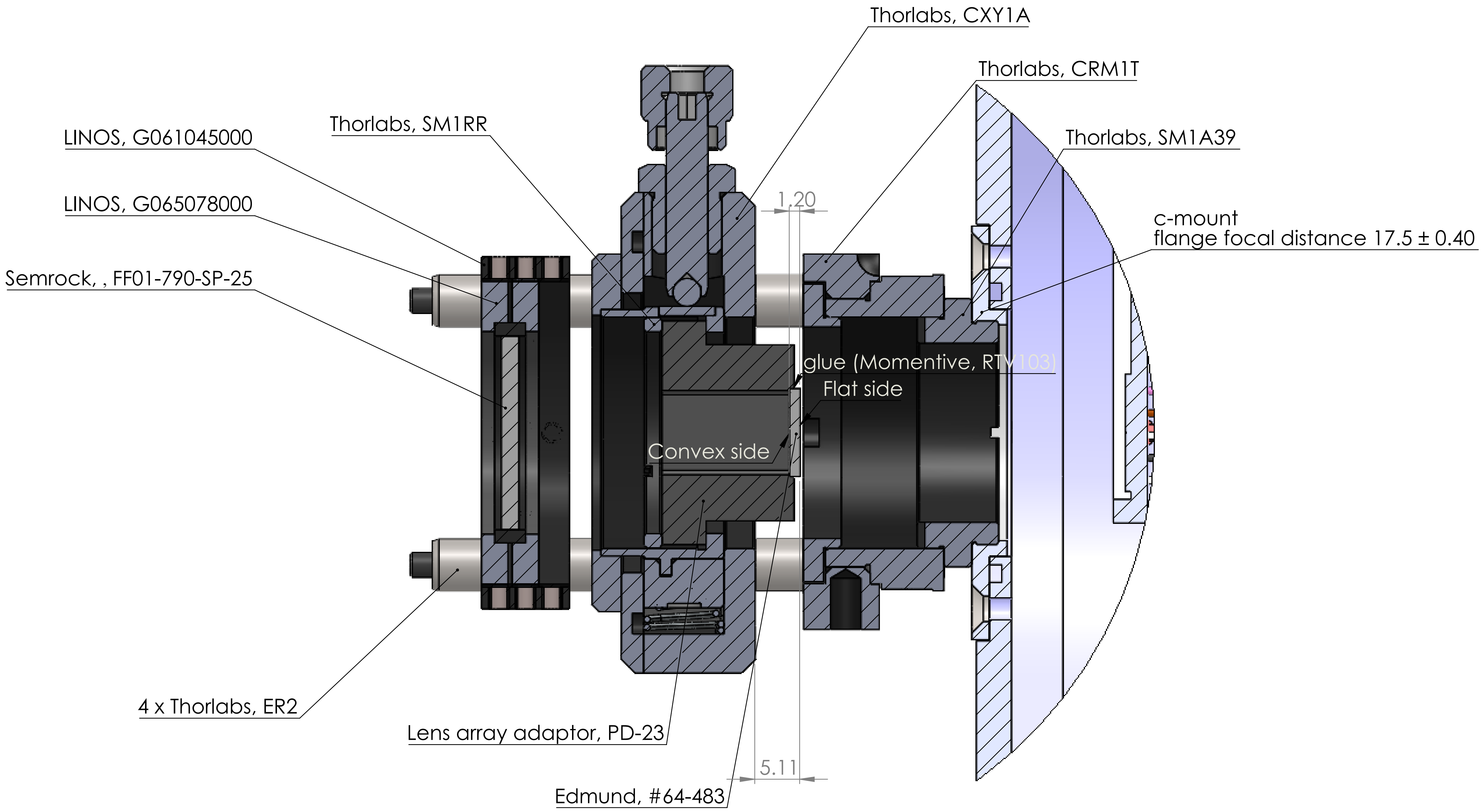


DETAIL B
SCALE 2 : 1

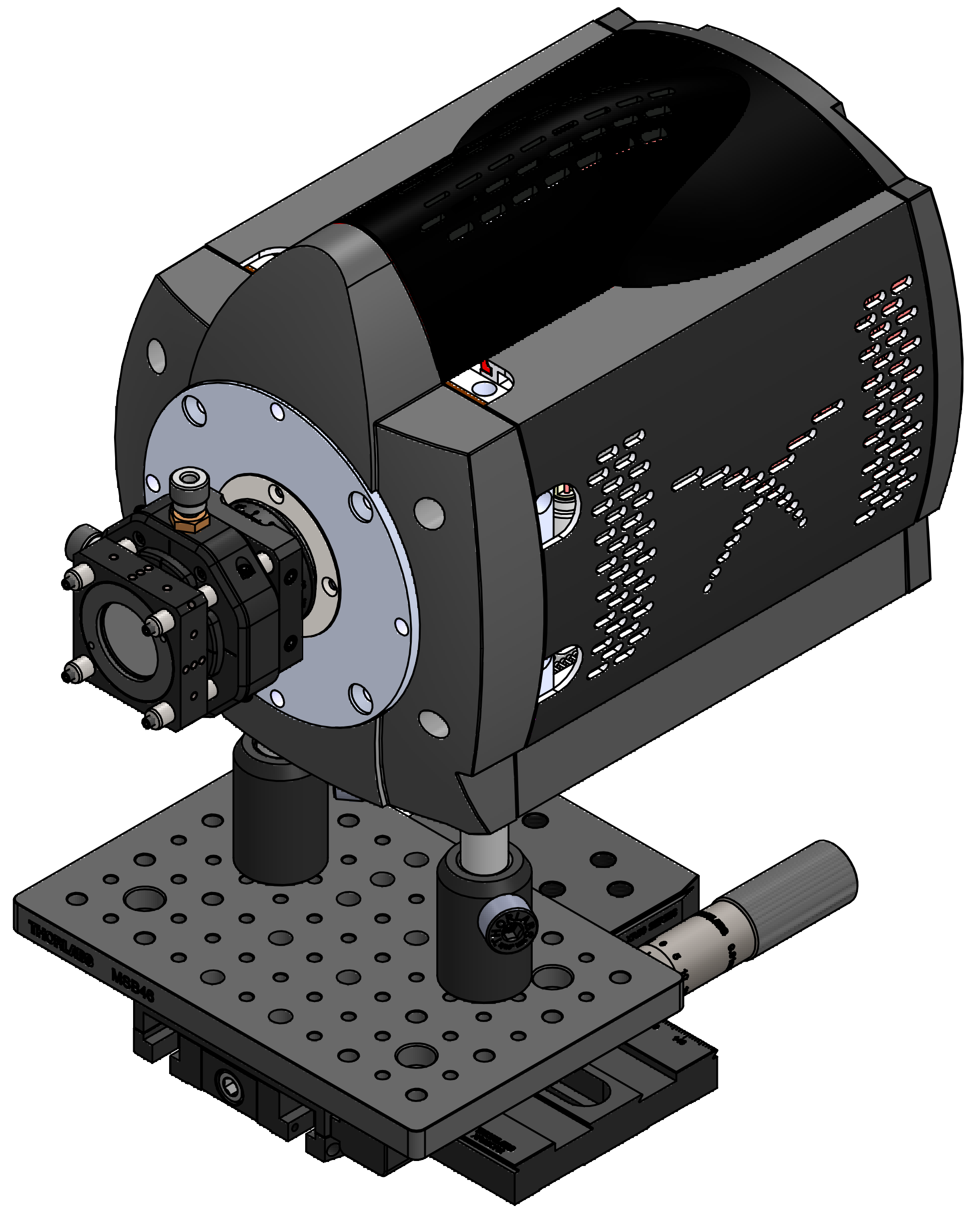

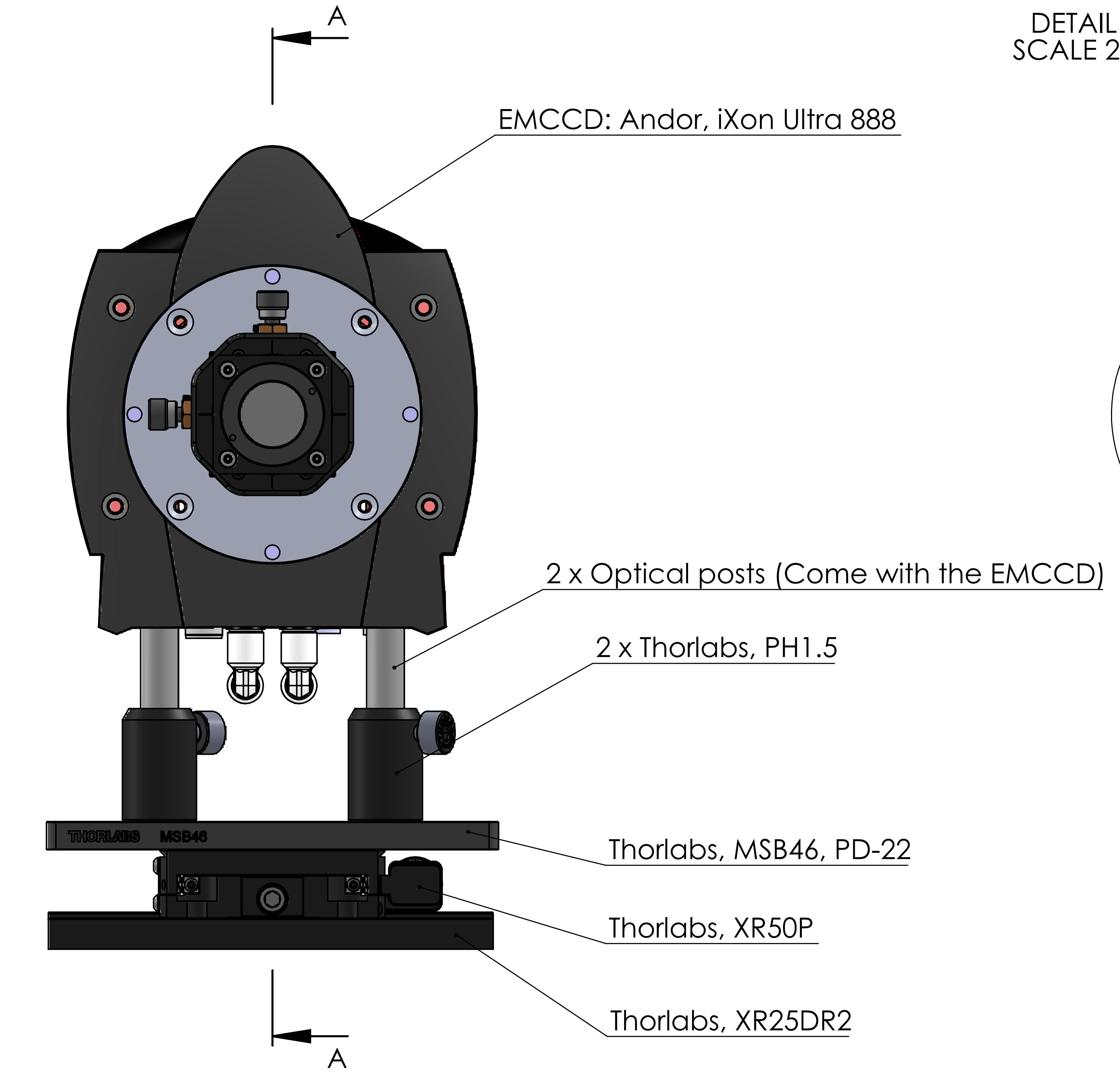


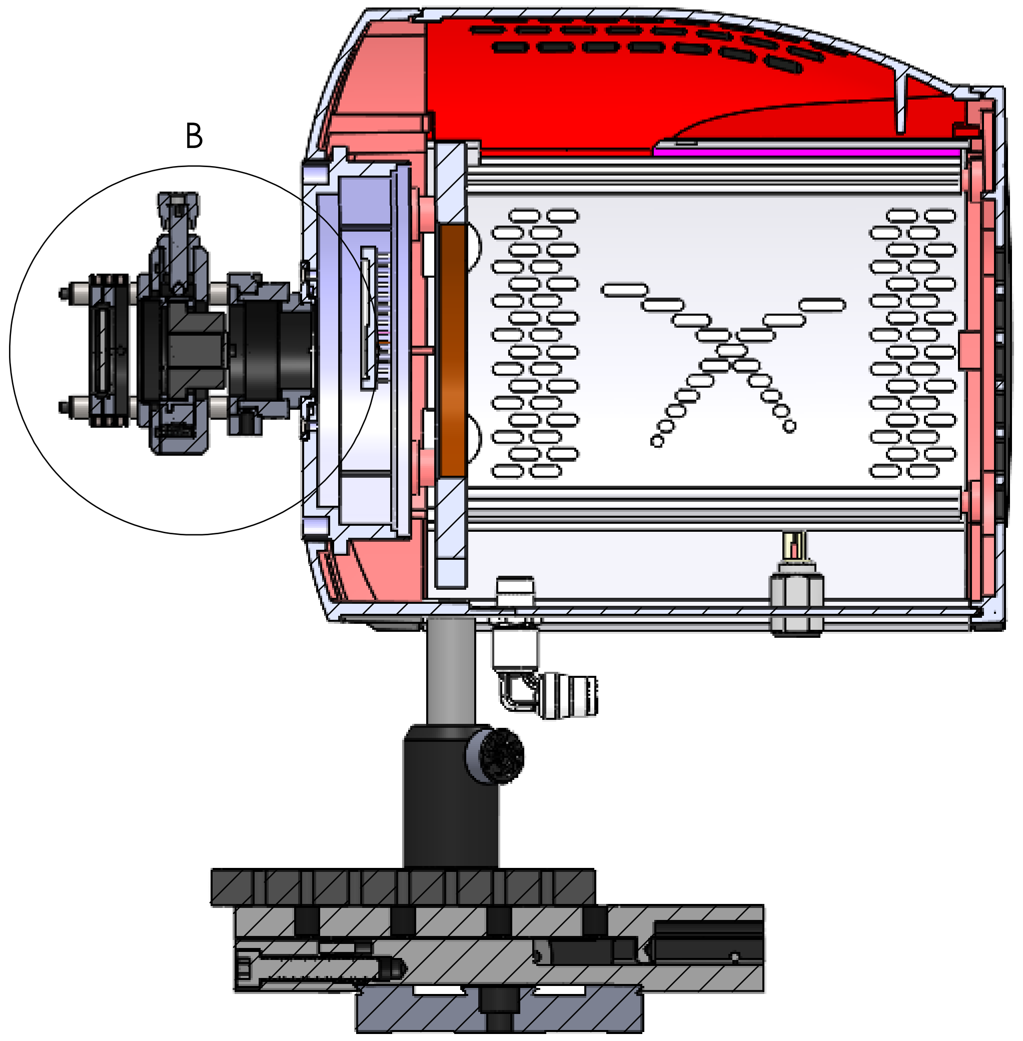


SECTION A-A
SCALE 1 : 2

| UNLESS OTHERWISE SPECIFIED: DIMENSIONS ARE IN MILLIMETERS SURFACE FINISH: TOLERANCES: LINEAR: ANGULAR: | FINISH: | DEBURR AND BREAK SHARP EDGES | DO NOT SCALE DRAWING | REVISION |
|---|---|---|---|---|
| | NAME | SIGNATURE | DATE | TITLE: |
| DRAWN | | | | |
| CHK'D | | | | |
| APPV'D | | | | |
| MFG | | | | |
| Q.A | | | MATERIAL: | DWG NO. AD-37 SHWS (with stage) A2 |
| | | | WEIGHT: | SCALE:1:5 SHEET 1 OF 1 |
