## Supplemental data, tables, and software for "Guide to the construction and use of an adaptive optics two-photon microscope with direct wavefront sensing": AD-38 SHWS on breadboard.pdf

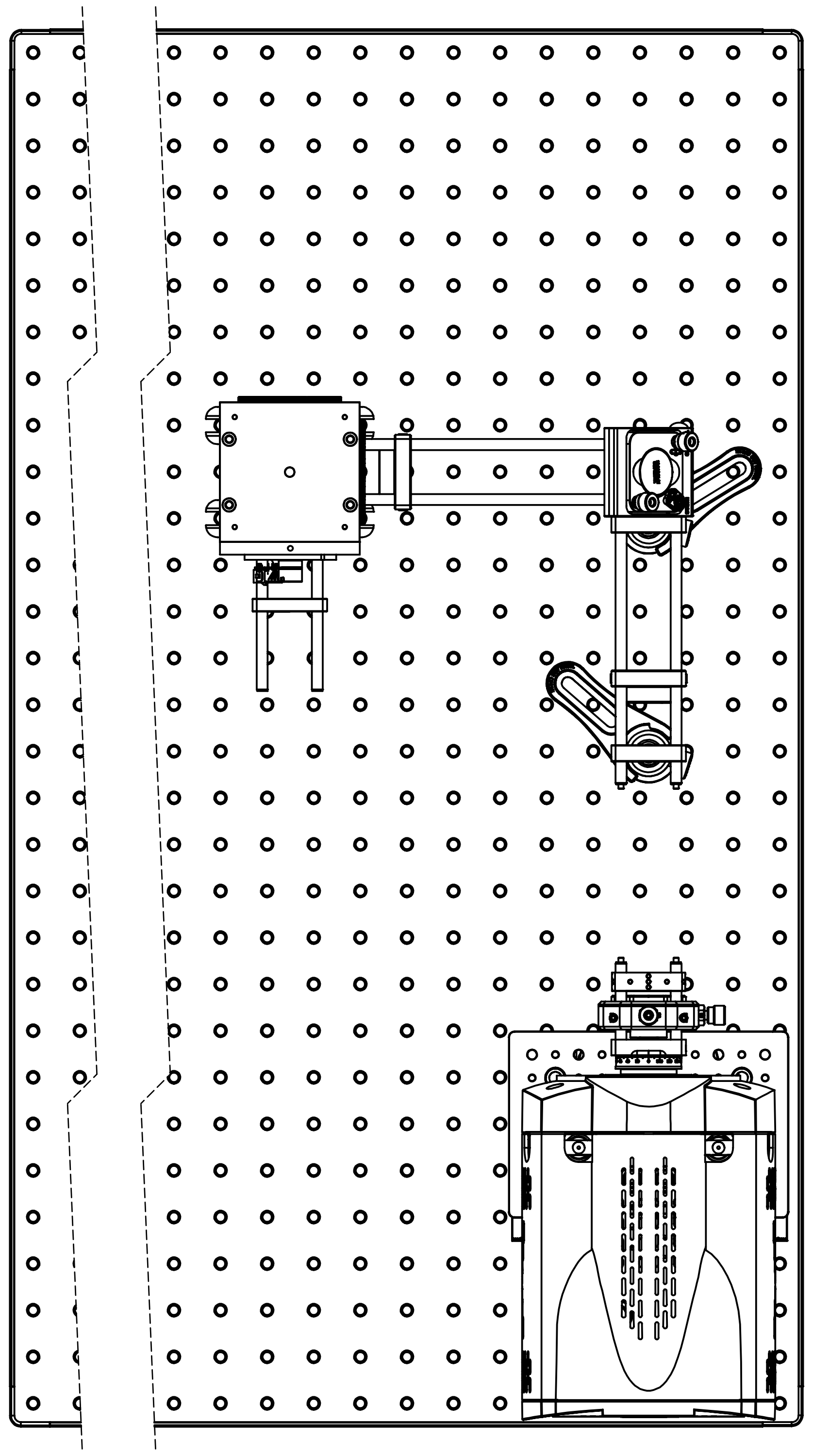

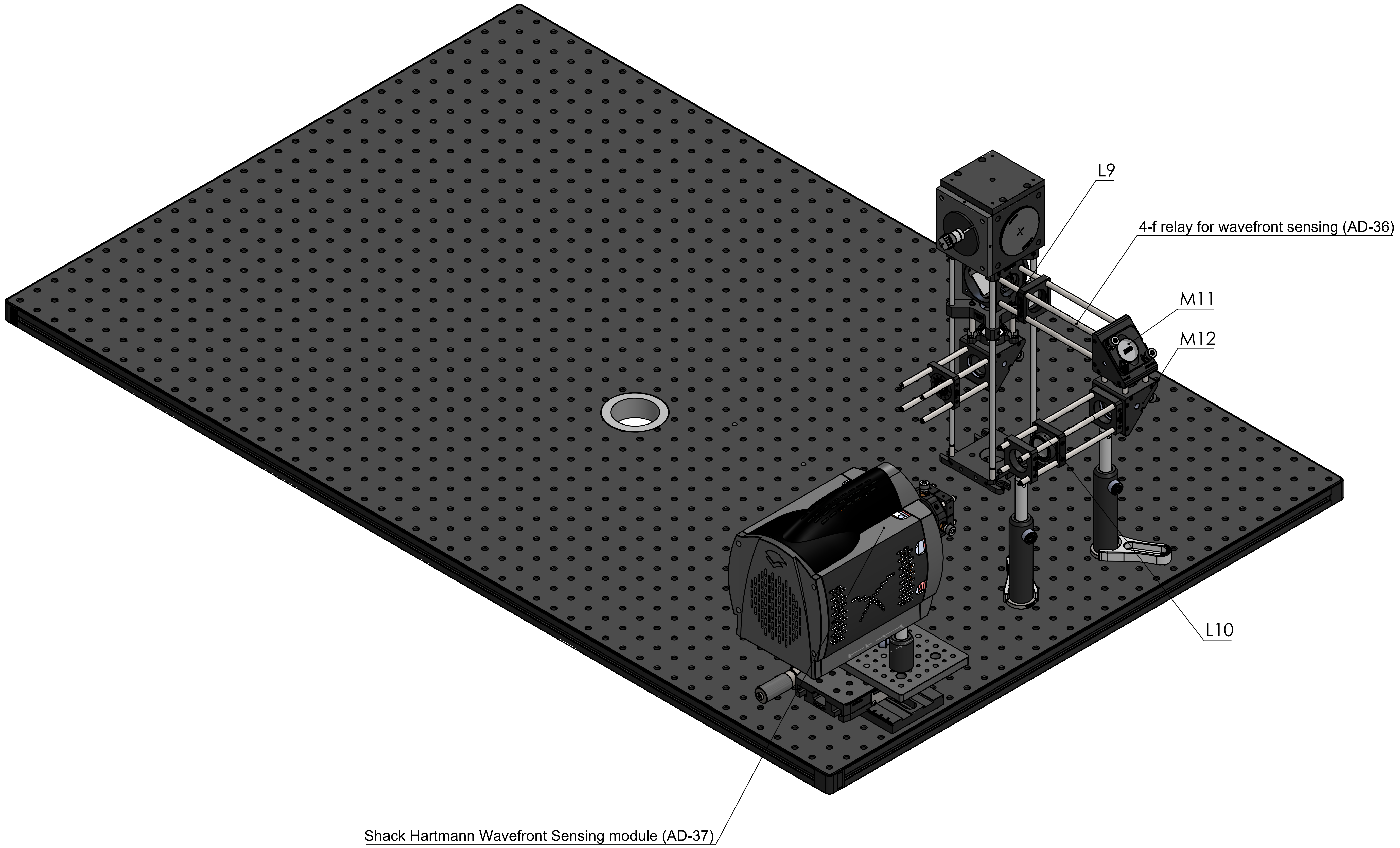


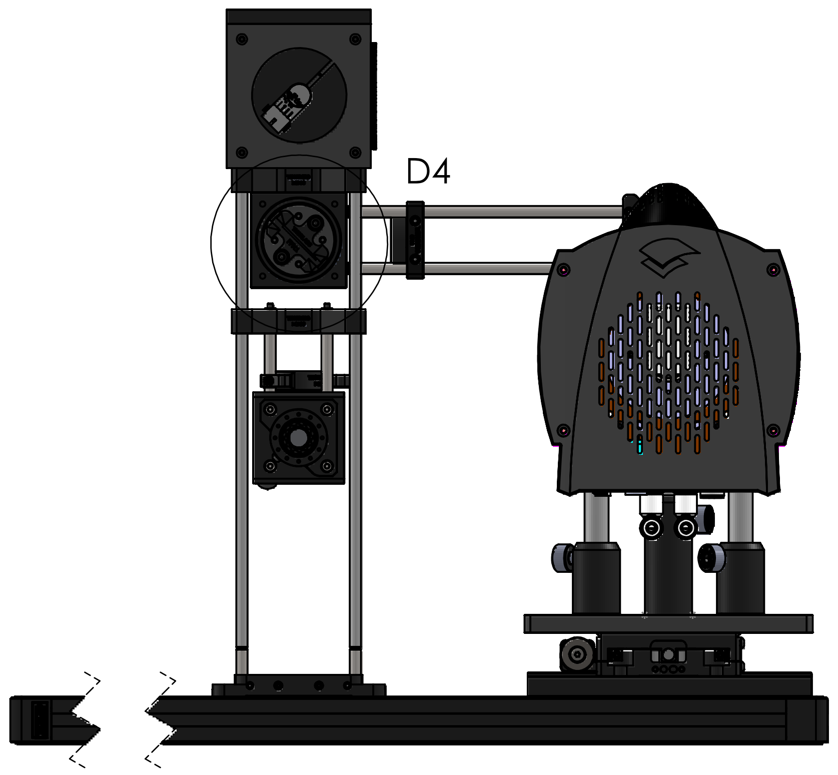


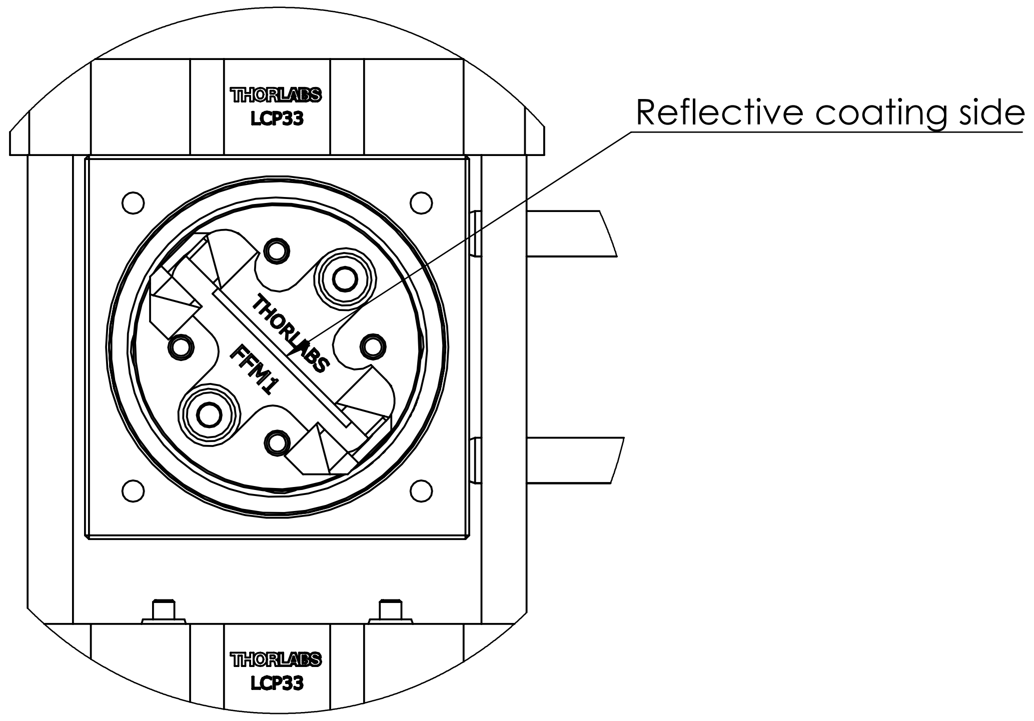


DETAIL D4
SCALE 1 : 1

| | | | | | | | |
|---|---|---|---|---|---|---|---|
| UNLESS OTHERWISE SPECIFIED: DIMENSIONS ARE IN MILLIMETERS SURFACE FINISH: TOLERANCES: LINEAR: ANGULAR: | FINISH: | | | | DEBURR AND BREAK SHARP EDGES | DO NOT SCALE DRAWING | REVISION |
| | NAME | SIGNATURE | DATE | | | TITLE: | |
| DRAWN | | | | | | | |
| CHK'D | | | | | | | |
| APPV'D | | | | | | | |
| MFG | | | | | | | |
| Q.A | | | | MATERIAL: | | DWG NO. AD-38 SHWS on breadboar[illegible] | A2 |
| | | | | WEIGHT: | | SCALE:1:10 | SHEET 1 OF 1 |
