## Supplemental data, tables, and software for "Guide to the construction and use of an adaptive optics two-photon microscope with direct wavefront sensing": AD-40 camera for DM conjugation.pdf

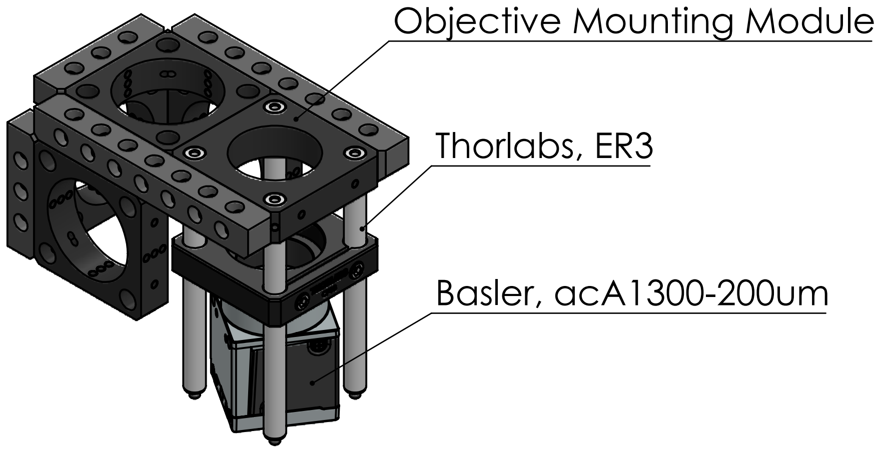

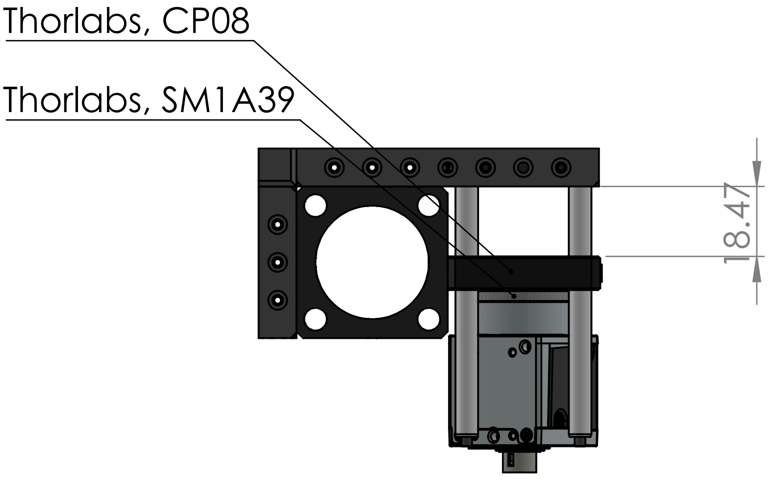

| UNLESS OTHERWISE SPECIFIED:<br>DIMENSIONS ARE IN MILLIMETERS<br>SURFACE FINISH:<br>TOLERANCES:<br>LINEAR:<br>ANGULAR: | FINISH: | DEBURR AND<br>BREAK SHARP<br>EDGES | DO NOT SCALE DRAWING | REVISION |
|---|---|---|---|---|

| | NAME | SIGNATURE | DATE |
|---|---|---|---|
| DRAWN | | | |
| CHK'D | | | |
| APPV'D | | | |
| MFG | | | |
| Q.A | | | |

TITLE:

MATERIAL:

DWG NO.

AD-40 camera for DM conjuga[illegible]

A4

WEIGHT:

SCALE:1:2

SHEET 1 OF 1
