## Supplemental data, tables, and software for "Guide to the construction and use of an adaptive optics two-photon microscope with direct wavefront sensing": AD-41 deformable mirror conjugation tool.pdf

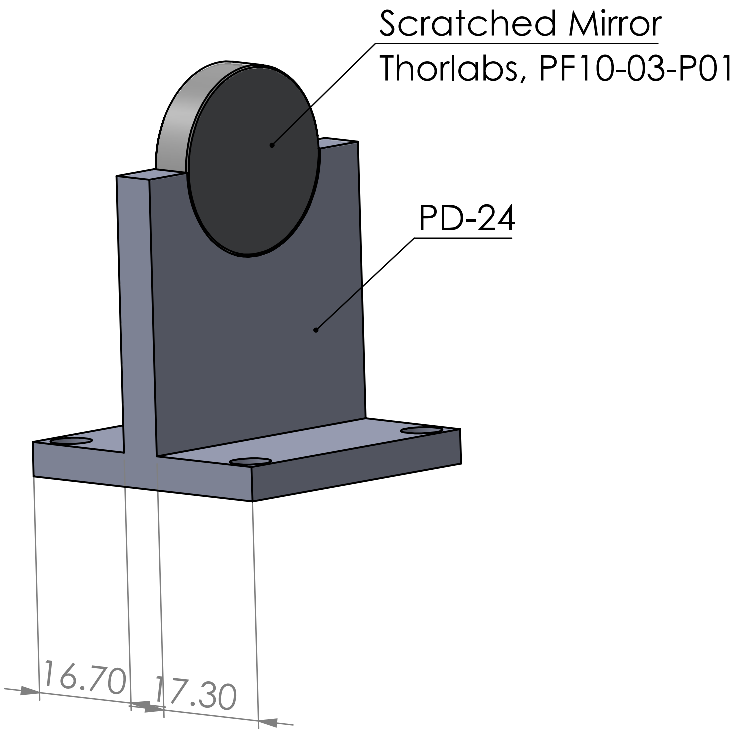


| | | | | |
|---|---|---|---|---|
| UNLESS OTHERWISE SPECIFIED: DIMENSIONS ARE IN MILLIMETERS SURFACE FINISH: TOLERANCES: LINEAR: ANGULAR: | FINISH: | DEBURR AND BREAK SHARP EDGES | DO NOT SCALE DRAWING | REVISION |

| | NAME | SIGNATURE | DATE | | | |
|---|---|---|---|---|---|---|
| DRAWN | | | | | | TITLE: |
| CHK'D | | | | | | |
| APPV'D | | | | | | |
| MFG | | | | | | |
| Q.A | | | | MATERIAL: | | DWG NO. A4 |
| | | | | WEIGHT: | | SCALE:1:1 SHEET 1 OF 1 |

deformable mirror conjugation
