## Supplemental data, tables, and software for "Guide to the construction and use of an adaptive optics two-photon microscope with direct wavefront sensing": PD-01 main breadboard.pdf

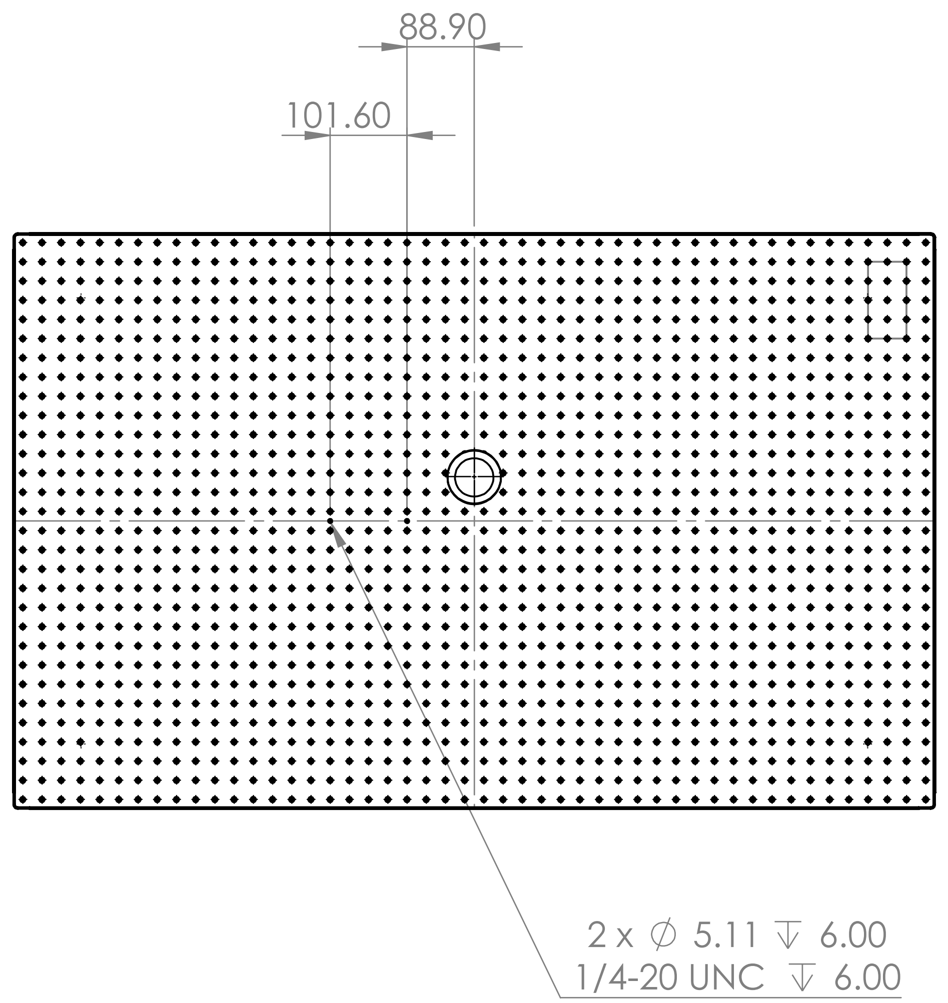


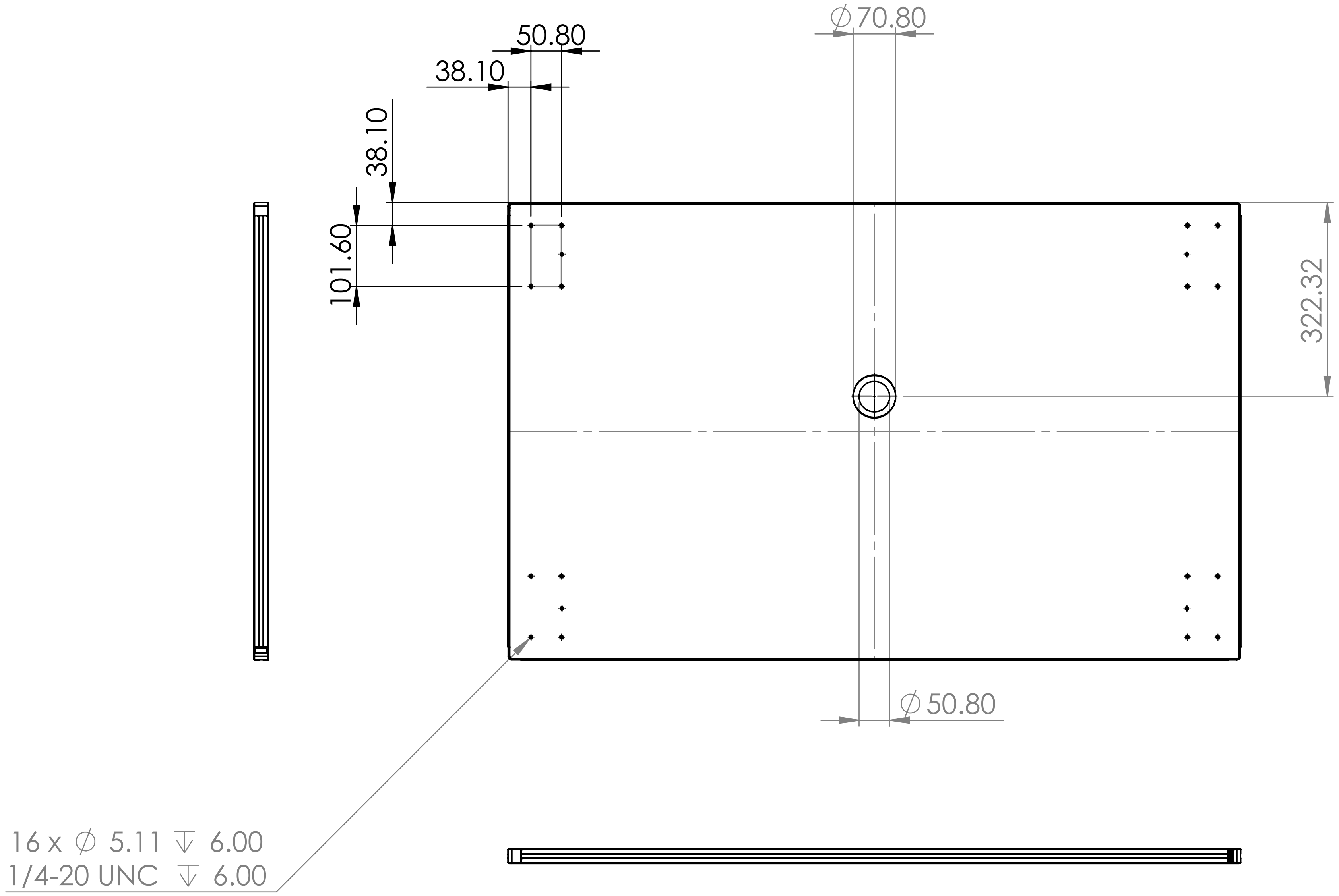


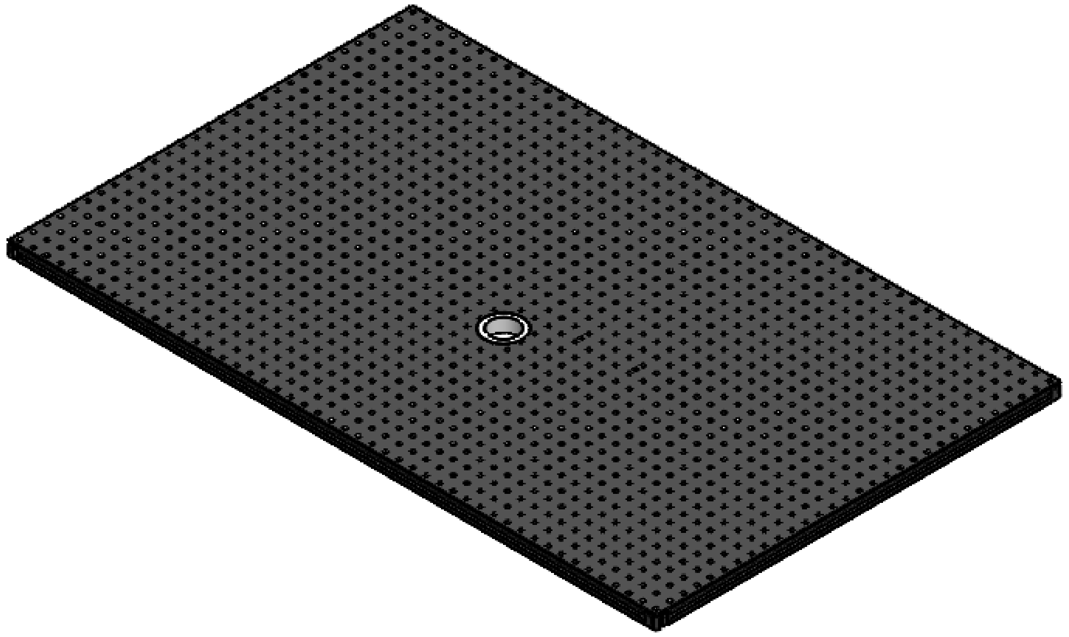

Modified from Thorlabs, PBG3048F

16 tapped holes at corners (back);

2 tapped holes at midline(front)

A center hole of 50.8 mm inner diameter with 10 mm thickness wall

| | | | | | |
|---|---|---|---|---|---|
| UNLESS OTHERWISE SPECIFIED: DIMENSIONS ARE IN MILLIMETERS SURFACE FINISH: TOLERANCES: LINEAR: ANGULAR: | FINISH: | | DEBURR AND BREAK SHARP EDGES | DO NOT SCALE DRAWING | REVISION |
| | | | | Quantity: 1 | |
| | NAME | SIGNATURE | DATE | TITLE: Main Breadboard | |
| DRAWN | | | | | |
| CHK'D | | | | | |
| APPV'D | | | | | |
| MFG | | | | | |
| Q.A | | | MATERIAL: 6082-T6 ALUMINIUM Anodized Aluminum | DWG NO. Part Drawing 01 | A3 |
| | | | WEIGHT: | SCALE:1:10 | SHEET 1 OF 1 |
