## Supplemental data, tables, and software for "Guide to the construction and use of an adaptive optics two-photon microscope with direct wavefront sensing": PD-02 z-motor adaptor.pdf

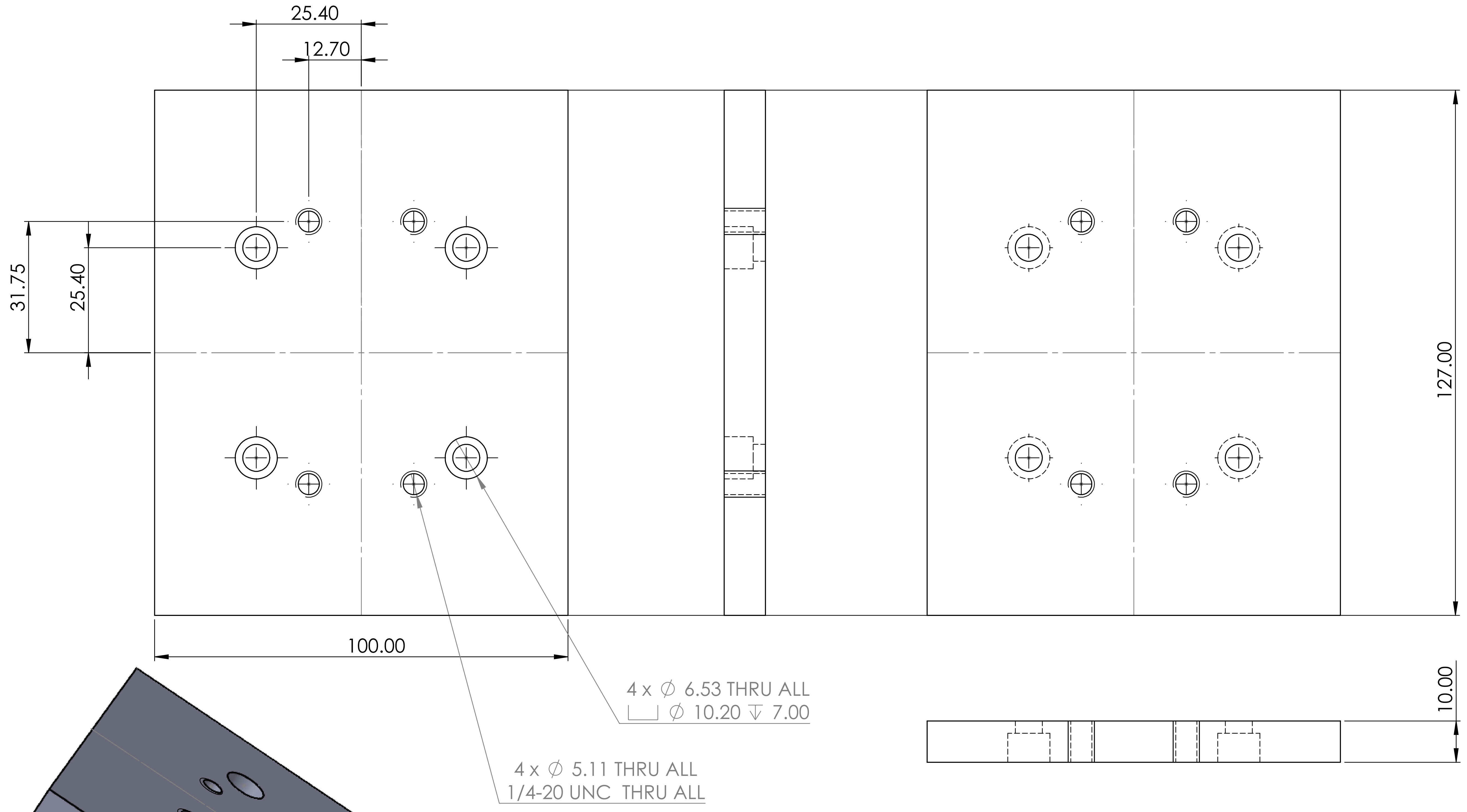

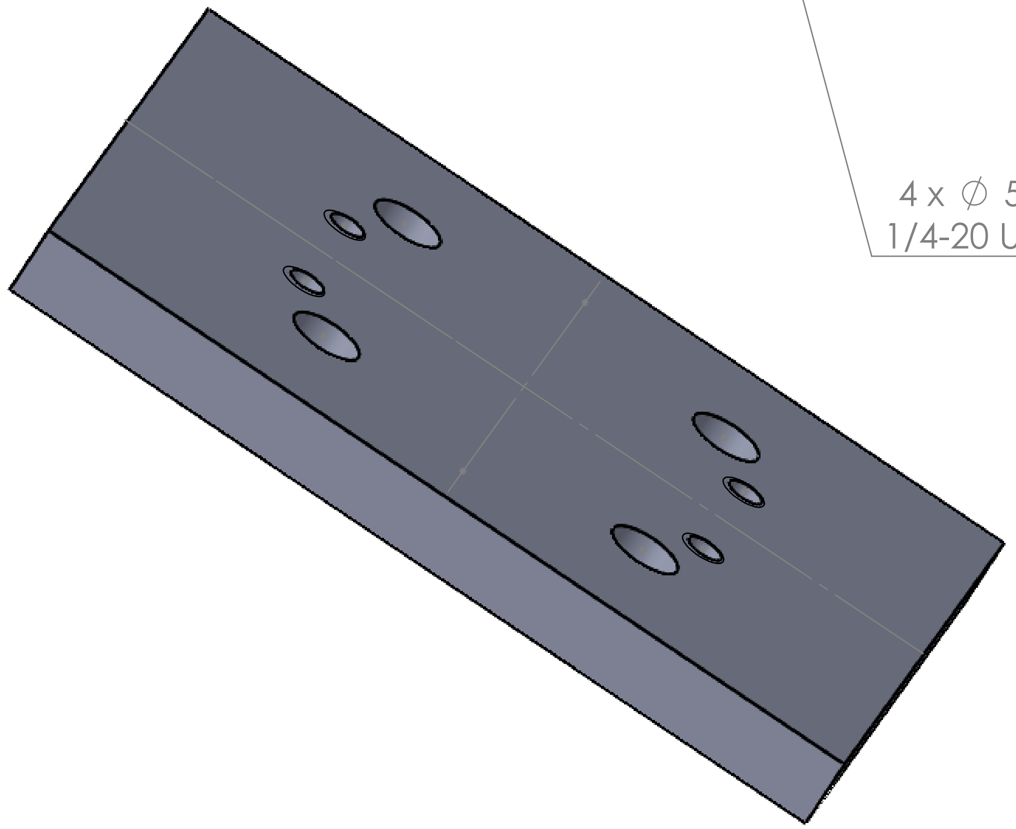

| | | | | | |
|---|---|---|---|---|---|
| UNLESS OTHERWISE SPECIFIED:<br>DIMENSIONS ARE IN MILLIMETERS<br>SURFACE FINISH:<br>TOLERANCES:<br>LINEAR:<br>ANGULAR: | FINISH: | | DEBURR AND BREAK SHARP EDGES | DO NOT SCALE DRAWING | REVISION |
| | | | | Quantity: 1 | |
| | NAME | SIGNATURE | DATE | TITLE: | |
| DRAWN | | | | Z-motor Adaptor | |
| CHK'D | | | | | |
| APPV'D | | | | | |
| MFG | | | | | |
| Q.A | | | MATERIAL: Anodized Aluminum | DWG NO. Part Drawing 02 | A3 |
| | | | WEIGHT: | SCALE:1:1 | SHEET 1 OF 1 |
