## Supplemental data, tables, and software for "Guide to the construction and use of an adaptive optics two-photon microscope with direct wavefront sensing": PD-04 Galvo mounting plate.PDF

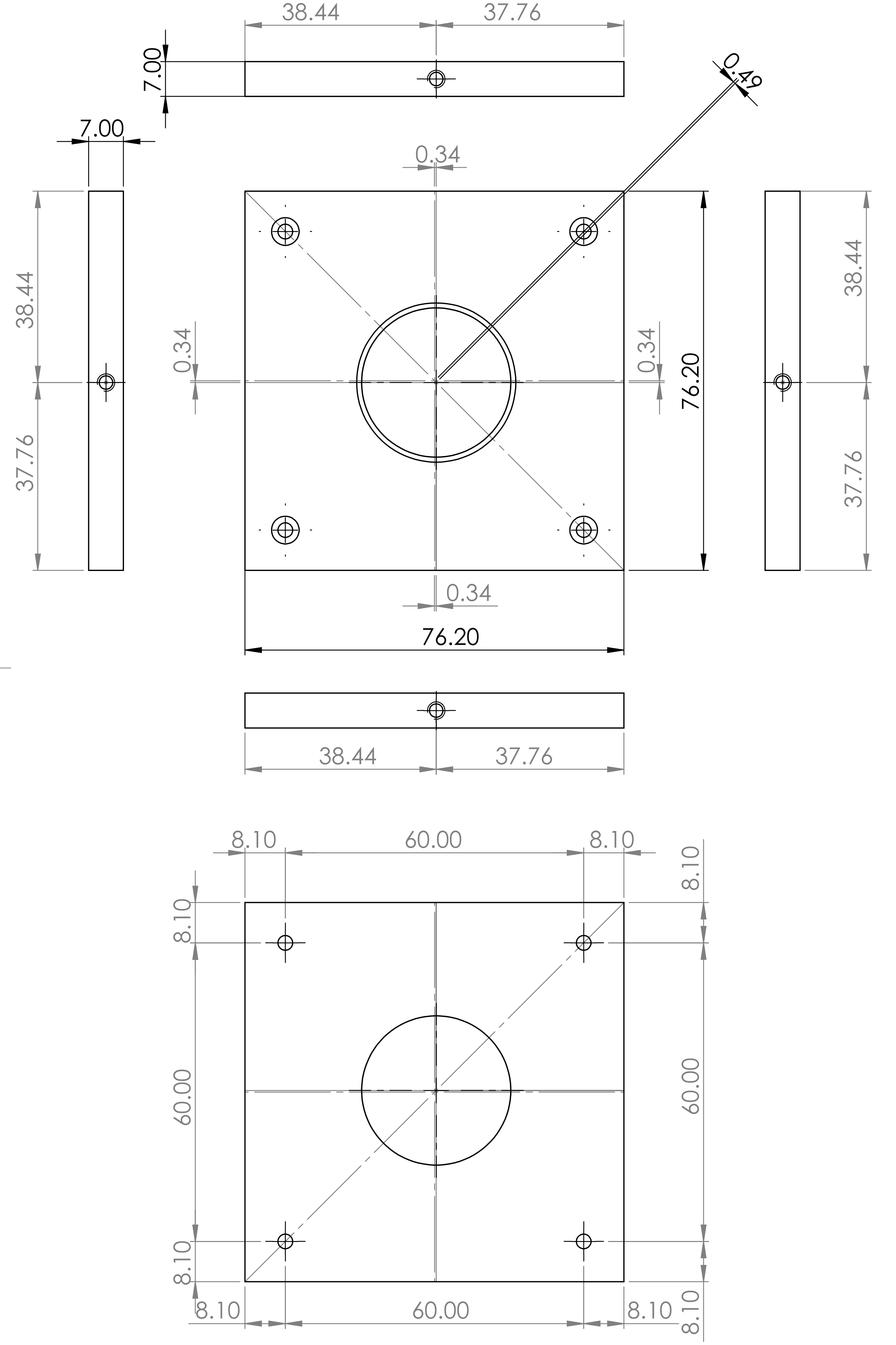


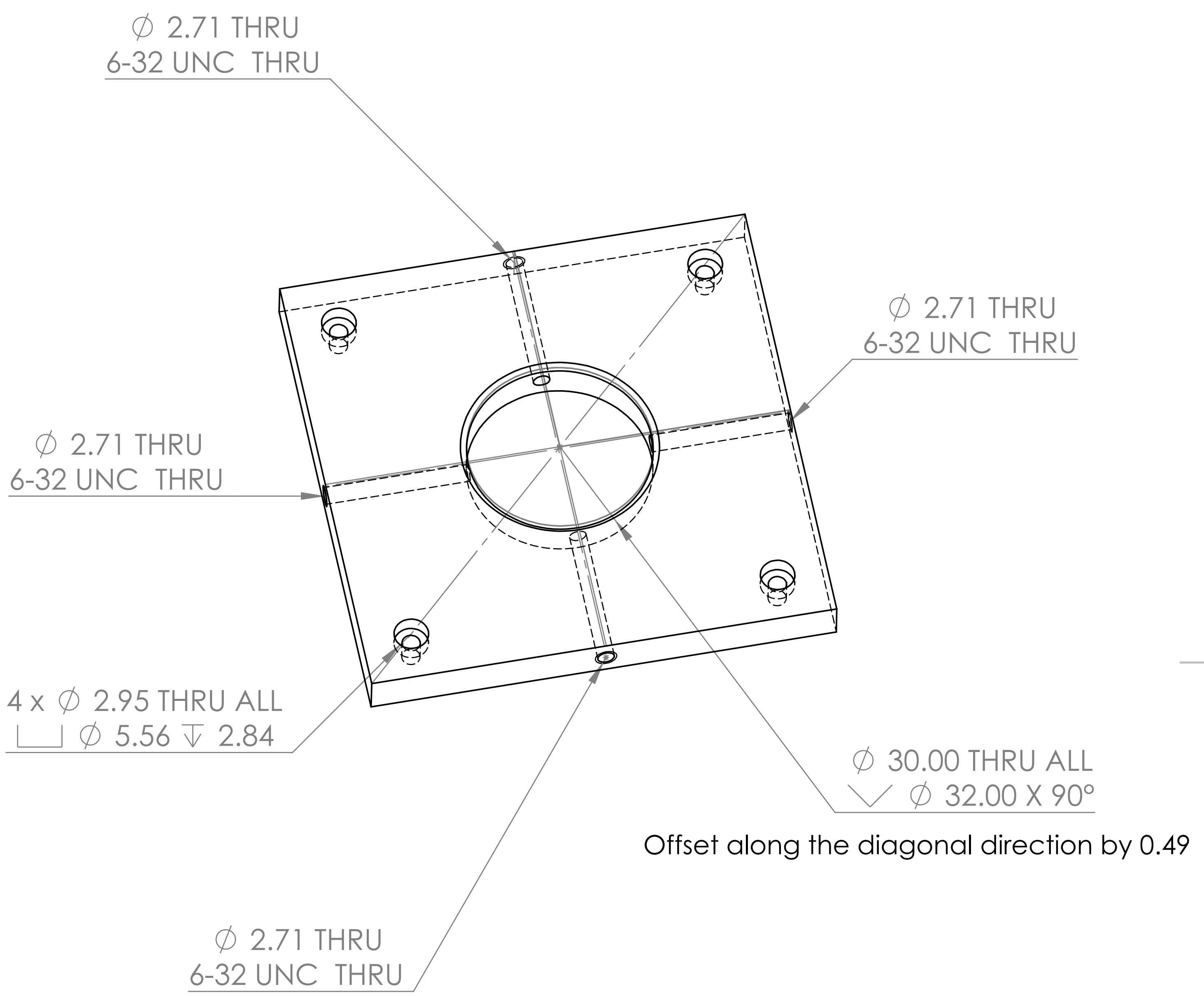


| | | | |
|---|---|---|---|
| UNLESS OTHERWISE SPECIFIED:<br>DIMENSIONS ARE IN MILLIMETERS<br>SURFACE FINISH:<br>TOLERANCES:<br>LINEAR:<br>ANGULAR: | FINISH: | DEBURR AND BREAK SHARP EDGES | DO NOT SCALE DRAWING / REVISION<br>Quantity: 1 |

| | NAME | SIGNATURE | DATE |
|---|---|---|---|
| DRAWN | | | |
| CHK'D | | | |
| APPV'D | | | |
| MFG | | | |
| Q.A | | | |

| | |
|---|---|
| TITLE: | Mounting Plate (galvo) |
| MATERIAL: | Anodized Aluminum |
| DWG NO. | Part Drawing 04 |
| SIZE | A3 |
| WEIGHT: | |
| SCALE:1:1 | SHEET 1 OF 1 |
