## Supplemental data, tables, and software for "Guide to the construction and use of an adaptive optics two-photon microscope with direct wavefront sensing": PD-05 backup Galvo holder (6-mm 8135K Y-galvo).PDF

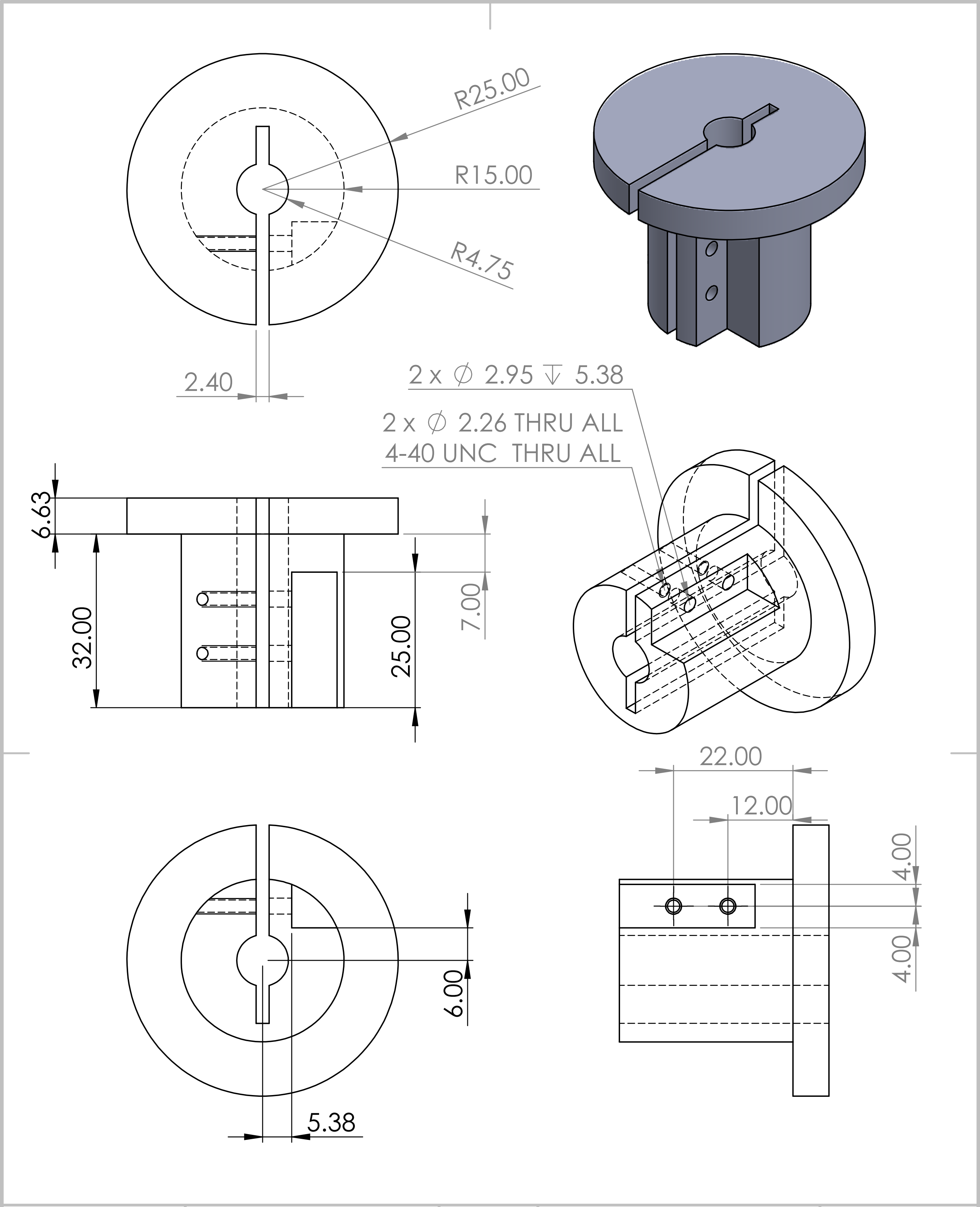


UNLESS OTHERWISE SPECIFIED:
DIMENSIONS ARE IN MILLIMETERS
SURFACE FINISH:
TOLERANCES:
LINEAR:
ANGULAR:

FINISH:

DEBURR AND BREAK SHARP EDGES

DO NOT SCALE DRAWING

REVISION

Quantity: 1

| | NAME | SIGNATURE | DATE |
|---|---|---|---|
| DRAWN | | | |
| CHK'D | | | |
| APPV'D | | | |
| MFG | | | |
| Q.A | | | |

TITLE:

### Galvo Holder
#### (6-mm 8315K Y-galvo)

MATERIAL: Anodized Aluminum

DWG NO. Part Drawing 05 (Backup)

A4

WEIGHT:

SCALE:1:1

SHEET 1 OF 1
