## Supplemental data, tables, and software for "Guide to the construction and use of an adaptive optics two-photon microscope with direct wavefront sensing": PD-06 resonent scanner mounting plate.PDF

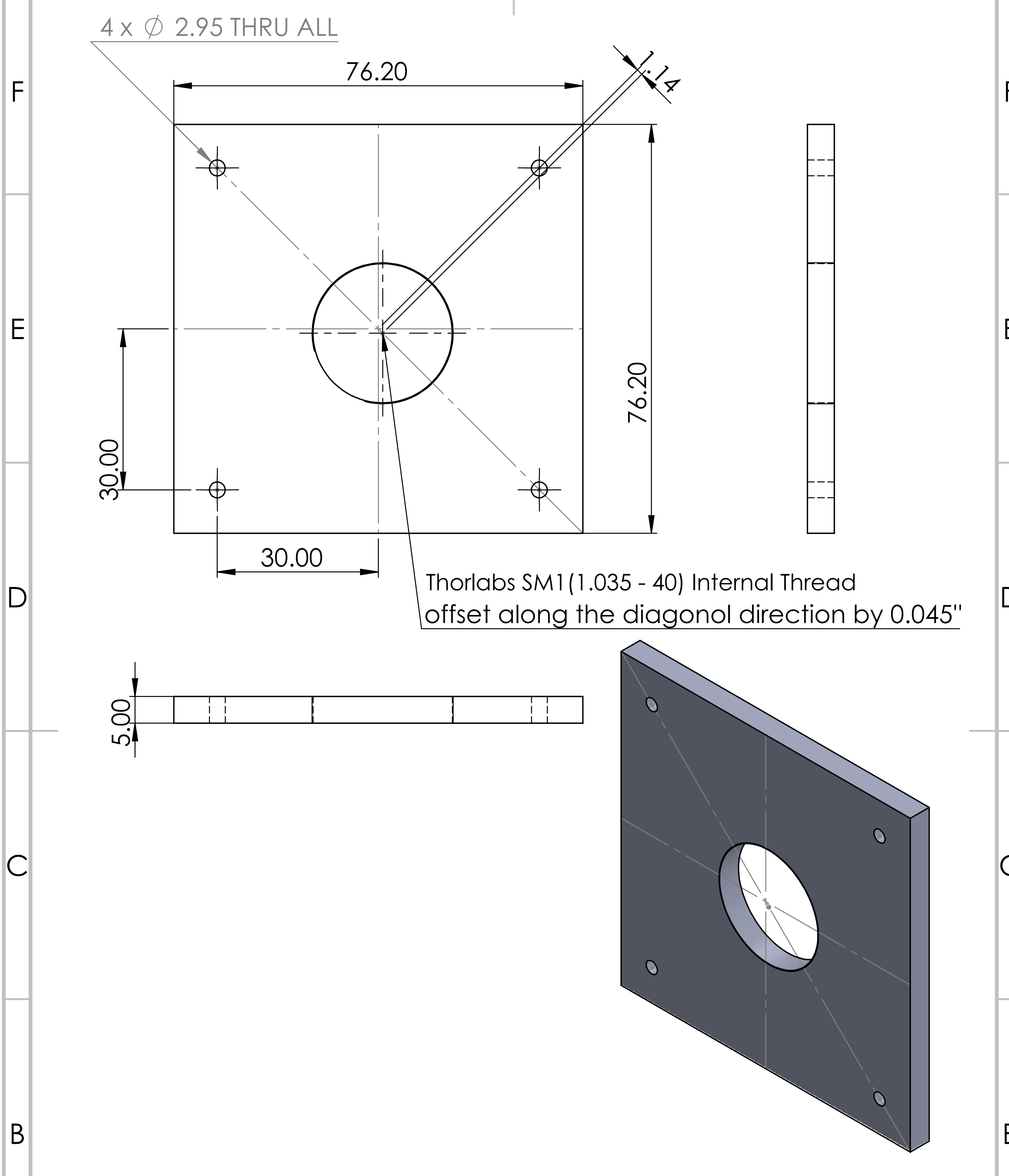

| | | | | |
|---|---|---|---|---|
| UNLESS OTHERWISE SPECIFIED: DIMENSIONS ARE IN MILLIMETERS SURFACE FINISH: TOLERANCES: LINEAR: ANGULAR: | FINISH: | DEBURR AND BREAK SHARP EDGES | DO NOT SCALE DRAWING | REVISION |
| | | | Quantity: 1 | |

| | NAME | SIGNATURE | DATE | |
|---|---|---|---|---|
| DRAWN | | | | |
| CHK'D | | | | |
| APPV'D | | | | |
| MFG | | | | |
| Q.A | | | | MATERIAL: Anodized Aluminum |
| | | | | WEIGHT: |

TITLE:

### Resonent Scanner Mounting Plate

DWG NO. Part Drawing 06 | A4

SCALE:1:1 | SHEET 1 OF 1
