## Supplemental data, tables, and software for "Guide to the construction and use of an adaptive optics two-photon microscope with direct wavefront sensing": PD-08 extra-thick retaining ring.PDF

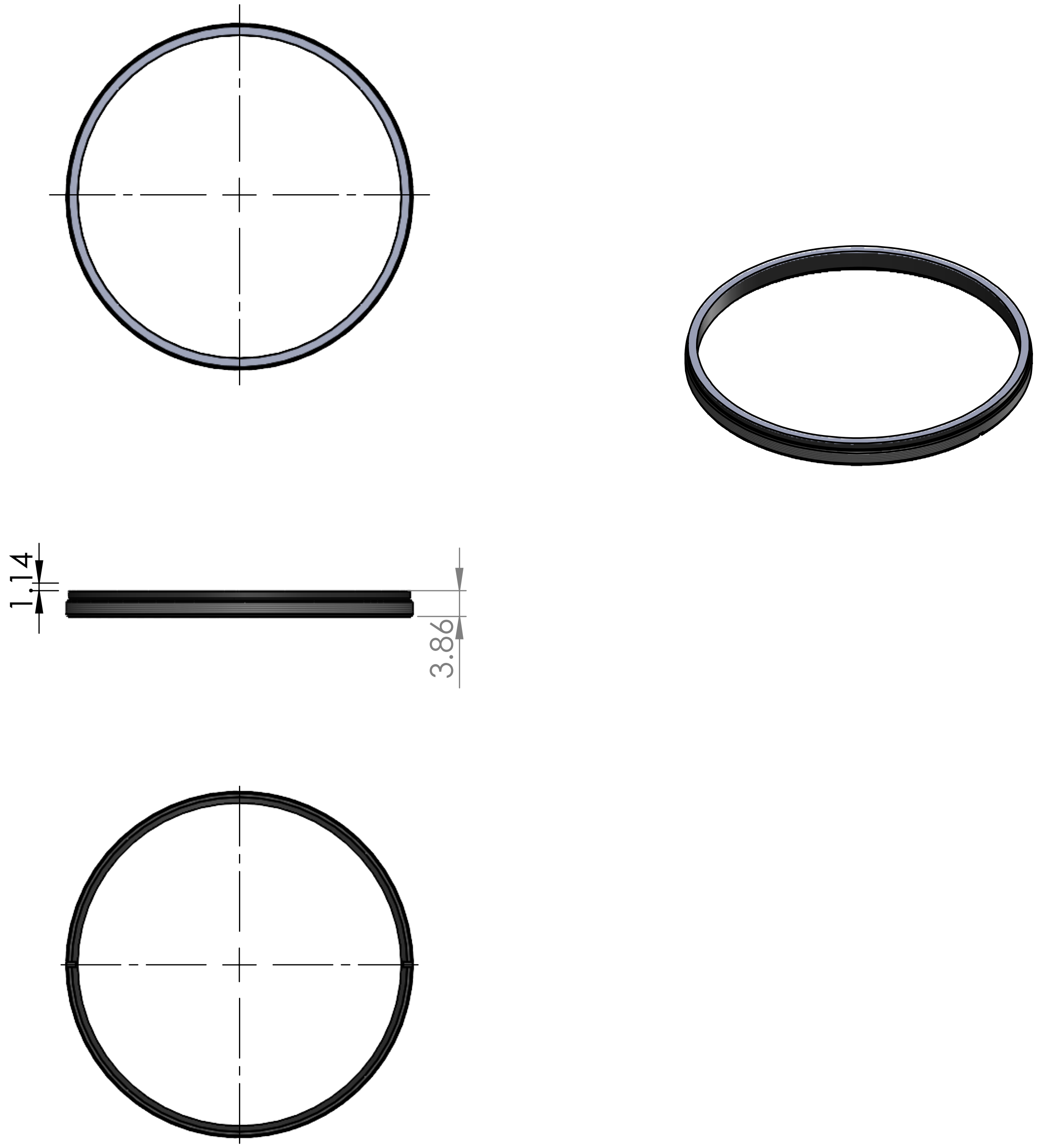


Modified from Thorlabs, SM2RRC
Thickness: 5 → 4

| | | | |
|---|---|---|---|
| UNLESS OTHERWISE SPECIFIED:<br>DIMENSIONS ARE IN MILLIMETERS<br>SURFACE FINISH:<br>TOLERANCES:<br>LINEAR:<br>ANGULAR: | FINISH: | DEBURR AND BREAK SHARP EDGES | DO NOT SCALE DRAWING / REVISION<br>Quantity: 1 |

| | NAME | SIGNATURE | DATE |
|---|---|---|---|
| DRAWN | | | |
| CHK'D | | | |
| APPV'D | | | |
| MFG | | | |
| Q.A | | | |

TITLE: Extra-Thick Retaining Ring (Modified)

MATERIAL: Anodized Aluminum

DWG NO. Part Drawing 08

A4

WEIGHT:

SCALE:1:1

SHEET 1 OF 1
