## Supplemental data, tables, and software for "Guide to the construction and use of an adaptive optics two-photon microscope with direct wavefront sensing": PD-10 ER 11 inch + 6 mm.PDF

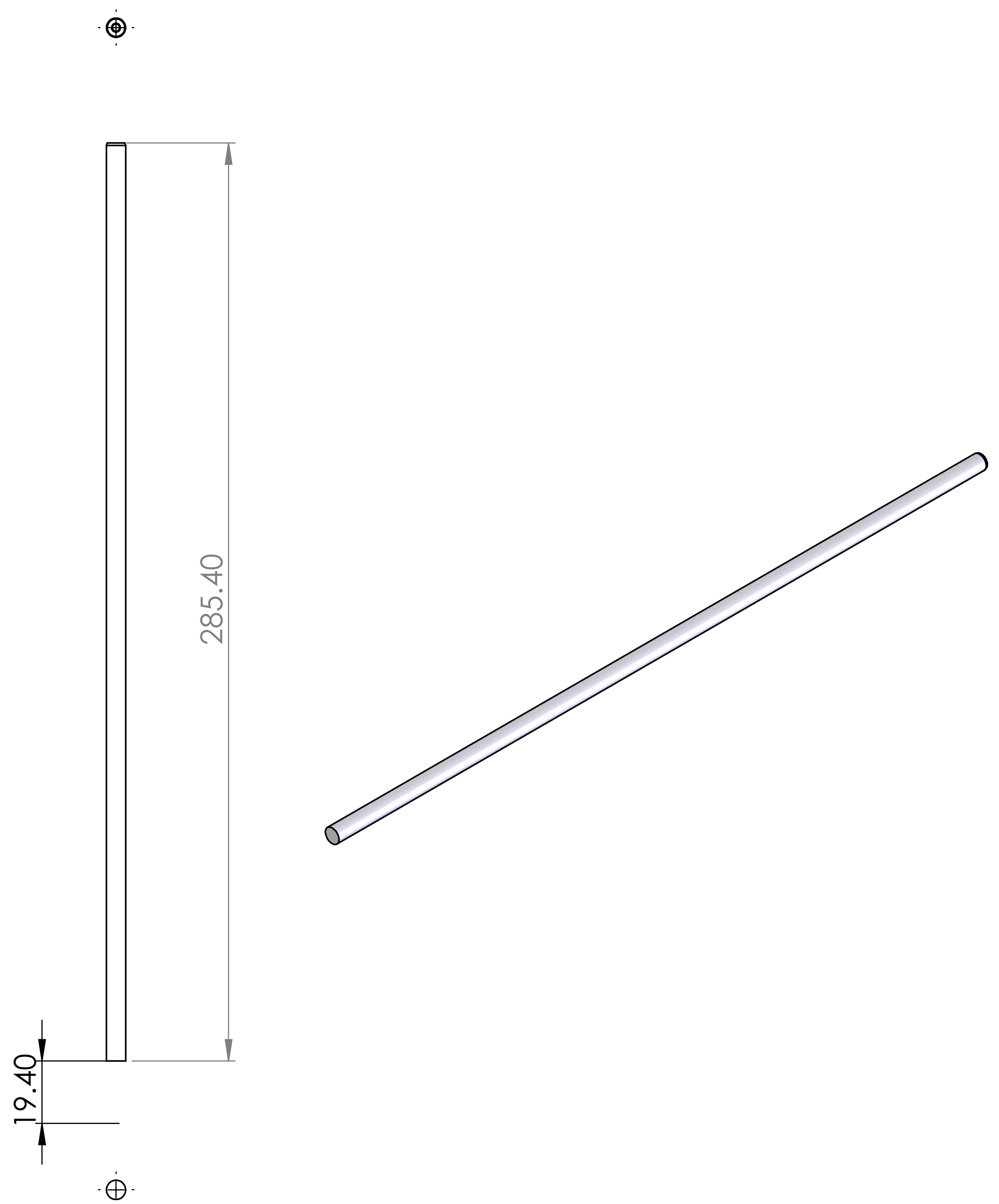

Modified from ER12
Reduce the length to 11" + 6 mm

| | | | |
|---|---|---|---|
| UNLESS OTHERWISE SPECIFIED:<br>DIMENSIONS ARE IN MILLIMETERS<br>SURFACE FINISH:<br>TOLERANCES:<br>LINEAR:<br>ANGULAR: | FINISH: | DEBURR AND BREAK SHARP EDGES | DO NOT SCALE DRAWING / REVISION<br>Quantity: 4 |

| | NAME | SIGNATURE | DATE | |
|---|---|---|---|---|
| DRAWN | | | | |
| CHK'D | | | | |
| APPV'D | | | | |
| MFG | | | | |
| Q.A | | | | MATERIAL: STAINLESS STEEL |
| | | | | WEIGHT: |

TITLE:

### ER11 inch + 6 mm

DWG NO. Part Drawing 10 — A4

SCALE:1:2 — SHEET 1 OF 1
