## Supplemental data, tables, and software for "Guide to the construction and use of an adaptive optics two-photon microscope with direct wavefront sensing": PD-11 objective mounting plate.PDF

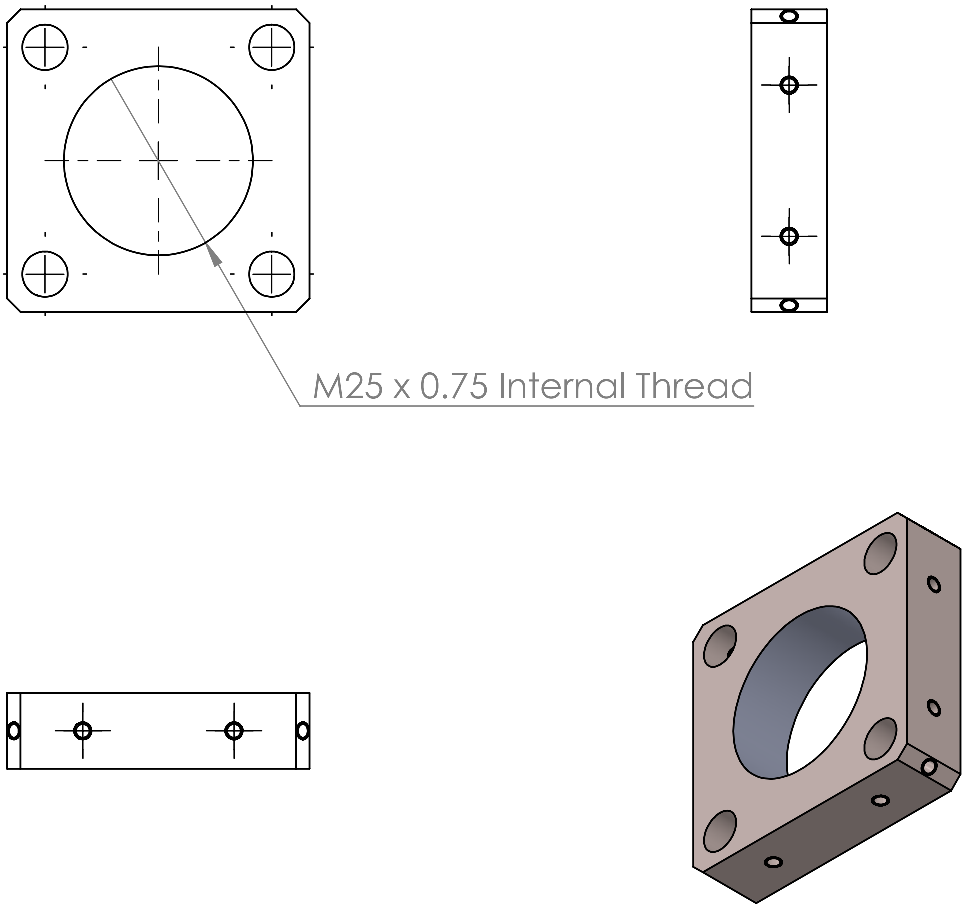


Modified from LINOS Microbench, G061008000
Tapped hole (M25 x 0.75) for the Olympus objective

| | | | |
|---|---|---|---|
| UNLESS OTHERWISE SPECIFIED:<br>DIMENSIONS ARE IN MILLIMETERS<br>SURFACE FINISH:<br>TOLERANCES:<br>LINEAR:<br>ANGULAR: | FINISH: | DEBURR AND BREAK SHARP EDGES | DO NOT SCALE DRAWING — REVISION<br>Quantity: 1 |

| | NAME | SIGNATURE | DATE |
|---|---|---|---|
| DRAWN | | | |
| CHK'D | | | |
| APPV'D | | | |
| MFG | | | |
| Q.A | | | |

TITLE: Objective Mounting Plate

MATERIAL: Anodized Aluminum

DWG NO. Part Drawing 11

A4

WEIGHT:

SCALE:1:1

SHEET 1 OF 1
