## Supplemental data, tables, and software for "Guide to the construction and use of an adaptive optics two-photon microscope with direct wavefront sensing": PD-15 dichroic mount (long).pdf

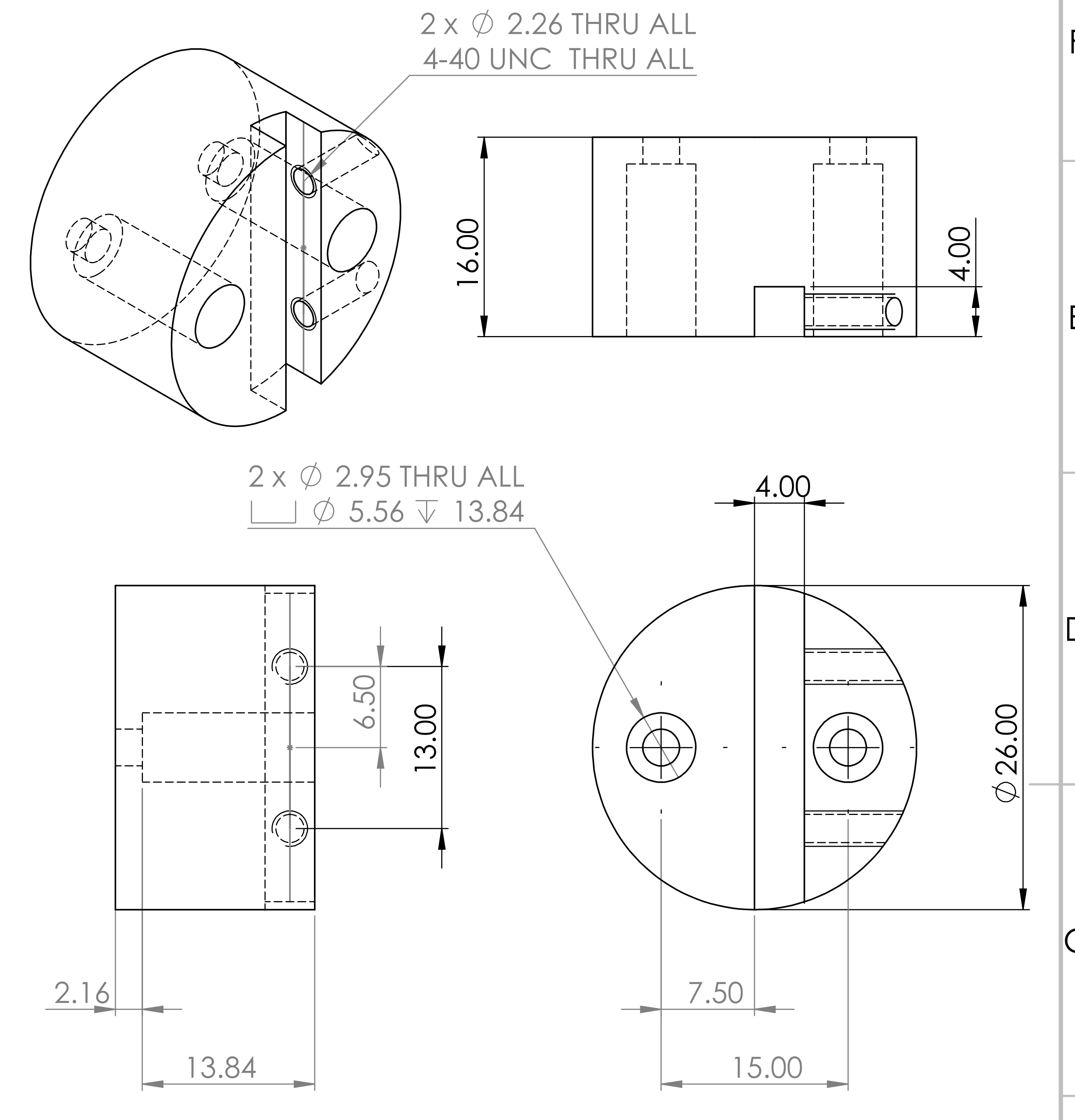


UNLESS OTHERWISE SPECIFIED:
DIMENSIONS ARE IN MILLIMETERS
SURFACE FINISH:
TOLERANCES:
LINEAR:
ANGULAR:

FINISH:

DEBURR AND BREAK SHARP EDGES

DO NOT SCALE DRAWING

REVISION

Quantity: 2

| | NAME | SIGNATURE | DATE |
|---|---|---|---|
| DRAWN | | | |
| CHK'D | | | |
| APPV'D | | | |
| MFG | | | |
| Q.A | | | |

TITLE:

### Dichroic Mount (long)

MATERIAL:
Anodized Aluminum

DWG NO.
Part Drawing 15

A4

WEIGHT:

SCALE:2:1

SHEET 1 OF 1
