## Supplemental data, tables, and software for "Guide to the construction and use of an adaptive optics two-photon microscope with direct wavefront sensing": PD-16 adjustable prism insert.pdf

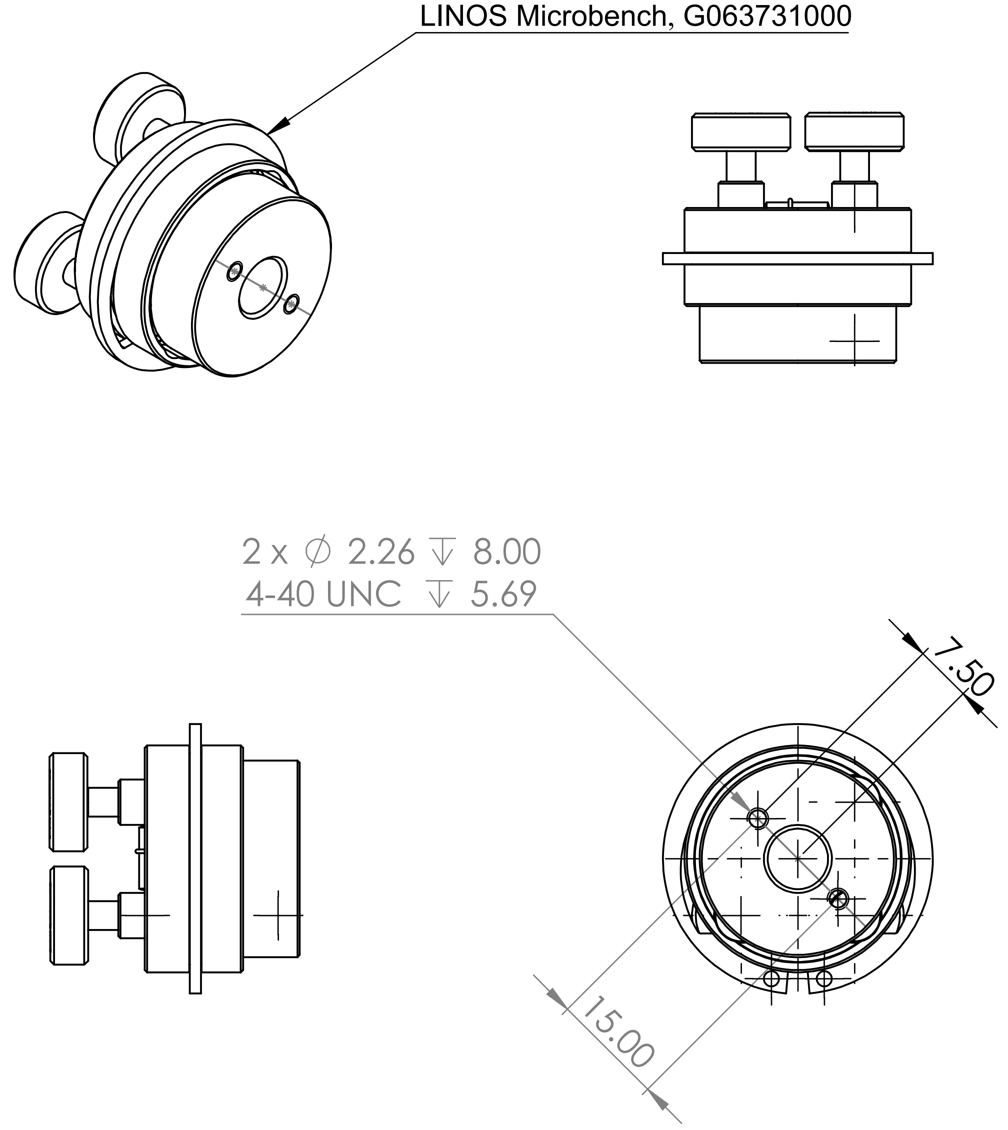


Modified from LINOS Microbench, G063731000
Add 2 tapped holes

| | | | |
|---|---|---|---|
| UNLESS OTHERWISE SPECIFIED: DIMENSIONS ARE IN MILLIMETERS SURFACE FINISH: TOLERANCES: LINEAR: ANGULAR: | FINISH: | DEBURR AND BREAK SHARP EDGES | DO NOT SCALE DRAWING |
| | | | REVISION |
| | | | Quantity: 4 |

| | NAME | SIGNATURE | DATE |
|---|---|---|---|
| DRAWN | | | |
| CHK'D | | | |
| APPV'D | | | |
| MFG | | | |
| Q.A | | | |

TITLE: Adjustable Prism Insert (Modified)

MATERIAL: Anodized Aluminum

DWG NO. Part Drawing 16

A4

WEIGHT:

SCALE:1:1

SHEET 1 OF 1
