## Supplemental data, tables, and software for "Guide to the construction and use of an adaptive optics two-photon microscope with direct wavefront sensing": PD-18 retaining ring (L12 L13 l14).pdf

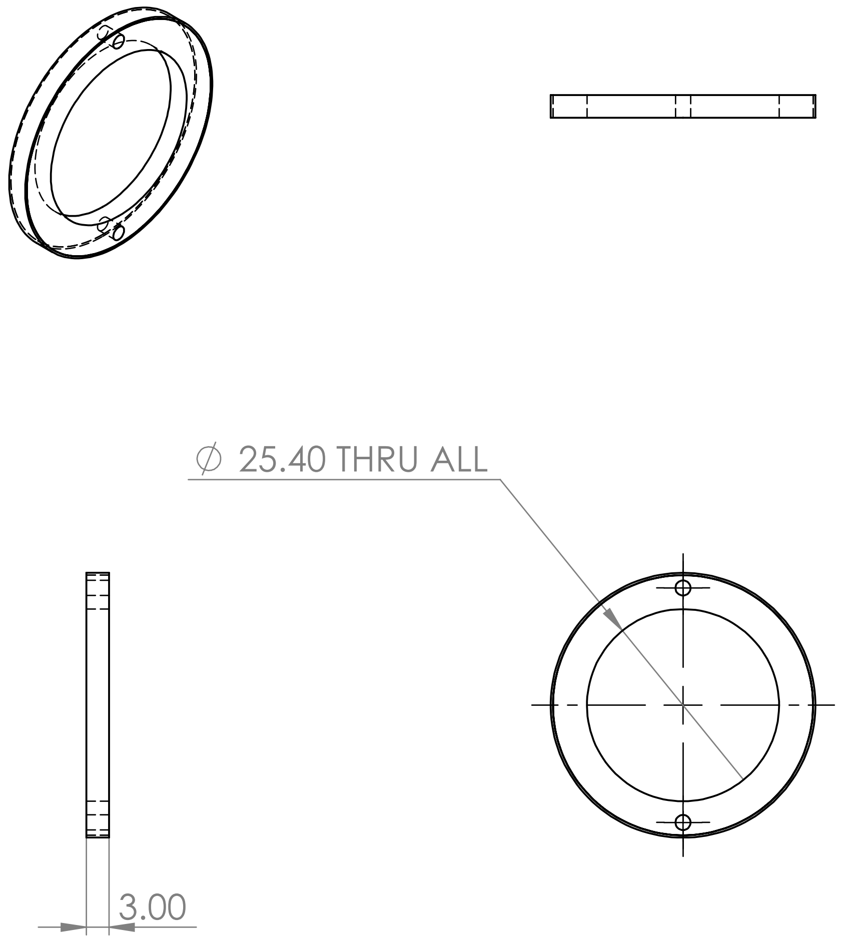


Modified from LINOS Microbench, G065078000
Enlarge center hole

| | | |
|---|---|---|
| UNLESS OTHERWISE SPECIFIED:<br>DIMENSIONS ARE IN MILLIMETERS<br>SURFACE FINISH:<br>TOLERANCES:<br>LINEAR:<br>ANGULAR: | FINISH: | DEBURR AND BREAK SHARP EDGES |

DO NOT SCALE DRAWING | REVISION

Quantity: 3

| | NAME | SIGNATURE | DATE |
|---|---|---|---|
| DRAWN | | | |
| CHK'D | | | |
| APPV'D | | | |
| MFG | | | |
| Q.A | | | |

TITLE:

### Retaining Ring (for L12 ~ L14)

MATERIAL:
Anodized Aluminum

DWG NO.
Part Drawing 18

A4

WEIGHT:

SCALE:1:1

SHEET 1 OF 1
