## Supplemental data, tables, and software for "Guide to the construction and use of an adaptive optics two-photon microscope with direct wavefront sensing": PD-21 deformable mirror breadboard.PDF

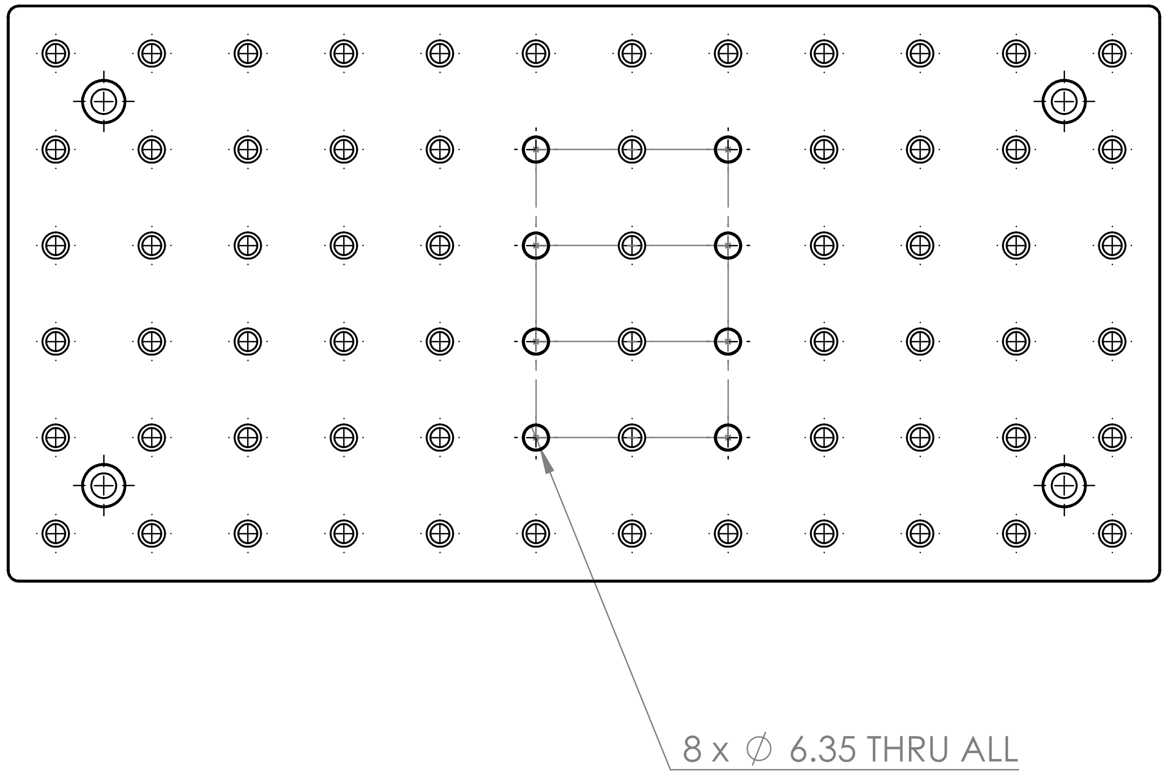


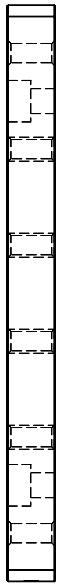

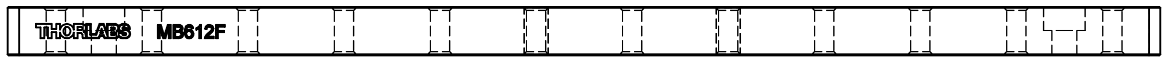


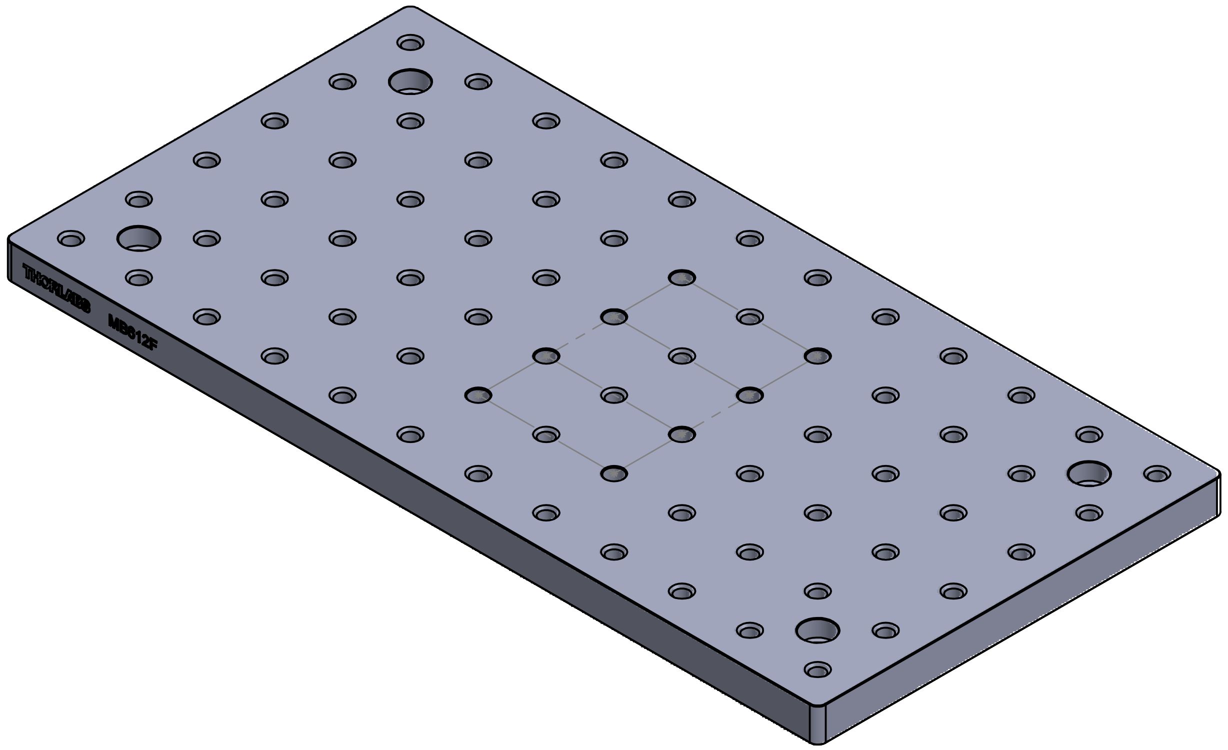

Modified from Thorlabs, MB612F
Remove the threads for 8 tapped holes

| | | | | |
|---|---|---|---|---|
| UNLESS OTHERWISE SPECIFIED: DIMENSIONS ARE IN MILLIMETERS SURFACE FINISH: TOLERANCES: LINEAR: ANGULAR: | FINISH: | DEBURR AND BREAK SHARP EDGES | DO NOT SCALE DRAWING | REVISION |
| | | | Quantity: 1 | |

| | NAME | SIGNATURE | DATE |
|---|---|---|---|
| DRAWN | | | |
| CHK'D | | | |
| APPV'D | | | |
| MFG | | | |
| Q.A | | | |

TITLE: Deformable Mirror Breadboard

MATERIAL: Anodized Aluminum

DWG NO. Part Drawing 21

A3

WEIGHT:

SCALE:1:2

SHEET 1 OF 1
