## Supplemental data, tables, and software for "Guide to the construction and use of an adaptive optics two-photon microscope with direct wavefront sensing": PD-22 EMCCD breadboard.pdf

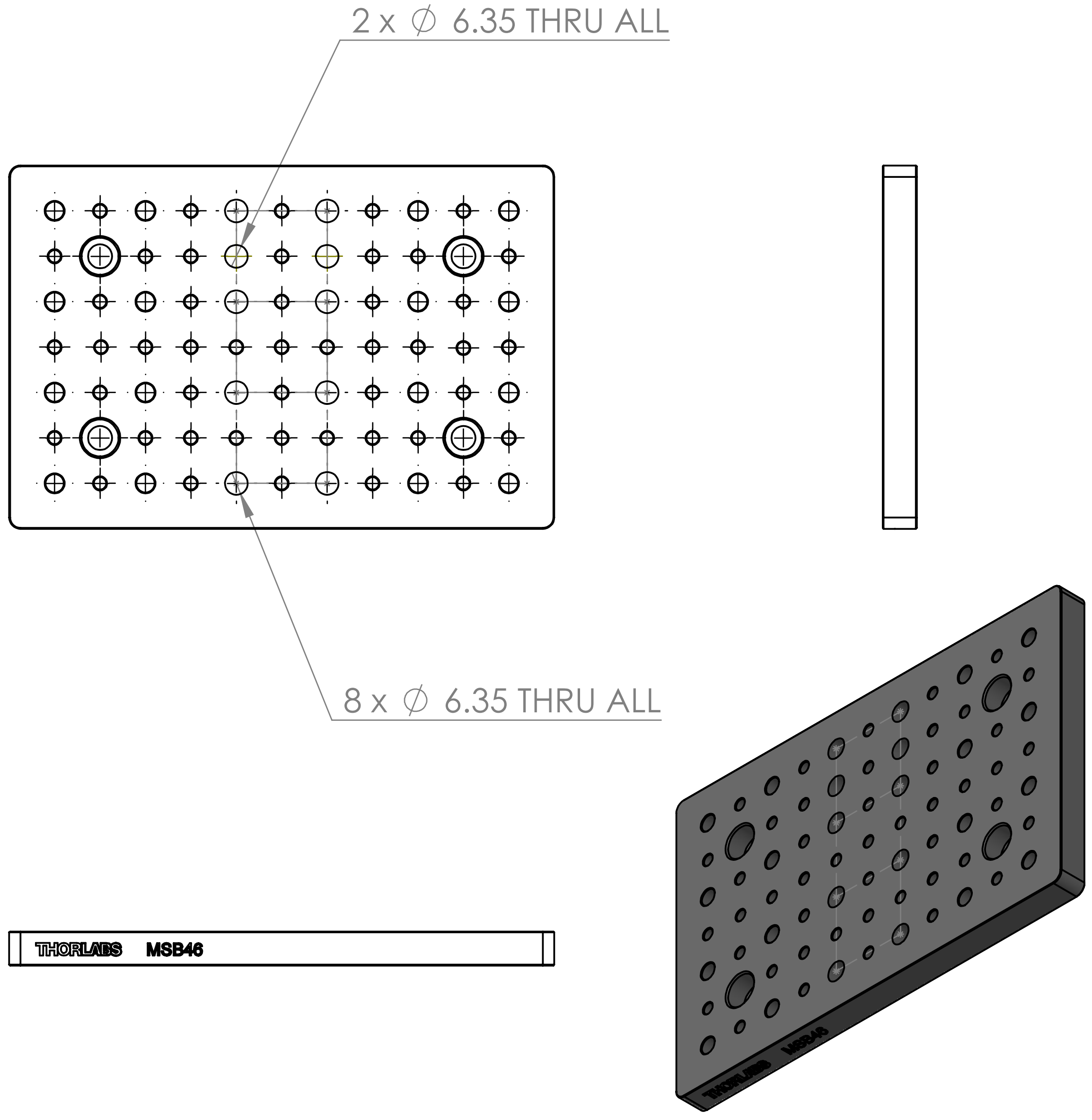


Modified from Thorlabs, MSB46
Remove the threads for 8 tapped holes

| | | | | |
|---|---|---|---|---|
| UNLESS OTHERWISE SPECIFIED: DIMENSIONS ARE IN MILLIMETERS SURFACE FINISH: TOLERANCES: LINEAR: ANGULAR: | FINISH: | DEBURR AND BREAK SHARP EDGES | DO NOT SCALE DRAWING | REVISION |
| | | | Quantity: 1 | |

| | NAME | SIGNATURE | DATE | |
|---|---|---|---|---|
| DRAWN | | | | TITLE: EMCCD Breadboard |
| CHK'D | | | | |
| APPV'D | | | | |
| MFG | | | | |
| Q.A | | | MATERIAL: Anodized Aluminum | DWG NO. Part Drawing 22 — A4 |
| | | | WEIGHT: | SCALE:1:5 — SHEET 1 OF 1 |
