## Supplemental data, tables, and software for "Guide to the construction and use of an adaptive optics two-photon microscope with direct wavefront sensing": PD-24 deformable mirror conjugation tool.PDF

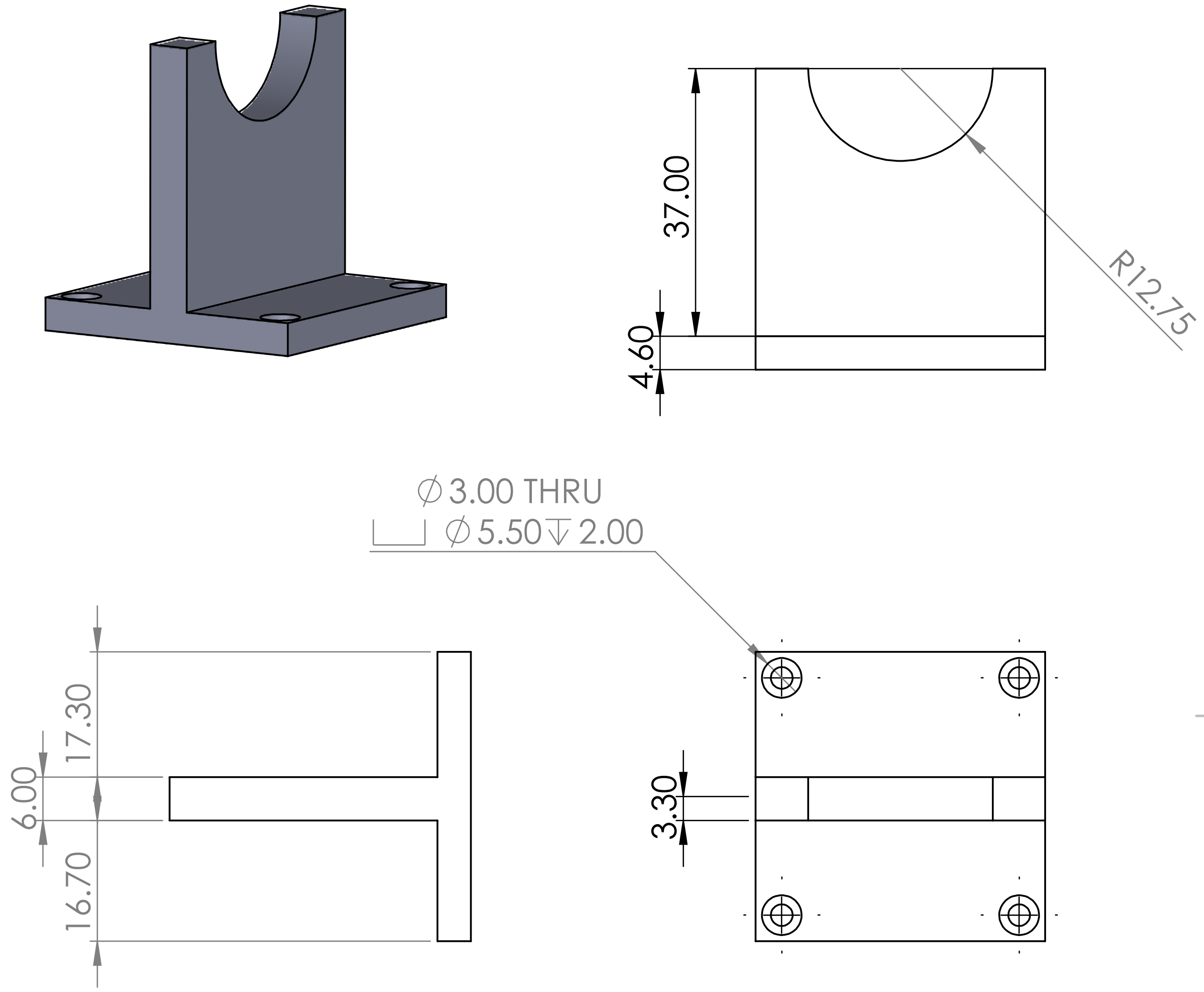


UNLESS OTHERWISE SPECIFIED:
DIMENSIONS ARE IN MILLIMETERS
SURFACE FINISH:
TOLERANCES:
LINEAR:
ANGULAR:

FINISH:

DEBURR AND BREAK SHARP EDGES

DO NOT SCALE DRAWING

REVISION

Quantity: 1

| | NAME | SIGNATURE | DATE |
|---|---|---|---|
| DRAWN | | | |
| CHK'D | | | |
| APPV'D | | | |
| MFG | | | |
| Q.A | | | |

TITLE:
### Deformable Mirror Conjugation Tool

MATERIAL:
Anodized Aluminum

DWG NO.
Part Drawing 24

A4

WEIGHT:

SCALE:1:1

SHEET 1 OF 1
